# Integration of Bioinformatics and Machine Learning to characterize *Fusobacterium nucleatum*’s pathogenicity

**DOI:** 10.1101/2025.08.21.671586

**Authors:** Zihan Tian, Pietro Liò

## Abstract

*Fusobacterium nucleatum* has been found to be associated with cancer lesions in both oral and colon cancers. Although important studies have dissected the clinical aspects of its remarkable pathogenicity, there is a lack of molecular studies. This study aimed to computationally predict potential pathogenicity islands in *F. nucleatum* ATCC 25586, with candidate functional relationships among PAI-encoded proteins, and generate testable hypotheses regarding iron-dependent virulence mechanisms. We employed an integrative bioinformatics pipeline combining genomic island prediction (IslandViewer), promoter analysis (PePPER), codon adaptation index calculation, protein interaction prediction (STRING), co-expression network inference (bnlearn), structural modeling (AlphaFold-Multimer), and genome-scale metabolic modeling (CarveMe/COBRApy). Computational predication was integrated with published literature to formulate mechanistic hypotheses. Our analysis identified three candidate genomic islands, with Region 2 (PAI2; 1,496,613 to 1,523,855 bp) exhibiting the characteristics most consistent with a functional pathogenicity island, including predicted mobile genetic elements, putative toxin-antitoxin systems, and strong promoter motifs. We propose a mechanistic hypothesis linking hemolysin-mediated iron acquisition to cancer promotion through oxidative stress and Hippo pathway modulation. Our work has two immediate and important benefits: the improved understanding of the biological processes that shape the pathogenicity and evolution of *Fusobacterium nucleatum* at the molecular level and the improved ability to integrate and automate the state-of-the-art bioinformatics tools and machine learning approaches in the inference of the mechanistic interpretability of a pathogenic phenotype.

## Introduction

Various studies indicate that microbial dysbiosis is associated with oral and systemic diseases (Han, 2015). *Fusobacterium nucleatum* (*F.nucleatum*), a Gram-negative anaerobe naturally found in the oral cavity of humans, contributes to periodontal disease, as well as oral and colorectal cancers (Patel et al., 2022; Queen et al., 2025). The oral cavity is full of circulating saliva that contains a high level of dissolved oxygen, and oral microbes, such as *F. nucleatum*, are subject to oxidative stress in the form of reactive oxygen species. An adaptive response that allows *F. nucleatum* to withstand oxidative stress enables the bacterium to play a prominent role in shaping the oral biofilm (McGregor and Wolthers, 2025). The association of *F. nucleatum* with oral squamous cell carcinoma (OSCC) highlights its role via oncogenic signaling pathways and has metabolic adaptability in the tumor microenvironments (Lim et al., 2025).

Beside being a pathogen, *F. nucleatum* could induce resistance to antitumor therapies, tumor recurrence, and be associated with a poor prognosis in patients. Chemotherapy resistance is one of the major contributors to the poor prognosis in colorectal cancer. Mechanistic studies have shown that F. nucleatum can inhibit pyroptosis, a form of programmed cell death induced by chemotherapeutic agents, by down-regulating the Hippo signaling pathway and up-regulating antiapoptotic BCL2 (Han et al., 2020; Wang et al., 2024). In addition, growing evidence suggests that removal of tumor-associated *F. nucleatum* can induce the host immune response to recognize tumor cells, suggesting potential future antibacteria / tumor vaccine applications (Chen and Huang, 2024).

*F.nucleatum* can coexist with more than 500 other species, and its elongated shape allows it to bridge other bacteria, maintaining the development of biofilm and facilitating the spreading of periodontal disease in humans. *F. nucleatum* enhances the invasive potential of other microorganisms, such as *Porphyromonas gingivalis*. Together, these bacteria form a pathogenic alliance that can significantly alter host responses and promote cancer progression (How et al., 2016; Mohammed et al., 2020; Patel et al., 2022).

Despite *F. nucleatum*’s association with various cancers in many different tissues and its contribution to therapeutic resistance, the genomic, proteomic, and metabolic characteristics of its specific pathogenic function remain poorly characterized (Ma et al., 2023). To address this gap, this study focuses on a specific strain of *F. nucleatum* ATCC 25586 that is naturally present in the human oral cavity. An integrated bioinformatics and machine learning framework is used to predict potential pathogenicity islands (PAIs). Protein and metabolic pathways that may be associated with cancer are identified. Structural predictions (AlphaFold, TMHMM, and String) are used to evaluate protein-protein interactions and potential transmembrane regions. In parallel, genome-scale metabolic modeling using COBRApy and flux balance analysis was used to explore iron dependency and pH tolerance. This study aims to better understand what contributes to *F. nucleatum* pathogenicity and offers insights into its connection to cancer progression and therapeutic resistance.

Notably, the term *F. nucleatum* can be defined in different ways, often referring to a subspecies group (*F. nucleatum subsp. animalis*, *F. nucleatum subsp. nucleatum*, *F. nucleatum subsp. polymorphum*, *F. nucleatum subsp. vincentii*) or specific lineages. In this study, all *F. nucleatum* used in computational results refers to *F. nucleatum subsp. nucleatum* (ATCC 25586, GenBank: AE009951.2), while the term is used in a broader way when referring to literature analysis (Zepeda-Rivera et al., 2025).

We emphasize that our findings are computational predictions that generate testable hypotheses for future experimental validation.

## Methodology: Building a Bioinformatics-Machine Learning integrated workflow

In this section we provide a brief description of the methods and software we have used. The analysis focused on *Fusobacterium nucleatum* subsp. nucleatum strain ATCC 25586, a critical periodontal pathogen with established links to various oral diseases and colorectal cancers.

The complete genome sequence of *Fusobacterium nucleatum* subsp. nucleatum strain ATCC 25586 (GenBank accession: AE009951.2) was retrieved from the NCBI Genome database. Genome annotation was performed using Prokka v1.14 (Seemann, 2014) to generate a detailed catalog of its genetic components, including coding sequences, rRNAs, and tRNAs. Additional functional annotation was performed using eggNOG-mapper v2.0 (Huerta-Cepas et al., 2018).

To identify potential pathogenic islands (PAI) in bacteria of interest, GC skew analysis, homology-based functional analysis (UniProt + BLAST), and IslandViewer (Bertelli et al., 2017) genomic island predictions were used.

Promoter predictions were generated using PePPER (Prokaryote Promoter Prediction) v2.0 (de Jong et al., 2012) and integrated with gene annotations from NCBI GenBank to evaluate open reading frames.

Synonymous codon usage bias was assessed using the Codon Adaptation Index (CAI). The ribosomal protein genes served as the reference set, and the CAI is calculated based on the API Reference (Bagnoli and Lìo, 1995; Lee, 2019). Genes were classified as high-CAI (top 20%, CAI *≥* 0.771) or low-CAI (bottom 20%, CAI *≤* 0.670) based on the genome-wide distribution. Promoter motif analysis was performed using WebLogo Crooks et al. (2004) on 8-mer sequences extracted from promoter regions. BLASTp (Altschul et al., 1990) is used to search protein sequences against the UniProt database.

Homologous proteins are identified and assigned corresponding functional annotations. For virulence factor prediction, we retrieved functional annotation from DAVID, a tool that integrates multiple databases, including KEGG pathway maps, to identify enriched biological themes and map genes to metabolic pathways. Biomni, a machine-learning tool, was applied to cross-reference existing literature and the output of DAVID to label potential pathogenic genes.

Protein-protein interactions were predicted using the STRING database v12.0 (Szklarczyk et al., 2024) with a minimum confidence score threshold of 0.4. Network topology was analyzed using NetworkX to calculate clustering coefficients and network density. Community detection was performed to identify functional modules within the interaction network.

Transmembrane domain predictions were generated using both TMHMM v2.0 (Krogh et al., 2001) and DeepTMHMM (Hallgren et al., 2022) to identify membrane-associated proteins. For selected protein pairs with predicted interactions, AlphaFold-Multimer was used to model complex structures (Jumper et al., 2021).

Expression data for *F. nucleatum* subsp. nucleatum ATCC 25586 was obtained from GEO dataset GSE161360 (Ponath et al., 2021), comprising 18 samples across three growth phases (exponential, mid-log, and stationary) and two treatment conditions. Bayesian network structure learning was performed using the bnlearn R package with the hill-climbing algorithm and BIC scoring. Network edges were validated using 500 bootstrap iterations, with edges retained if bootstrap strength *>* 0.5 and direction probability *>* 0.5. Co-expression relationships were compared with STRING-predicted interactions to identify consensus functional associations.

We constructed a genome-scale metabolic model (GEM) from the Prokka-annotated genome using CarveMe (Machado et al., 2018). The final model included 1,914 reactions, 1,339 metabolites, and 582 gene-protein-reaction associations. We then performed Flux Balance Analysis (FBA) with COBRApy (Ebrahim et al., 2013) to simulate growth across a range of nutrient conditions, specifically exploring iron dependency (70 iron-related reactions, 40 metabolites). We built the proposed Hemolysin-Iron-ROS-Hippo pathway by combining our computational predictions with published experimental data. We systematically adjusted iron uptake constraints to characterize how iron availability influences predicted growth rate. We also carried out essential nutrient analysis by setting individual exchange reaction bounds to zero and quantifying the corresponding changes in biomass flux.

We built the proposed Hemolysin-Iron-ROS-Hippo pathway by combining our computational predictions with published experimental data. Key components of the pathway were identified through genome annotation, and their functional interactions were verified against peer-reviewed studies focused on F. nucleatum pathogenicity and its association with cancer.

## Results and Discussion

### Genome Analysis

As shown in Figure 2, notable fluctuations in GC skew were observed between the 1.6 Mb and 1.9 Mb region, suggesting atypical nucleotide composition that may indicate genomic anomalies such as prophages or integrated mobile genetic elements (MGEs). In addition, genomic island predictions were performed using IslandViewer Bertelli et al. (2017), a computational tool that integrates mul-tiple prediction methods, including IslandPick, IslandPath-DIMOB, and SIGI-HMM. Three regions (366,847-372,320, 1,496,613-1,523,855, 1,967,193-1,999,047) that have the potential to be PAI have been identified.

**Figure 1:**
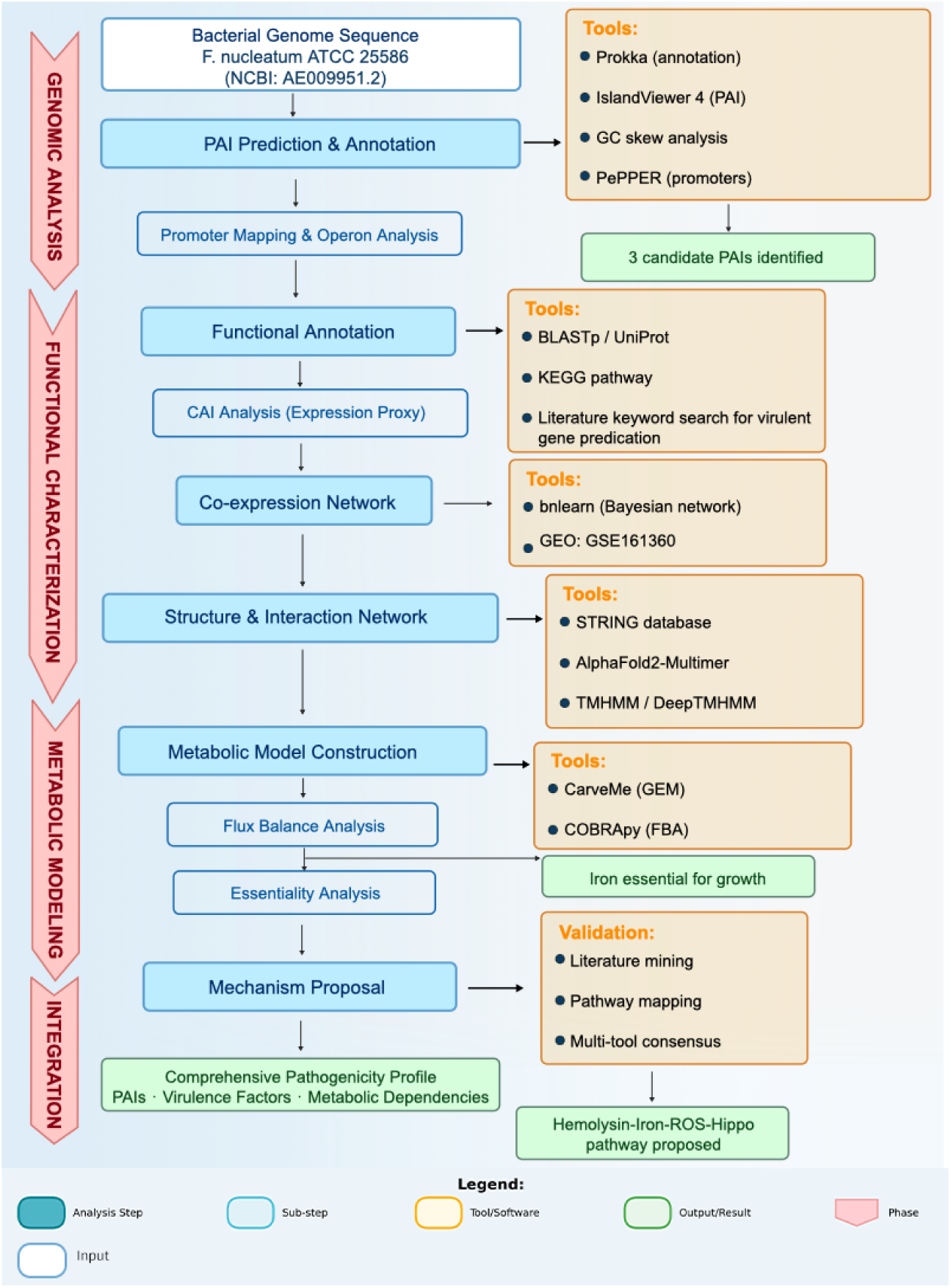
Integrated bioinformatics pipeline for analyzing bacterial pathogenicity. Overview of the computational workflow of four phases: 1) Genomic Analysis, including genome annotation with Prokka, GC skew analysis, and PAI prediction; 2) Functional characterization, covering homology searches, CAI analysis, co-expression network construction with bnlearn, and protein–protein interaction prediction via the STRING database; 3) Metabolic Modeling, including genome-scale model construction with CarveMe and flux balance analysis using COBRApy; and 4) Combination of computational predictions with existing literature evidence to propose a mechanistic hypothesis. Color coding: White boxes are the input. Blue boxes indicate main analysis steps; light blue boxes are sub-steps; yellow boxes indicate tools and software used; green boxes indicate outputs.

**Figure 2:**
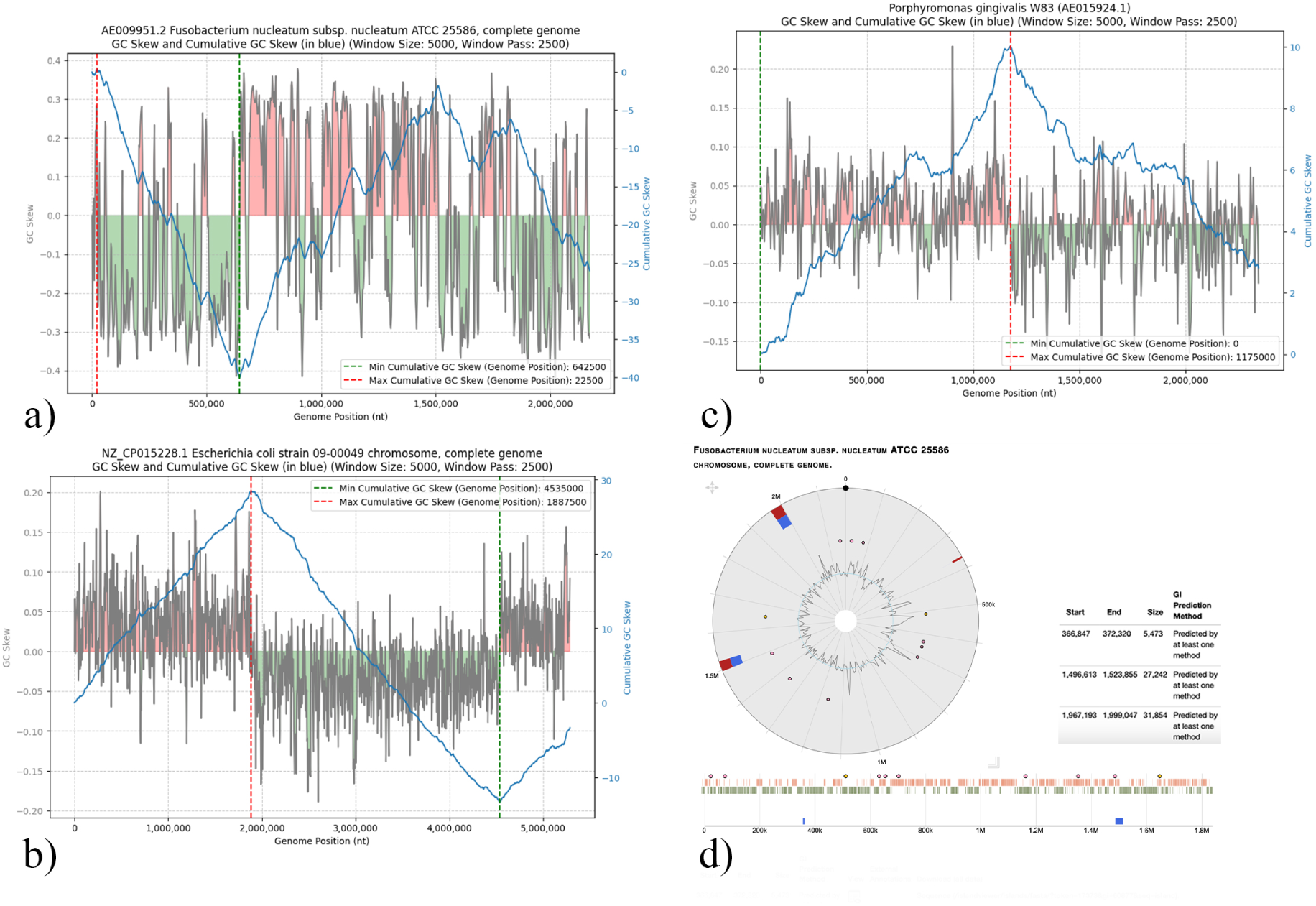
Panels a–c show GC Skew (in grey, where positive GC skew values have a red pattern and negative ones have a green one) and Cumulative GC Skew (blue) analyses of the complete genomes of Fusobacterium nucleatum subsp. nucleatum ATCC 25586 (GenBank: AE009951.2), Escherichia coli strain 09-00049 (GenBank: NZ CP015228.1), and Porphyromonas gingivalis W83 (GenBank: AE015924.1), respectively. These plots were generated using the GC skew-plot tool by Sergi Vallvé and highlight nucleotide composition asymmetries across each genome (Vallvé, 2023). Panel d presents genomic island predictions in F. nucleatum ATCC 25586 using IslandViewer, identifying three potential pathogenicity islands (PAIs) in Fusobacterium nucleatum subsp. nucleatum ATCC 25586. These regions were predicted by at least one of the integrated methods: IslandPick, IslandPath-DIMOB, or SIGI-HMM (Bertelli et al., 2017).

**Figure 3:**
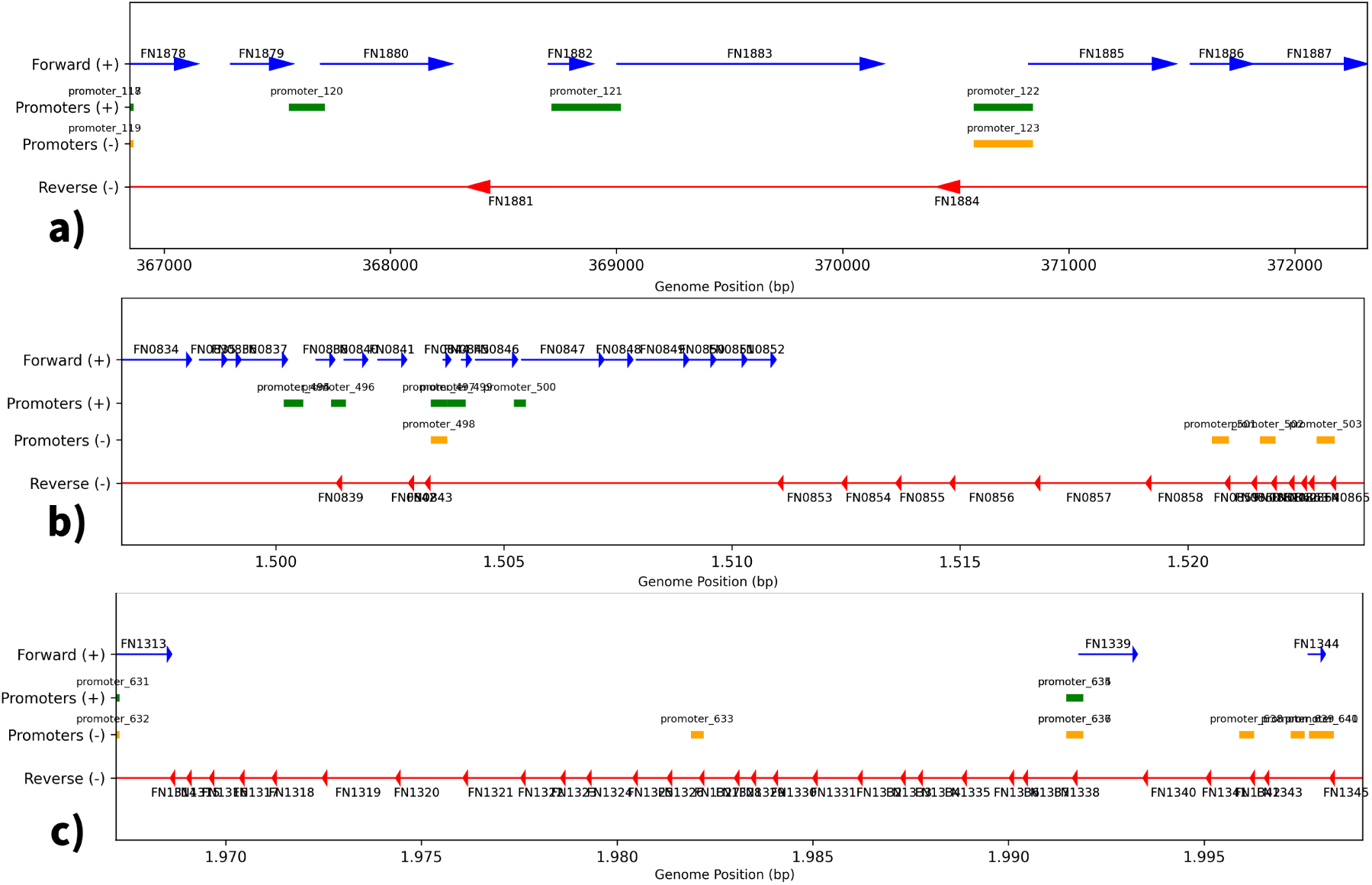
Promoter mapping of 3 predicts potential Pathogenicity Islands (PAIs), showing the dis-tribution of genes on both strands and ORF. Genes are represented by arrows, with blue arrows indicating forward strand (+) genes and red arrows indicating reverse strand (*−*) genes. Promoters are shown as bars, with green bars representing predicted promoters on the forward strand and yellow bars representing predicted promoters on the reverse strand. a) PAI 1 (region 366847–372320) includes 7 predicted promoters (promoter 117 to promoter 123) and 10 genes (locus tags FN1878 to FN1887). b) PAI 2(region 1496613–1523855) with 10 predicted promoters (promoter 494 to promoter 503) and 32 genes (FN0834 to FN0865). c) PAI 3(region 1967193–1999047) contain 11 predicted promoters (promoter 631 to promoter 641) and 33 genes (FN1313 to FN1345).

**Figure 4:**
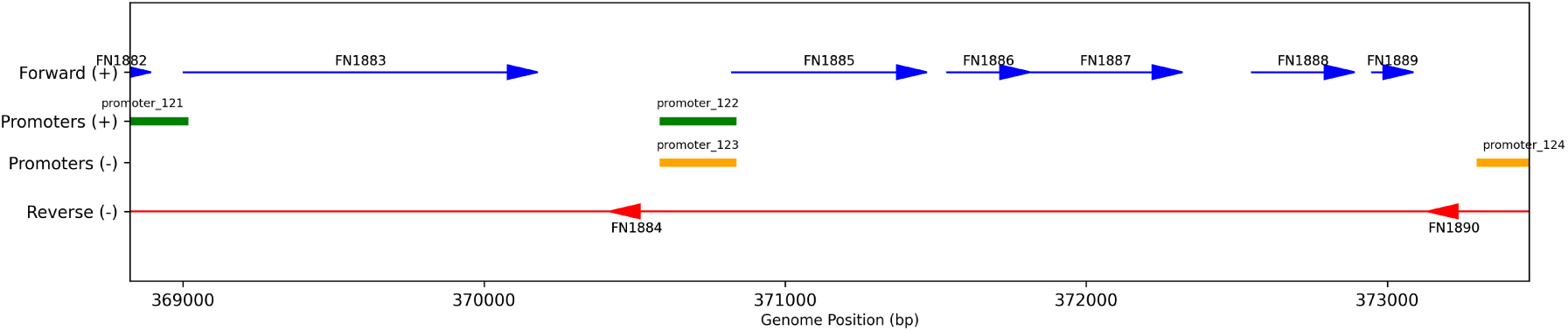
Genomic organization of the region spanning 368,823–373,471 bp surrounding locus tag FN1885. Blue arrows show coding sequences (ORFs) on the forward strand (+), while red arrows show on the reverse strand (*−*). Predicted promoters are depicted as colored bars. Green forward, yellow reverse. Promoter positions are plotted relative to their genomic coordinates and strand orientation.

**Figure 5:**
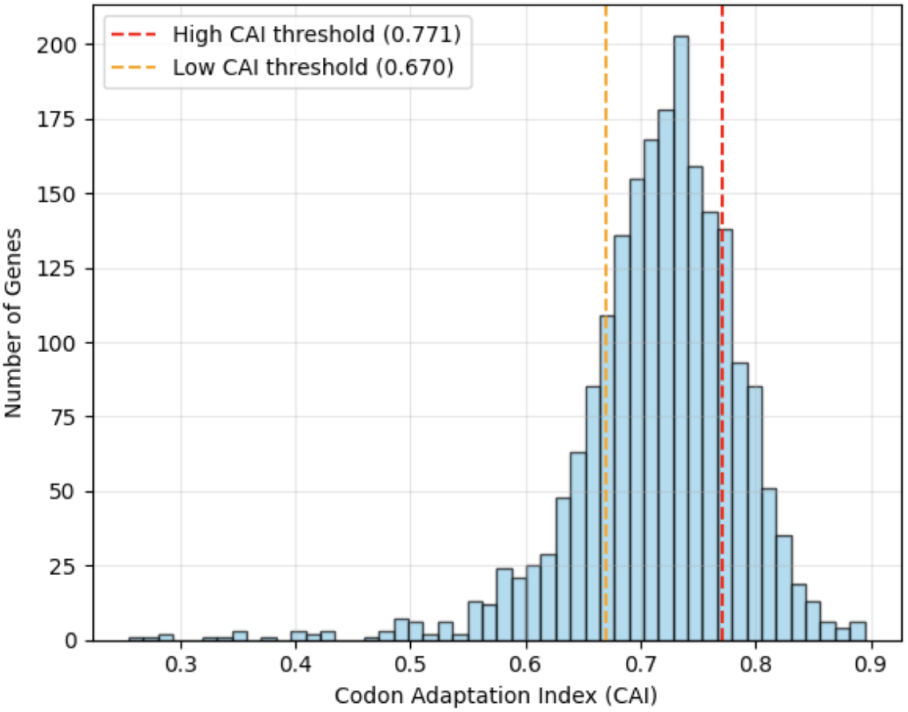
Distribution of Codon Adaptation Index (CAI) values of *F.nucleatum* subsp. *Nucleatum* ATCC 25586 genome. using ribosomal protein sequences as reference, calculated based on the API Reference (Bagnoli and Lìo, 1995; Lee, 2019). The mean CAI value is 0.716 (SD = 0.071) with values ranging from 0.256 to 0.895. The red line is the threshold for a high CAI (top 20%, CAI *≥* 0.771), while the yellow line is the threshold for a low CAI (bottom 20%, CAI *≤* 0.670). The distribution of genes with high CAIs is ribosomal proteins, translation factors (EF-G, EF-Ts, aminoacyl-tRNA syn-thetases), and metabolic or energy-related proteins (ATP-binding proteins, kinases, dehydrogenases). The distribution of genes with low CAIs is majoritly hypothetical proteins, with some other transport proteins (permeases and ABC transporters), metabolic or energy-related proteins (ATP synthase sub-units), and transcriptional regulators.

To characterize the genetic content of the predicted PAIs, gene sequences were downloaded from NCBI (GFF3 format), and corresponding protein sequences were extracted. Since a large number of hypothetical proteins were initially identified, protein sequences were queried against the UniProt database for id mapping and using BLAST to identify the top three potential identities for each protein (Altschul et al., 1990). The result, combined with annotations from NCBI and IslandViewer, was cross-referenced to make a best-fit prediction for each protein’s identity.

The functional profile for each region was done through a combined analysis of manual review, homology searches, and Biomni (Huang et al., 2025).

Region 1967193–1999047 was found to be dominated by genes related to metabolic functions. The proteins within this region primarily suggest a systemic or intracellular role, potentially related to nutrient acquisition or host interaction in deeper tissues.

Region 366847–372320 contains toxins, such as hemolysin, cluster. This region is of particular interest because it is the shortest of all 3 regions and only contains 10 identified open reading frames. Most of its genes could be associated with drug resistance, such as a nitroreductase family protein, acyl-CoA thioesterase, and a metallothionein. However, the presence of an SSU ribosomal protein S20P raises questions about its classification as a classic pathogenicity island (PAI). Generally, essential genes like those for ribosomes are not typically mobile genetic elements and are not associated with virulence.

Region 1496613–1523855 exhibited characteristics consistent with a pathogenicity island. It contains mobile genetic elements and the presence of toxin-antitoxin systems. Furthermore, this region corresponded directly to a significant fluctuation identified during the initial GC skew analysis.

### The Map of the Promoters reveals coexpression and gene organisation

The *F.nucleatum* ATCC 25586 genome contains 2,067 predicted protein-coding genes based on the National Institutes of Health (NIH) database. The PePPER analysis predicts that within this genome, there are 680 promoters, with promoter confidence scores ranging from 0.0 to 1.0, higher scores indicating stronger predicted promoter activity. All four promoters within PAI1 achieved the maximum prediction confidence score of 1.0, and 8 of 10 promoters in PAI2 scored *≥* 0.82, suggesting these regions may contain highly expressed elements. The genome-wide promoter-to-gene ratio is 0.33. When estimating 3 predicts PAI region using Biopython (Cock et al., 2009), PAI 2 and PAI 3 have promoter-to-gene ratios closely matching the genome average, with 0.31 and 0.33, respectively. However, PAI 1 has a notably higher ratio of 0.7. Focusing on PAI 1 and its surrounding genomic region, a hemolysin cluster is found located between 296–307 kb, which includes PAI1. Within PAI1 itself (coordinates: 366,847–372,320 bp), the gene at locus tag FN1885 encodes hemolysin III, a pore-forming cytolytic toxin with a potentially high pathogenic potential due to its direct membrane-damaging activity.

After promoter mapping of predicted PAIs, several functional clustering appear to suggest potential coexpression and co-regulation patterns. In PAI 1, the hemolysin III (locus tag FN1885) and the adjacent unknown protein (locus tag FN1886) on the forward strand, with only one promoter upstream, suggest a potential linkage in their transcriptional regulation. Hemolysin III (FN1885), as a pore-forming toxin, provides computational evidence consistent with virulence potential by allowing host cell lysis and immune evasion. The downstream unknown protein (FN1886) could imply an accessory factor that will either enhance the function of hemolysin or represent some form of interaction with the host in an unknown and uncharacterized manner.

In PAI 2 there appears to be a biotin synthesis cluster on forward strand FN0849 (synthase) → FN0850 → FN0851 (bioC) → FN0852 (hydrolase) and the glycogen metabolism operon on the reverse strand(FN0853-FN0858, all glucose related), potentially implementing co-adaptation to metabolic processes possibly coordinating vitamin biosynthesis with energy storage/utilization under nutrient limited situations. The 2 predicted operons do not directly contribute virulence factors, but may facilitate colonization in host cells. Biotin synthesis genes (FN0849–FN0852) may aid the survival in nutrient-restricted niches where the host limits vitamin availability. The glycogen metabolism operon (FN0853–FN0858) allows energy storage, supporting persistence during fluctuating nutrient conditions. Overall, both operons may contribute to pathogenicity through nutritional adaptation, helping the bacterium withstand stresses imposed by its host.

In PAI 3, 2 potential coexpression clusters could be RNA polymerase sigma factors cluster on the reverse strand with FN1317 to FN1318 (rpoD) and the close link metabolic enzymes with FN1322 (metalloprotease) → FN1323 (thymidylate kinase)→ FN1324 (reductoisomerase). One could be regulatory control of metabolic actives, while the other could function in tissue degradation (protease activity) and nucleotide/amino acid metabolism (thymidylate kinase, reductoisomerase), ensuring replication inside the host. Those 2 operons support PAI3’s role as a metabolic regulator but not as a pathogenicity island.

Analyzing the genomic environment of the gene with locus tag FN1885 includes 9 genes and 4 putative promoters, of which FN1885 is in a forward-strand gene arrangement with 7 other genes in the same direction and has no large intergenic gaps. The reverse-strand genes (FN1884 and FN1889) are interspersed but not aligned in the same transcriptional direction, making them less likely to be part of the same operon. Within the forward strand gene, there are 2 upstream promoters, particularly promoter 122, located proximal to FN1885, indicating overlapping or sequential transcriptions and coregulation of this gene cluster. Based on the promoter positioning, FN1885 is most likely coexpressed with FN1886 to 1889, potentially in response to signals for coordinated biological function.

### Codon Adaptation Index as a Proxy for Protein Abundance

High CAI values suggest strong codon optimization, likely for efficient translation. A large proportion of the top genes in the bacterial genome were ribosomal proteins (for example, L24P, S19P, L20P), which are typically highly expressed and exhibit high CAI values. Some non-ribosomal genes (e.g., flavodoxin, transcriptional regulators, hypothetical proteins) also show high CAI, which may indicate functional importance or horizontal gene transfer.

PAI1 (region 366847–372320) contains genes with moderate to high CAI, including a ribosomal protein and a hemolysin, which may suggest functional integration into the host genome.

CAI values ranged from 0.505 to 0.863, with most genes exceeding 0.65, indicative of moderate-to-high codon optimization relative to ribosomal gene reference sets. High CAI values in hypothetical proteins AAL93983.1 (locus tag: FN1884, CAI: 0.863) and AAL93978.1 (SSU ribosomal protein S20P, CAI: 0.828) suggest potential functional relevance, although one has uncertain annotation. At the same time, lower CAI scores in protein AAL93981.1 (locus tag: FN1882, CAI: 0.505) and AAL93982.1 (locus tag: FN1883, CAI: 0.581) may indicate either lower expression under standard conditions or acquisition from distinct genomic backgrounds via horizontal gene transfer. Functionally, the cluster contains a potentially highly pathogenic hemolysin gene and a likely coexpressed gene that could potentially facilitate the hemolysin pathway. Also, this region contains accessory hypothetical proteins and mobile genetic elements (transposases), consistent with horizontally acquired virulence modules. However, the inclusion of a ribosomal protein (housekeeping function) and metabolic enzymes (e.g., NAD(P)H nitroreductase) diverges from the traditional pathogenicity islands’ composition.

Despite containing a high CAI gene, promoter motif analysis of the gene in the PAI1 region shows a pattern highly similar to low-CAI promoters, suggesting limited or intermittent transcriptional activity. Pathogenic islands often contain virulence genes that are actively transcribed when needed (Hacker and Kaper, 2000; Zhao et al., 2018); a region with weak or inconsistent promoter motifs may indicate that these genes are not strongly expressed. Combined with previous analyses, PAI1 may be more likely a remnant of DNA from horizontal transfer, and a coexpressed cluster of virulence genes rather than a functional PAI.

PAI2 (region 1496613–1523855) mostly contains hypothetical proteins and metabolic enzymes. Inspection of the region via blast and UniProt ID mapping found several transposases and toxin-antitoxin protein-coding genes. The high-CAI hypothetical proteins, which could be recently acquired genes that are being adapted for expression, and the presence of metabolic enzymes, suggest potential functional roles in niche adaptation. Promoter motif analysis reveals poly-A-tracts, which are associated with strong DNA curvature (Fiorini et al., 2011). This has been further supported by DNA curvature analysis, which indicated that 7 of 10 annotated promoters exhibit pronounced macroscopic curvature. On the other hand, only 5 of 32 protein-coding genes within the same region show comparable curvature (Gohlke, 2025). This aligns with the established role of DNA shape in transcriptional regulation, which recognizes that intrinsic curvature tends to be concentrated in promoter regions than coding sequences in the same genome (Kozobay-Avraham, 2006) and is consistent with the identification of the promoters. DNA curvature enhances promoter–RNA polymerase interactions and stable open complex formation, which can associate with stronger transcription (Asayama and Ohyama, 2000). The presence of mobility genes, adaptive metabolic functionalities, and strong promoter motifs supports the interpretation that PAI2 represents a horizontally acquired genomic region with adaptive significance, and it is reasonable to propose that PAI2 may be a functional pathogenicity island, although additional experimental validation will be necessary to verify its role in virulence.

PAI3 (region 1967193–1999047) contains genes involved in translation and nucleotide metabolism, which are typically core functions. Core functional regions are generally not integrated from other genes.

## Proteomics

### Prediction of Protein interaction within PAI2

The genomic region PAI2 (1496613–1523855) encodes 32 protein-coding genes, several of which are associated with mobility elements (e.g., tyrosine-type recombinase/integrase and transposases), consistent with acquisition via horizontal gene transfer. To further characterize the functional organization of PAI2, we STRING database analysis combined with BLAST-based functional annotation of all 32 encoded proteins, transmembrane domain prediction, followed by AlphaFold-Multimer structural validation of predicted interactions (Hallgren et al., 2022; Jumper et al., 2021; Krogh et al., 2001; Szklarczyk et al., 2024).

STRING network analysis organizes the 32 proteins encoded within PAI2 into 4 visually connected components and 1 isolated protein, as shown in Figure 8a (Szklarczyk et al., 2024). The proteins showing minimal connectivity to the main network are FN0838 (uncharacterized). This suggests they could not be functionally linked to any pathway, likely represent housekeeping genes captured within the island boundaries rather than a core PAI2 component.

**Figure 6:**
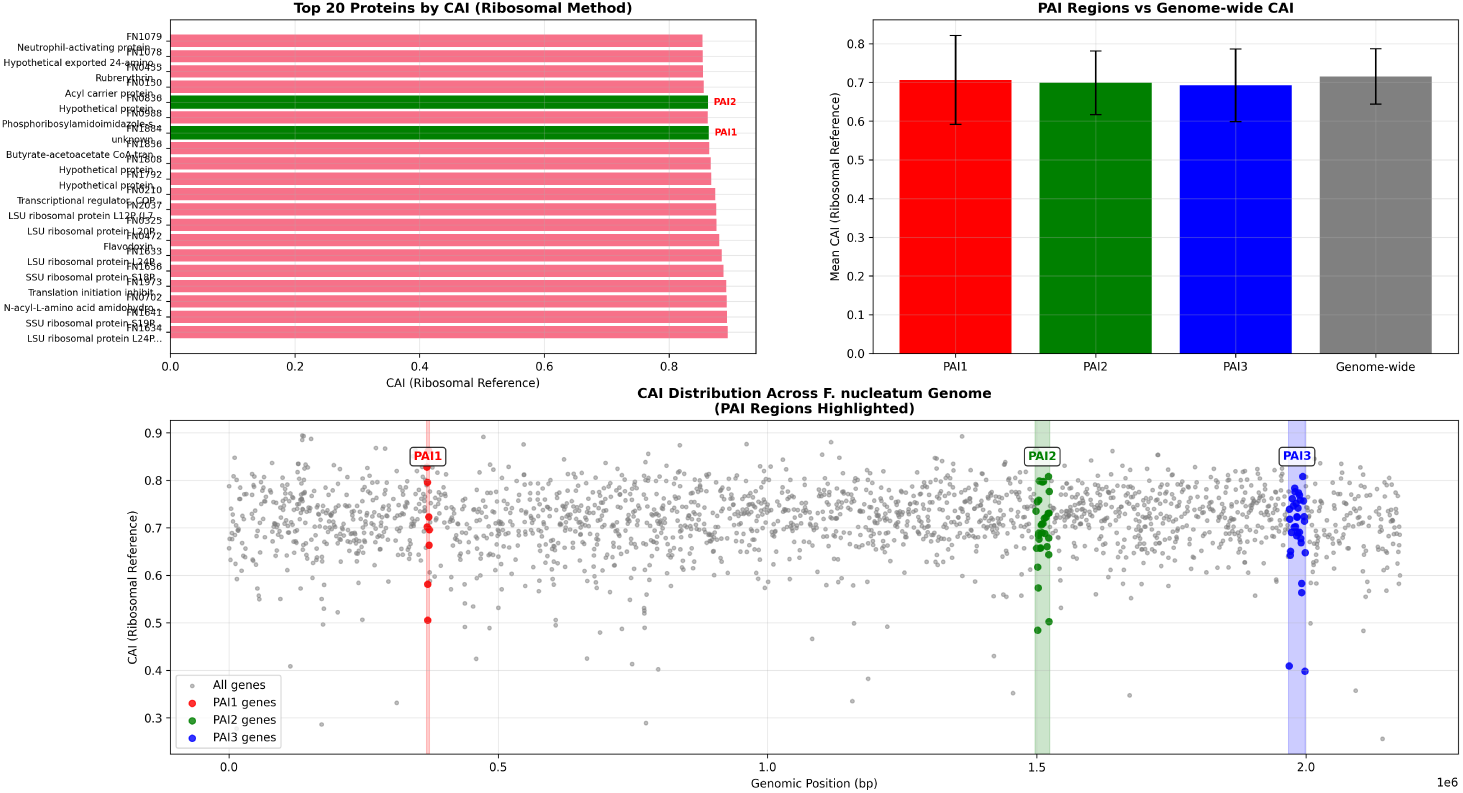
Distribution of the Codon Adaptation Index (CAI) along the genome of Fusobacterium nucleatum, with specific attention to potential Pathogenicity Islands (PAI1: 366,847–372,320; PAI2: 1,496,613–1,523,855; PAI3: 1,967,193–1,999,047). The analysis was performed using a ribosomal gene set as the reference for optimal codon usage.

**Figure 7:**
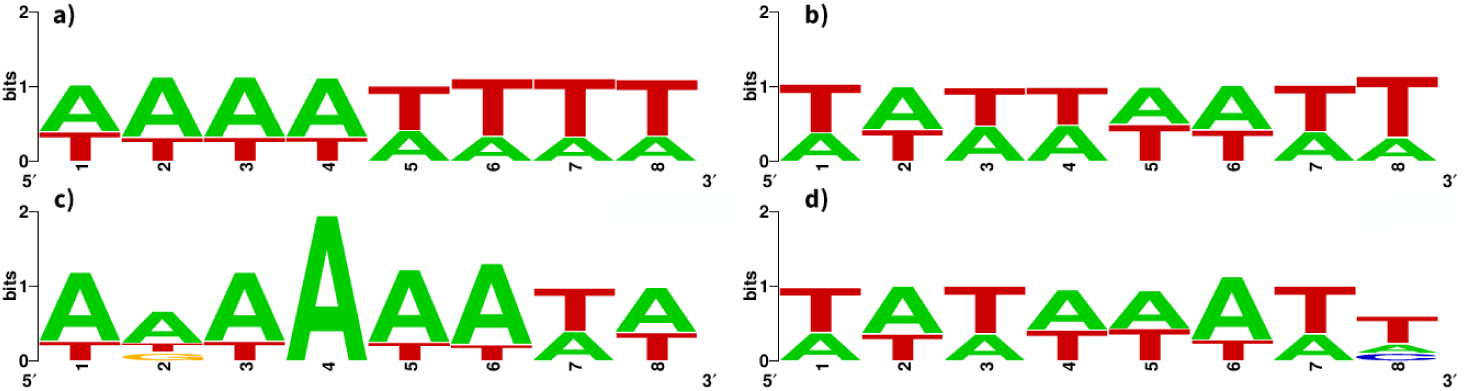
Promoter motif analysis of high and low CAI genes and PAI loci using WebLogo (Crooks et al., 2004; Schneider and Stephens, 1990). a) Sequence logo representation of enriched 8-mer motifs found in the promoters of high CAI genes (CAI *>* 0.771). Promoter motifs were found using the coding pipeline that calculates CAI, filtered sequences by minimum occurrence, sorted by frequency, and generated consensus motifs with WebLogo. b) Sequence logo representation of promoter motifs from low CAI genes (CAI*<* 0.670), generated using the same approach. c)Promoter motif analysis of the gene in PAI2. d)Promoter motif analysis of genes in PAI1.

**Figure 8:**
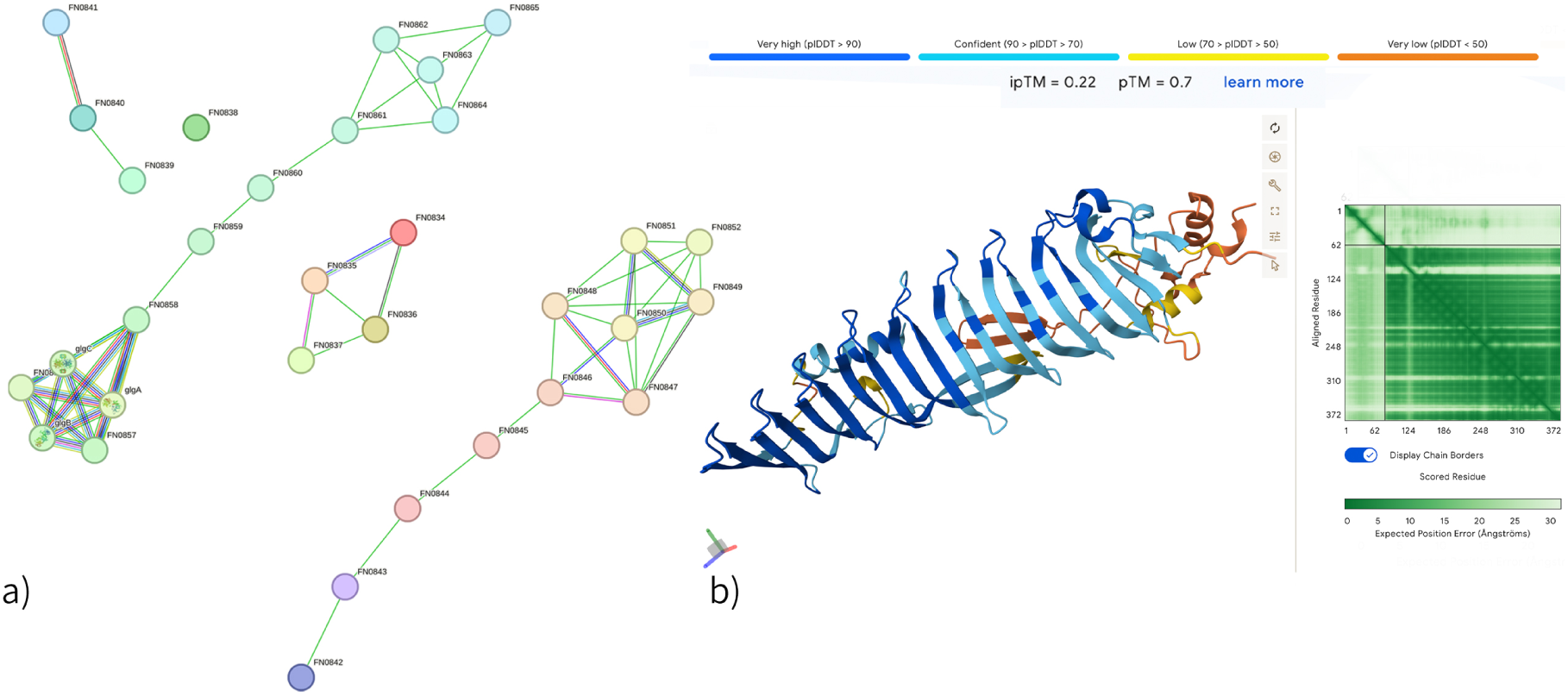
Predicting structural and interaction analysis for 32 proteins in region 1496613–1523855 (PAI2). a) STRING network analysis of predicted protein–protein interactions among proteins in PAI2 (Szklarczyk et al., 2024) b) AlphaFold-Multimer predicted 3D structure of FN0845 and FN0846 interaction with high pTM (0.70), suggesting confident structure prediction (Jumper et al., 2021).

**Figure 9:**
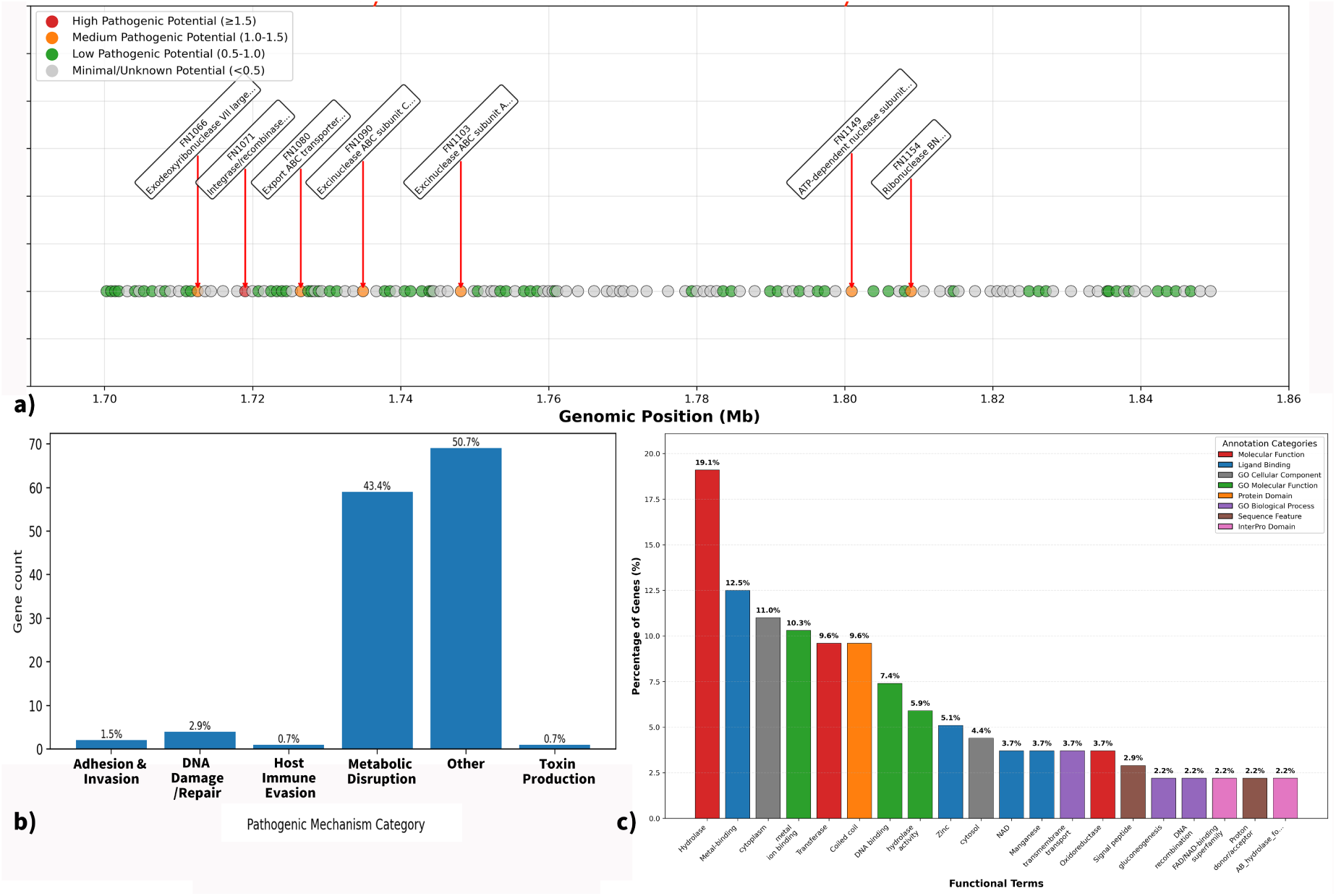
Functional and Pathogenic Annotation of the 1.7–1.85 Mb Genomic Region in *F.nucleatum* ATCC 25586. 1) Gene organization schematic for 1.70-1.85 Mb region. The genes of high and medium pathogenicity are labeled. Classification of potential pathogenicity was completed by Biomni’s (Huang et al., 2025) annotation pipeline. b) Predicted pathogenicity mechanisms among genes in these genomic regions based on PubMed-assisted text mining. c) The functional annotation output of DAVID for the same gene set (Huang et al., 2008; Sherman et al., 2022). The total percentages are higher than 100%, as this is a feature of DAVID where gene-term associations overlap and are counted multiple times across similar or redundant terms.

**Figure 10:**
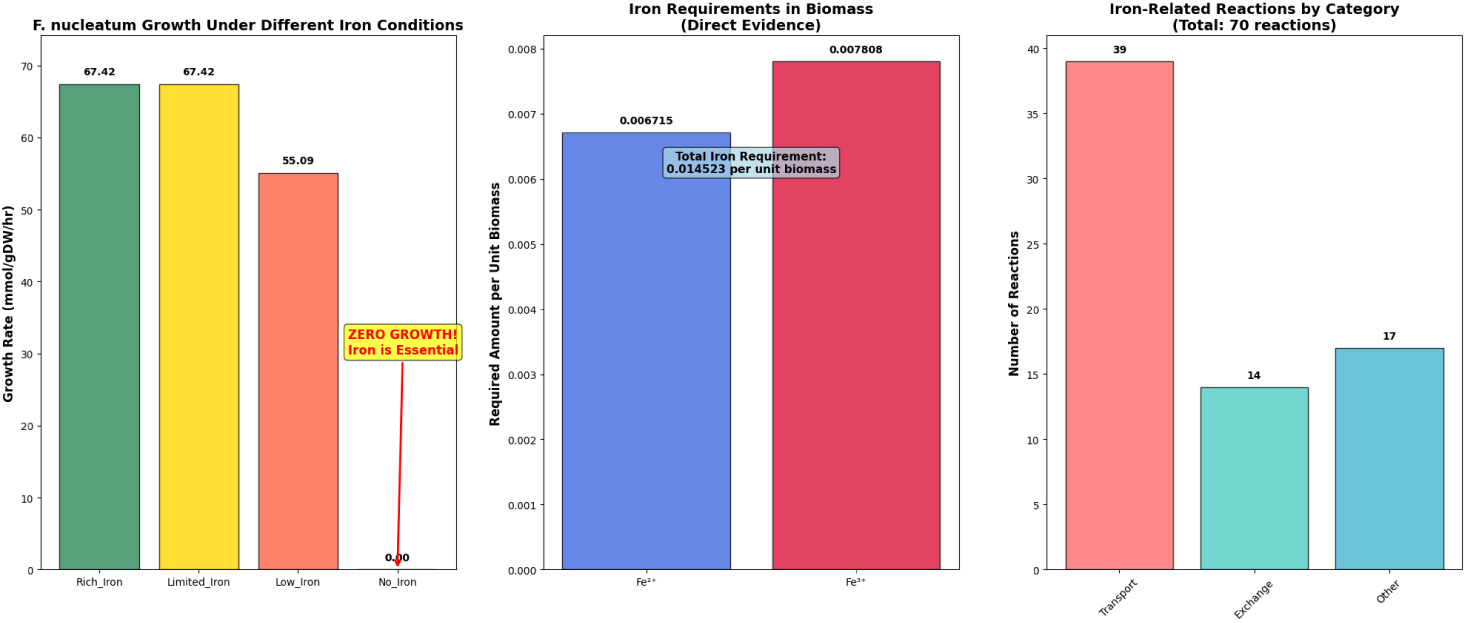
Multi-panel visualization that provides a comprehensive overview of iron’s essentiality for *F. nucleatum*, FBA flux analyzed using cobrapy. The panels show that iron deprivation completely halts growth, while rich and limited iron conditions support high growth rates, emphasizing iron’s metabolic importance. Additionally, the presence of diverse iron acquisition systems in the model supports iron critical role in survival and virulence.

Deep Learning tools, Biomni, analysis confidence scores derived from STRING interaction data using NetworkX reveal an average clustering coefficient of 0.560 and a network density of 0.138, indicating moderate connectivity with distinct functional modules (Huang et al., 2025). Community detection identified 6 functional modules, with 2 inter-module connections suggesting pathway crosstalk (FN0845 and N0846; FN0858 and FN0859).

A core glycogen metabolism cluster comprising six proteins represents the largest and most densely connected module observed. This cluster includes glycogen synthase (*glgA*, FN0853), glucose-1-phosphate adenylyltransferase (*glgC*, FN0854; *glgC2*, FN0855), 1,4-*α*-glucan branching enzyme (*glgB*, FN0856), glycogen phosphorylase (*glgP*, FN0857), and 4-*α*-glucanotransferase (*malQ*, FN0858), with STRING confidence interaction scores ranging from 0.976 to 0.999. The tight clustering of these enzymes suggests coordinated expression and potential physical interaction, which may facilitate efficient substrate channeling during glycogen metabolism.

Another prominent cluster is associated with biotin biosynthesis, including 8-amino-7-oxononanoate synthase (*bioF*, FN0849), pimeloyl-ACP methyl esterase (*bioG*, FN0850), malonyl-ACPO-methyltransferase (*bioC*, FN0851), and an *α/β* hydrolase (FN0852). These proteins exhibit confidence scores of 0.914 to 0.988 and are supported by gene neighborhood conservation and co-expression evidence across multiple bacterial species.

A third cluster comprises proteins with potential virulence association. BLAST analysis indicated that FN0846 belongs to the YwqK family of type II toxin-antitoxin systems, frequently linked to pathogenicity islands. FN0846 interacts with a TPR-repeat-containing protein (FN0847) and DUF4125 domain-containing protein (FN0848), with a confidence score of 0.971, suggesting functional cooperation in stress response or host adaptation. Notably, interaction is observed between this virulence cluster and the biotin biosynthesis module, suggesting that *F. nucleatum*’s virulence may depend on metabolic pathways such as biotin production. While there is no direct evidence linking biotin (vitamin B7) biosynthesis to virulence in *F. nucleatum*, this metabolic-virulence interface warrants further investigation.

A cluster containing a RiboL-PSP-HEPN domain protein (FN0845), a BLAST-verified DUF2335 domain-containing protein (FN0842), and two uncharacterized proteins (FN0843, FN0844) was identified. HEPN (Higher Eukaryotes and Prokaryotes Nucleotide-binding) domains possess RNase activity and are commonly found in toxin-antitoxin systems. The genomic proximity of FN0845 to the YwqK antitoxin (FN0846) and their STRING-predicted interaction suggests these proteins may constitute a functional toxin-antitoxin module, with FN0845 potentially serving as the cognate toxin.

In addition to metabolic and virulence clusters, clusters of mobility-associated proteins were identified, consistent with the horizontally acquired nature of PAI2. Four mobility-associated proteins, a DUF6882 domain-containing exported protein (FN0834), a DUF6882 domain-containing protein (FN0835), a Dabb family stress-response protein (FN0836), and a tyrosine-type recombinase/integrase (FN0837), where integrase likely mediates chromosomal integration of PAI2. The transposase cluster (FN0840 and FN0841) may facilitate internal rearrangements or mobilization events.

Out of 32 analyzed genes, eight encode uncharacterized proteins. FN0843 and FN0844 show a potential link with the HEPN domain module; FN0838 cannot be functionally related to known pathways, and FN0839 is in proximity to the transposase cluster (FN0840, FN0841), possibly implicating mobility.

The remaining hypothetical proteins appear to interact with each other and are predicted to be membrane-associated. This cluster includes a predicted lipoprotein (FN0859), a TM2 domain-containing protein (FN0862), and four uncharacterized proteins (FN0860, FN0861, FN0863, FN0864). The presence of a lipoprotein suggests surface localization and potential host-pathogen interaction. STRING analysis indicated functional associations based on gene neighborhood conservation, although the specific functions are undetermined.

### Structure Support

To further characterize these proteins, we performed transmembrane domain analysis using TMHMM2.0 and DeepTMHMM (Table 2) (Hallgren et al., 2022; Krogh et al., 2001).

**Table 1:**
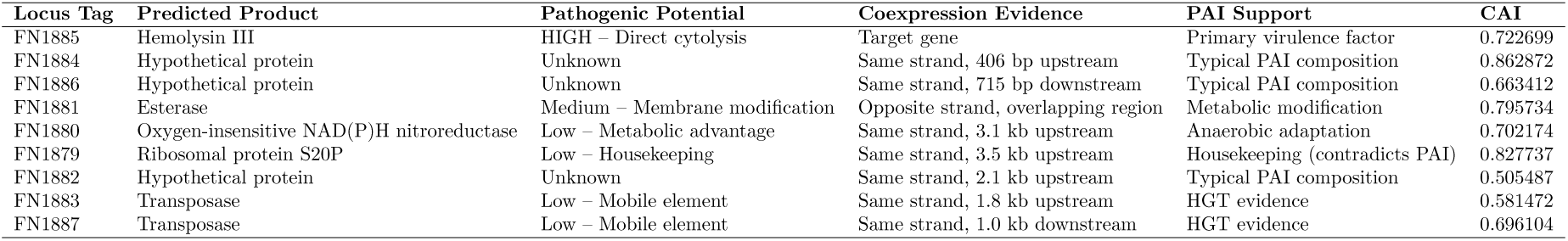
Summary of genes in the predicted PAI1 region in *Fusobacterium nucleatum*, including predicted products, pathogenic potential, coexpression evidence, genomic context, and Codon Adaptation Index (CAI) values. High CAI values suggest codon optimization.

| Locus Tag | Predicted Product | Pathogenic Potential | Coexpression Evidence | PAI Support | CAI |
| --- | --- | --- | --- | --- | --- |
| FN1885 | Hemolysin III | HIGH – Direct cytotoxicity | Target gene | Primary virulence factor | 0.722699 |
| FN1884 | Hypothetical protein | Unknown | Same strand, 406 bp upstream | Typical PAI composition | 0.862872 |
| FN1886 | Hypothetical protein | Unknown | Same strand, 715 bp downstream | Typical PAI composition | 0.663412 |
| FN1881 | Esterase | Medium – Membrane modification | Opposite strand, overlapping region | Metabolic modification | 0.795734 |
| FN1880 | Oxygen-insensitive NAD(P)H nitroreductase | Low – Metabolic advantage | Same strand, 3.1 kb upstream | Anaerobic adaptation | 0.702174 |
| FN1879 | Ribosomal protein S20P | Low – Housekeeping | Same strand, 3.5 kb upstream | Housekeeping (contradicts PAI) | 0.827737 |
| FN1882 | Hypothetical protein | Unknown | Same strand, 2.1 kb upstream | Typical PAI composition | 0.505487 |
| FN1883 | Transposase | Low – Mobile element | Same strand, 1.8 kb upstream | HGT evidence | 0.581472 |
| FN1887 | Transposase | Low – Mobile element | Same strand, 1.0 kb downstream | HGT evidence | 0.696104 |

**Table 2:** Comparison of TMHMM and DeepTMHMM predictions. Only proteins with TMHs are shown; displays number of predicted TMHs.

| locus_tag | protein_id | TMHMM | DeepTMHMM | product |
| --- | --- | --- | --- | --- |
| FN0839 | AAL95035.1 | 1 | 0 | unknown |
| FN0842 | AAL95038.1 | 2 | 2 | DUF2335 domain-containing protein |
| FN0844 | AAL95040.1 | 2 | 2 | unknown |
| FN0859 | AAL95055.1 | 1 | 0 | Lipoprotein |
| FN0862 | AAL95058.1 | 2 | 2 | TM2 domain-containing protein |
| FN0864 | AAL95060.1 | 2 | 0 | unknown |
| FN0865 | AAL95061.1 | 0 | 12 | Outer membrane protein |

Minor discrepancies were observed for FN0839, FN0859, and FN0864 between the two prediction methods. A notable difference was observed for FN0865, where TMHMM2.0 predicted no transmembrane helices, while DeepTMHMM predicted 12 membrane spanning regions. Detailed analysis of the DeepTMHMM output indicated that FN0865 is predicted to adopt a beta-barrel topology with alternating beta-sheets and periplasmic/extracellular loops, characteristic of outer membrane proteins (OMPs) in Gram-negative bacteria. This beta-barrel architecture, rather than alpha-helical trans-membrane domains, explains the discrepancy between prediction methods, as TMHMM2.0 focused on alpha helical TM topology (Hallgren et al., 2022). The OMP classification of FN0865 is consistent with its potential role in host-pathogen interactions and supports its inclusion as a virulence-associated factor.

To validate predicted protein-protein interactions and assess the structural plausibility of complex formation, we performed AlphaFold-Multimer analysis on protein pairs/complexes prioritized based on STRING evidence and functional relevance (Jumper et al., 2021). Interface predicted TM-score (ipTM) values above 0.5 indicate confident interactions, while predicted TM-score (pTM) values exceeding 0.7 reflect reliable overall folding.

The FN0845 and FN0846 pair (putative HEPN toxin and YwqK antitoxin) exhibited the highest overall structural confidence (pTM = 0.70). FN0846 formed a canonical beta-barrel fold characteristic of DNA-binding antitoxins, and FN0845 displayed a helical domain typical of HEPN-family toxins (Figure 8b). Although the interface confidence (ipTM = 0.22) was not ideal, suggesting transient or conditional interaction, the predicted architectures align with established toxin-antitoxin modules.

For the glycogen metabolism pair (FN0853 and FN0854), individual domains folded with moderate confidence (pTM = 0.55), but interface confidence was low (ipTM = 0.14). Despite a strong functional association (STRING score 0.999), this pattern supports their role as sequential enzymes in glycogen biosynthesis rather than as stable complex partners.

The predicted structure for the four-protein membrane-associated complex (FN0860, FN0861, FN0863, FN0864) had low overall confidence (pTM = 0.37, ipTM = 0.28), forming a helical bundle. The low confidence likely reflects the limitations of AlphaFold-Multimer in modeling membrane environments, as these proteins are predicted to be membrane-associated.

Among transmembrane protein pairs, FN0839 and FN0860 showed the highest interface confidence (ipTM = 0.32), with helical architectures suggesting potential membrane complex formation. However, overall low confidence scores for membrane protein pairs further underscore AlphaFold-Multimer’s constraints when predicting interactions dependent on lipid bilayer context.

### Co-expression relationships among PAI2-encoded proteins

To further support computational predictions of protein interactions, we employed Bayesian network structure learning using the bnlearn R package (Scutari, 2010). Expression-based evidence was obtained from GSE161360 (GEO), a publicly available RNA-seq data from Ponath et al. (2021), who characterized the transcriptional landscape of *F. nucleatum* across multiple subspecies and growth conditions. We extracted 18 samples corresponding to *F. nucleatum* subsp. nucleatum ATCC 25586, containing three growth phases (exponential, mid-log, and stationary) and two treatment conditions.

The original study focused on identifying novel non-coding RNAs and transcriptional start sites; here, we used their expression data and analysis with bioinformatics tools to infer protein co-expression networks within the PAI2 region (Ponath et al., 2021). Although optimal Bayesian network learning typically requires 30–50 samples, we are only focused on 32 genes rather than the entire genome (18 samples provide sufficient statistical power for meaningful inference).

Using the hill-climbing algorithm with BIC score and 500 bootstrap iterations, bnlearn identified 56 significant edges (bootstrap strength *>* 0.5, direction *>* 0.5). Comparison with STRING database predictions revealed 14 overlapping interactions supported by both expression-based and database-based evidence. The strongest overlapping interactions involve glycogen metabolism genes (FN0853 (glgA) and FN0854 (glgC), FN0856 (glgB) and FN0857 (glgP), FN0855 (glgC) and FN0856 (glgB)). This concordance is biologically expected, as these enzymes function coordinately in glycogen biosynthesis and degradation. Similarly, the biotin biosynthesis module showed strong co-expression((FN0849 (bioF) and FN0851 (bioC), FN0849 and FN0850, FN0850 and FN0852)).

For the hypothetical protein cluster (FN0860-FN0865), bnlearn detected several co-expression relationships not captured by STRING. There is interaction of FN0864 and FN0837 (strength = 0.774, direction = 0.651), FN0860 and FN0848 (strength = 0.620, direction = 0.619), and FN0859 and FN0848 (strength = 0.598, direction = 0.540). Moreover, within the PAI2 hypothetical protein seems to coexpress with each other even when not adjacent to each other, for instance, FN0861 exhibited strong co-expression with FN0844 (strength = 0.950), suggesting potential functional coordination between uncharacterized proteins in this region. Regarding the hypothesis that hypothetical proteins may interact at the membrane based on function and structure evidence, bnlearn data also provided partial support. FN0865 (outer membrane protein) showed co-expression with FN0842 (strength = 0.712), FN0849 (strength = 0.722), and FN0850 (strength = 0.706). FN0859 (lipoprotein) exhibited co-expression with FN0844 (strength = 0.638) and FN0848 (strength = 0.598). Additionally, FN0863 showed strong co-expression with FN0846 (exported protein; strength = 0.726).

The expression data is related limited on sample size (n=18), which may result in both false negatives (missed true interactions) and false positives (spurious edges). Expression-based evidence suggests the protein pair identified showed co-expression patterns, which may reflect shared regulatory control, membership in common pathways, or indirect functional relationships rather than direct protein-protein binding. The low overlap between bnlearn and STRING (only 14 pairs) is expected, as STRING aggregates literature and experimental evidence across organisms, while bnlearn captures condition-specific co-expression patterns unique to the experimental conditions examined.

### PAI predicted interactome

The integration of STRING network analysis, BLAST functional annotation, transmembrane domain prediction, and AlphaFold-Multimer structural validation suggests PAI2 as a mosaic genomic island with characteristics of metabolic adaptation, virulence, and mobility.

All computational tools used in this section, including STRING, AlphaFold-Multimer, bnlearn, none supported direct physical interactions among these hypothetical proteins. AlphaFold-Multimer interface predictions showed generally low confidence (ipTM *<* 0.5), indicating that predicted interactions may be transient, require additional cofactors, or occur in membrane contexts not captured by the algorithm. Furthermore, the functional associations predicted by STRING may represent co-expression or genomic proximity rather than direct physical interactions. Finally, 8 of 32 proteins remain functionally uncharacterized despite predictions made by computational analysis. To verify that those proteins actually form complexes and exhibit pore-forming or toxin-related activity in vitro, biochemical and functional assays will be necessary.

Taken together, our analysis of PAI2 suggests its function as a pathogenicity island combining mobility-associated genes, virulence factors including a toxin-antitoxin system, metabolic adaptation genes, and membrane-associated hypothetical proteins. The AlphaFold-Multimer structural validation of the FN0845 and FN0846 toxin-antitoxin pair provides structural evidence consistent with the virulence-associated nature of this island. The transmembrane domain predictions and STRING-derived interaction networks predicted that some uncharacterized proteins may assemble into membrane-associated complexes. However, these insights are primarily based on computational predictions. Further experimental validation is needed to confirm biological significance.

### Pathogenic cluster analysis and interesting pathogenic activity

Since region 1496613–1523855 is identified as a potential PAI, regions around it that also seem to deviate significantly from the normal GC skew pattern were analyzed. Adjacent to region 1496613–1523855, a 150-kb region (1.7–1.85 Mb), which contains 136 annotated proteins, was analyzed for pathogenic potential. The same process of ID mapping, homology-based search (BLAST), and functional catego-rization was performed on this region. Of the 136 proteins present in this region, 37/136 (27.21%) are unannotated in the NCBI database. The incomplete characterization of this region underscores the potential presence of novel or strain-specific functions.

Functional classification based on PubMed-assisted text mining showed that Genes for Metabolic Disruption had the highest abundance, while Genes for DNA Damage/Repair were second. This suggests that these regions may be playing a key role in the metabolic flexibility and / or genomic integrity of bacteria. DAVID functional annotation provided further support, where the most enriched categories were for Hydrolase activity (19.1%), Metal-ion binding (12.5%), and Ion Transport (11%). These functions are likely involved in host interaction, nutrient acquisition, and stress-facilitation, providing additional evidence consistent with possible pathogenic capacity. The biomni analysis needs to be interpreted cautiously, because the method of evaluating pathogenic potential is unknown, and the prediction of pathogenic protein products could potentially involve either virulence or normal cellular pathways.

Similarly, constraints are present for the PubMed-assisted text mining analysis. Most genes have roles in multiple pathways, and the same protein in different pathways may play entirely different roles. The abundance of hypothetical proteins limits functional interpretability, and automatically mined text introduces variability in the results based on the time of database indexing, revisions to entries, and the sensitivity of the keyword search. In addition, the percentage values in DAVID are greater than 100%; this is a recognized artifact resulting from overlapping gene-term associations, for instance, where genes belong to more than one functional category.

The analysis suggests that this area of the genome may represent a contribution to the overall pathogenic potential, and the metal-ion pathway potentially related to iron acquisition should be given special attention. However, these are computational results that may be strain dependent. Experimental evidence, likely involving transcriptomics or functional assays, could help validate the involvement of this region in virulence.

### The iron in “bacterial economy”: hemolysin and iron ions transporters

Hemolysin genes are identified throughout the genome. This gene encodes a TlyA family protein, present in phylogenetically distant bacteria, being primarily found in distantly related bacteria such as *Mycobacterium*, *Helicobacter*, and *Brachyspira* species, rather than within the Fusobacteriota phylum (Arenas et al., 2011; Rahman et al., 2015). This TlyA protein shows pathogen-specific distribution patterns, where it is observed in pathogenic species like *M. tuberculosis* and *H. pylori* but is absent in non-pathogenic relatives, suggesting a strong selective pressure for its retention in pathogens (Keith et al., 2022; Sharma and Singh, 2022). Functionally, hemolysin operates as a pore-forming cytotoxin that lyses host erythrocytes, releasing iron and nutrients, allowing F. nucleatum to acquire nutrients in inflamed periodontal pockets (Ma et al., 2023; Samainukul et al., 2019).

### Hemolysin and iron acquisition mechanism in F. nucleatum

The identification of hemolysin suggests that F. nucleatum (Fn) may specialize in a heme utilization pathway to acquire iron from the host, indicating its potential ability to colonize and thrive in blood-rich environments such as periodontal pockets and tumor microenvironments.

From genome annotation, key proteins and genes involved in the heme-based iron acquisition pathway are identified. This system relies on ABC transporters (HmuUV), which comprise Hemin Transport System Permease Protein HmuU (encoded by genes hmuU 1 and hmuU 2) and the Heme Import ATP-Binding Protein HmuV (encoded by gene hmuV). HmuV, as the ATP-binding component, provides the energy to power HmuU. HmuU then facilitates the physical transport of the heme molecule from the blood and tissues across the cell membrane to the bacterial cells. While these proteins are dedicated to heme uptake, other ABC transporters for various metals, such as an Inner Membrane ABC Transporter Permease Protein (ydcV), a Zinc Import ATP-Binding Protein (znuC), Manganese ABC transporter substrate-binding lipoprotein (psaA), Iron import ATP-binding/permease protein (irtA), and many others, were also identified, highlighting a broader strategy for metal acquisition (Delepelaire, 2019; Louvel et al., 2006; McGregor et al., 2023).

Following import, the heme molecule is enzymatically degraded to release iron, a process in which flavodoxin (encoded by fldA, ispg, and isiB; highly expressed, cai of 0.880656) acts as an electron-transfer protein, reducing the ferrous complex and facilitating catalysis (Boynton et al., 2009; McGregor et al., 2023).

The iron is processed so it can be used by the cell for various cellular processes. In addition to basic metabolism sustaining bacteria, these iron acquisition mechanisms may disrupt erythrocytes and potentially promote metastasis in colorectal cancer. The virulence protein, DNA protection during starvation protein (Dps), for instance, serves as a bacterial ferritin to protect DNA from oxidative stress while also acting as a potent pathogenic factor. Besides disrupting and lysing the host’s erythrocytes to acquire iron, Dps also upregulates the chemokine CCL2/CCL7 to enhance the bacterium’s survival in macrophages. The expression of CCL2/CCL7 also contributes to cancer progression by inducing epithelial-mesenchymal transition (EMT) and promoting colorectal cancer (CRC) metastasis in vivo (Wu et al., 2023).

Genomic evidence showed complete iron acquisition machinery, and *F. nucleatum* relies on iron-dependent enzymes for core metabolic processes. An additional metabolic model analysis is done to see if iron is essential for *F. nucleatum* survival and proliferation.

### How scavenging iron helps metabolic growth

The annotated genome *F. nucleatum* ATCC 25586 was utilized with CarveMe (Machado et al., 2018) to reconstruct a genome-scale metabolic model, comprising 1,914 reactions and 1,339 metabolites, with 582 associated genes. Flux Balance Analysis (FBA) was applied to determine the optimal growth rate, which was predicted to be 67.42 mmol/gDW/hr under ideal nutrient-rich conditions. The flux through the biomass reaction represents the rate at which the bacterium produces those metabolites necessary for its growth (e.g., amino acids, lipids, cofactors, and proteins). The biomass flux returned by COBRA is the specific growth rate that can be used to calculate the specific biomass of the bacteria at a specific time using an exponential function. Therefore, biomass flux can be used as an indicator of the growth rate of the bacterium. Iron is required in a minimal medium for maintaining the growth rate as indicated by the CarveMe-generated metabolic model. The model identified uptake fluxes of 0.452702 mmol/gDW/h for ferrous iron (EX fe2 e) and 0.524388 mmol/gDW/h for ferric iron (EX fe3 e) as necessary to maintain the organism’s growth rate in a minimal medium.

Although it contains iron requirements in the composition of the biomass and the model contains reactions involving iron metabolites, COBRApy analyses performed using the metabolic model gener-ated by Carveme evaluate biomass production under all iron conditions, including complete absence of iron, limitation of iron uptake, and unlimited iron availability, give identical results (67.416545 mmol / gDW / hour). This indicates missing iron dependencies or unrealistic constraints in the model.

The failure of the stimulation could be explained by FBA’s inherent reliance on functional optimization in the face of biological redundancy. The metabolic networks contain multiple iron-chelate compounds (enterobactin, salmochelin, heme) and alternative metabolic pathways that can perform equivalent functions, masking true iron dependencies. In this simulation, only the EX fe2 e and EX fe3 e are set to (0,0), while other iron exchange reactions allow unrealistic, unlimited iron uptake. The FBA algorithm, pursuing optimization of biomass flux output, can simply re-route metabolic flux through these redundant pathways. Consequently, the simulation predicts no impact on growth, leading to a classification of the gene as non-essential (Boone, 2025).

**Figure 11:**
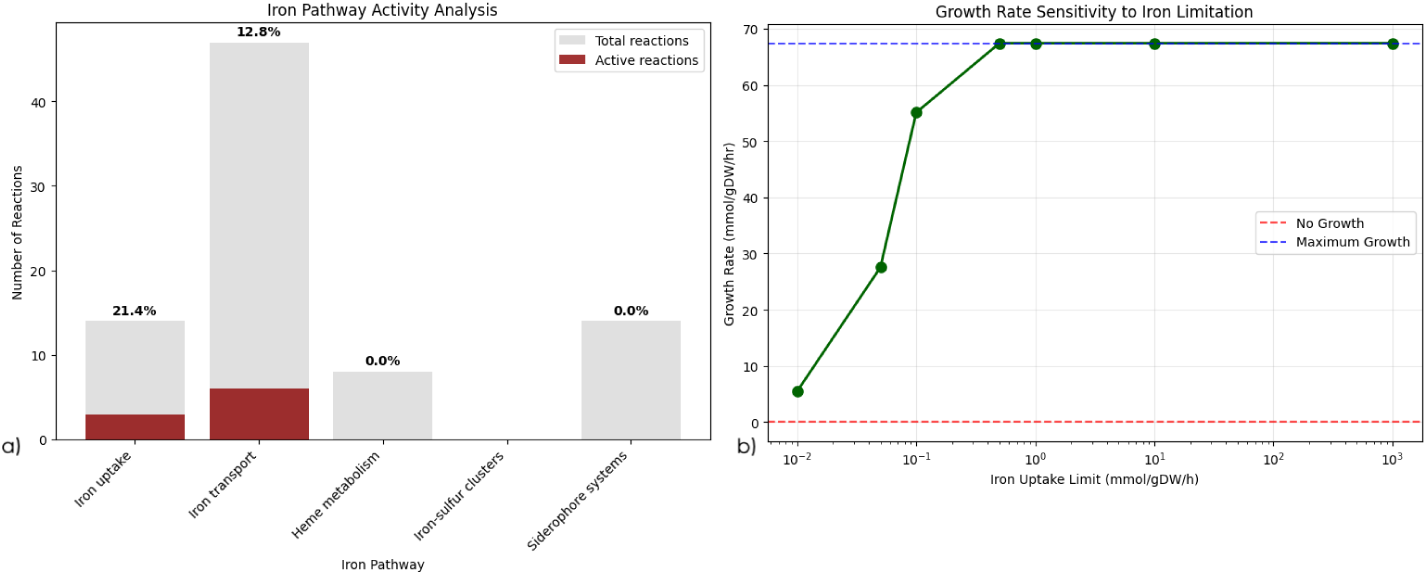
Detailed flux analysis, sensitivity curves, and pathway activity, using COBRApy. a) Bar graph compares the total and active reactions across five iron-related metabolic pathways: Iron Uptake, Iron Transport, Heme Metabolism, Iron-Sulfur Clusters, and Siderophore Systems. Grey bars represent the total number of reactions, while red bars indicate the subset of reactions that are active under the studied conditions. Percentages above each bar denote the proportion of active reactions, highlighting differential pathway utilization and potential regulatory prioritization. b) This plot illustrates the relationship between iron uptake limits and cellular growth rate. The x-axis shows iron uptake limits (mmol/gDW/hr) on a logarithmic scale, while the y-axis indicates growth rate (mmol/gDW/hr).

Identifying this potential limitation, 70 reactions and 40 metabolites related to iron across all three cellular compartments (cytoplasm, periplasm, and extracellular space) are used as the new constraints. The refined code indicates iron is required in biomass composition, for *F. nucleatum* cannot produce biomass without iron. FBA under different iron availability conditions is then performed. The model predicts that complete iron deprivation results in the arrest of growth. The metabolic consequences of iron limitation show that although iron is essential for bacterial growth, it only requires a limited amount of iron for maximum growth (*<* 1 *mmol/gDW/h*). Under iron-limited conditions, the model shows a compensatory activation of multiple siderophore systems to maximize scavenging, although this still results in an 18.3% reduction in growth rate. In addition, the bacterium possesses complex iron acquisition mechanisms that involve direct iron transport (ferrous iron (Fe^2+^ transport, ferric iron (Fe^3+^) transport, and ABC transporters), siderophore-mediated iron acquisition (enterobactin system, salmochelin system, and staphyloferrin systems), and heme iron uptake.

Furthermore, this bacterium appears to demonstrate a strong preference for ferric iron (Fe^3+^) over ferrous iron (Fe^2+^). An examination of iron transport systems in *F. nucleatum* using the metabolic model generated by Carvme from annotated Prokka through an AI-assisting tool, Biomni. The results suggested that 77.7% of total iron transport within the model is dedicated to Fe^3+^ uptake, with direct ferric iron uptake flux being 0.526, ferripyoverdine complex formation flux being 0.526, and Fe^3+^-pyoverdine exchange flux being 0.526. The direct ferrous iron transport flux is only 0.453.

Ferrous iron (Fe^2+^) is usually more soluble over a larger pH range than ferric iron (Fe^3+^), which hydrolyzes and precipitates as iron(III) hydroxide at near-neutral and higher pH values. The preference of *F. nucleatum* for Fe^3+^ may represent a metabolic adaptation to obtain iron in acidic niches, typically seen in tumor microenvironments.

**Figure 12:**
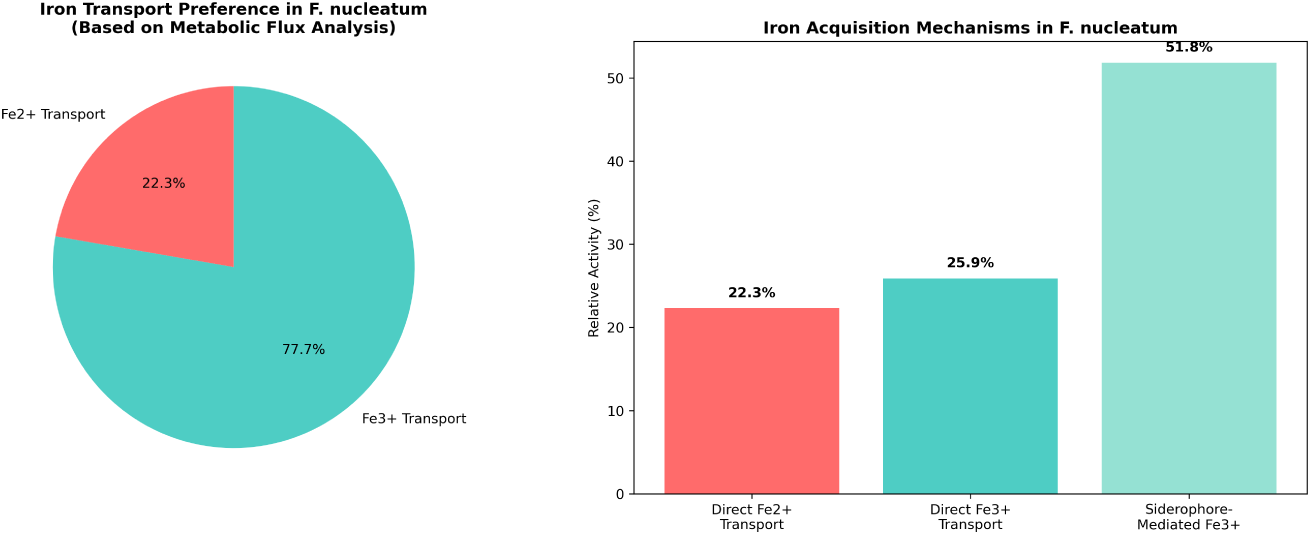
Examination of iron transport systems in *F. nucleatum* using the metabolic model generated by Carvme from annotated Prokka through an AI-assisting tool, Biomni.(Huang et al., 2025)

### F. nucleatum and cancer

Previous studies show *F.nucleatum* is frequently found in cancer tissue as well as associated with promoting tumor progression, supporting colonization in the cells, and altering the immune response within the TME. High levels of *F. nucleatum* is associated with the number of tumor-promoting immune cells, including tumor-associated macrophages (TAMs), tumor-associated neutrophils (TANs), dendritic cells (DCs), and myeloid-derived suppressor cells (MDSCs), and a decrease in the function of T-cells and natural killer (NK) cells. *F. nucleatum* contains virulence factors, such as Fap2, FadA, RadD, and FomA, which enable it to interact with cancer cells, modulating key signaling pathways, altering cytokine secretion, and promoting cancer progression. This bacterium can also secrete outer membrane vesicles that facilitate metastasis by promoting autophagy in oral cancer (Rubinstein et al., 2013; Willenbockel et al., 2024).

Moreover, lysozyme inhibitor Fn1792, a recently identified virulence factor, inhibits the apoptosis of host phagocytes through CX3CR1 expression. CX3CR1 expression can recruit CX3CR1^+^ PD-L1^+^ phagocytes to tumor tissues to adapt their microenvironment, resulting in immune evasion (Chen et al., 2024). *F.nucleatum* has lysed erythrocytes to free hemoglobin for growth as an iron source. Iron accumulated in macrophages enhances Fn-induced chemokine production through the TLR4/NF-*κ*B signaling pathway (Yamane et al., 2022). Therefore, iron availability may be predicted to affect the CX3CR1 induction by Fn1792, where further experimental studies would be required to confirm the possibility.

Recently, emerging evidence suggests it also contributes to mutagenesis (genomic instability) and DNA damage, particularly in the context of chronic inflammation. *F.nucleatum* contributes to mutagenesis indirectly through inflammatory responses and directly through metabolite production rather than directly inserting mutations, but these interactions accelerate the acquisition of mutations in tumor cells. One potential interpretation is that inflammation signals by *F.nucleatum* enhance the production of reactive oxygen species (ROS) and recruit inflammatory cytokines (Zhao et al., 2022).

Our genomic analysis identified a predicted hemolysin cluster (coordinates: 366,847–372,320 bp). Hemolysin III (FN1885) with CAI of 0.723 (moderate-high codon optimization) is a pore-forming cytotoxin with direct membrane-damaging activity. Promoter mapping indicated that FN1885 shares a promoter with the downstream hypothetical protein FN1886, suggesting coordinated expression. This genomic organization is consistent with a functional virulence module acquired through horizontal gene transfer, as evidenced by the presence of flanking transposases (FN1883, FN1887).

Hemolysis is generally associated with iron overload and excess hemoglobin in the bloodstream. Hemolysins lyse red blood cells, releasing iron, indirectly increasing iron concentration in blood (Millot et al., 2016). Free iron undergoes Fenton reactions, generating reactive oxygen species (ROS) such as hydroxyl radicals Fe^2+^ + H_2_O_2_ *→* Fe^3+^ + OH *·* +OH*^−^*, causing oxidative damage to lipids, proteins, and DNA (Xuan et al., 2025; Zhao, 2019).

### Hemolysis expression is context dependent

A critical observation is that hemolysin genes are typically not highly expressed under standard laboratory culture conditions. Transcriptomic analysis of *F. nucleatum* subsp. animalis during Caco-2 cell infection indicated that hemolysin III was among only 5 genes consistently upregulated during invasion (Cochrane et al., 2020). Computational essentiality analysis using NetGenes identified 202 essential genes in *F. nucleatum ATCC 25586* (Database, 2024). Several entries in the NetGenes essential gene set lacked specific functional annotation. To address this, we cross-referenced all essential genes with genome annotation from the NCBI GenBank database (NCBI accession: AE009951.2). 20 essential genes with predicted roles in iron homeostasis and transport is identified. Among these, FbpC (FN0376), a central ATP-binding protein for *Fe*^3+^, several ABC transporters, and additional iron transport genes were included in the essential gene set. Notably, hemolysin genes were not among the essential genes. This is consistent with the understanding that virulence factors are dispensable for in vitro growth but critical for in vivo colonization and pathogenesis (Kitamoto et al., 2015; Los et al., 2013; Nicholas et al., 1999; Opal et al., 1990). The absence of hemolysin from the essential gene set, combined with transcriptomic evidence of host-induced expression, suggests hemolysin represents a conditionally expressed virulence factor activated specifically during host infection. This explains why hemolysin activity may be underestimated in laboratory studies while being highly relevant in clinical contexts such as tumor colonization.

### Proposed mechanism: Hemolysin-Iron-ROS-Hippo pathway

Our flux balance analysis (FBA) using a genome-scale metabolic model indicated that iron is essential for *F. nucleatum*. Through genomic analysis of *F. nucleatum strain ATCC 25586*, direct cytotoxicity virulence factors are identified (shlB 3, shlB 2, shlB 1, tlyA). Iron acquisition by *F. nucleatum* through hemolysis is proposed to be responsible for *F. nucleatum* modulation of the hippo pathway, which may allow *F. nucleatum* to play a role in cancer progression and therapy resistance (Han et al., 2020; Millot et al., 2016; Wang et al., 2024).

Hemolysin enables *F. nucleatum* to acquire iron through a specific pathway. Hemolysin secretion leads to pore formation in erythrocyte membranes, releasing hemoglobin into the extracellular environment (Samainukul et al., 2019). Intracellular heme catabolism liberates iron, which participates in the Fenton reaction, generating reactive oxygen species (ROS) (Xuan et al., 2025; Zhao, 2019). ROS inhibit LATS1/2 activity, leading to YAP/TAZ dephosphorylation and nuclear entry. Activated YAP-TEAD signaling drives anti-apoptotic responses by upregulating BCL-2 and suppressing caspase-3–mediated GSDME cleavage, thereby reducing host cell pyroptosis(Amanda et al., 2024; Chen et al., 2017; Wang et al., 2024).

**Figure 13:**
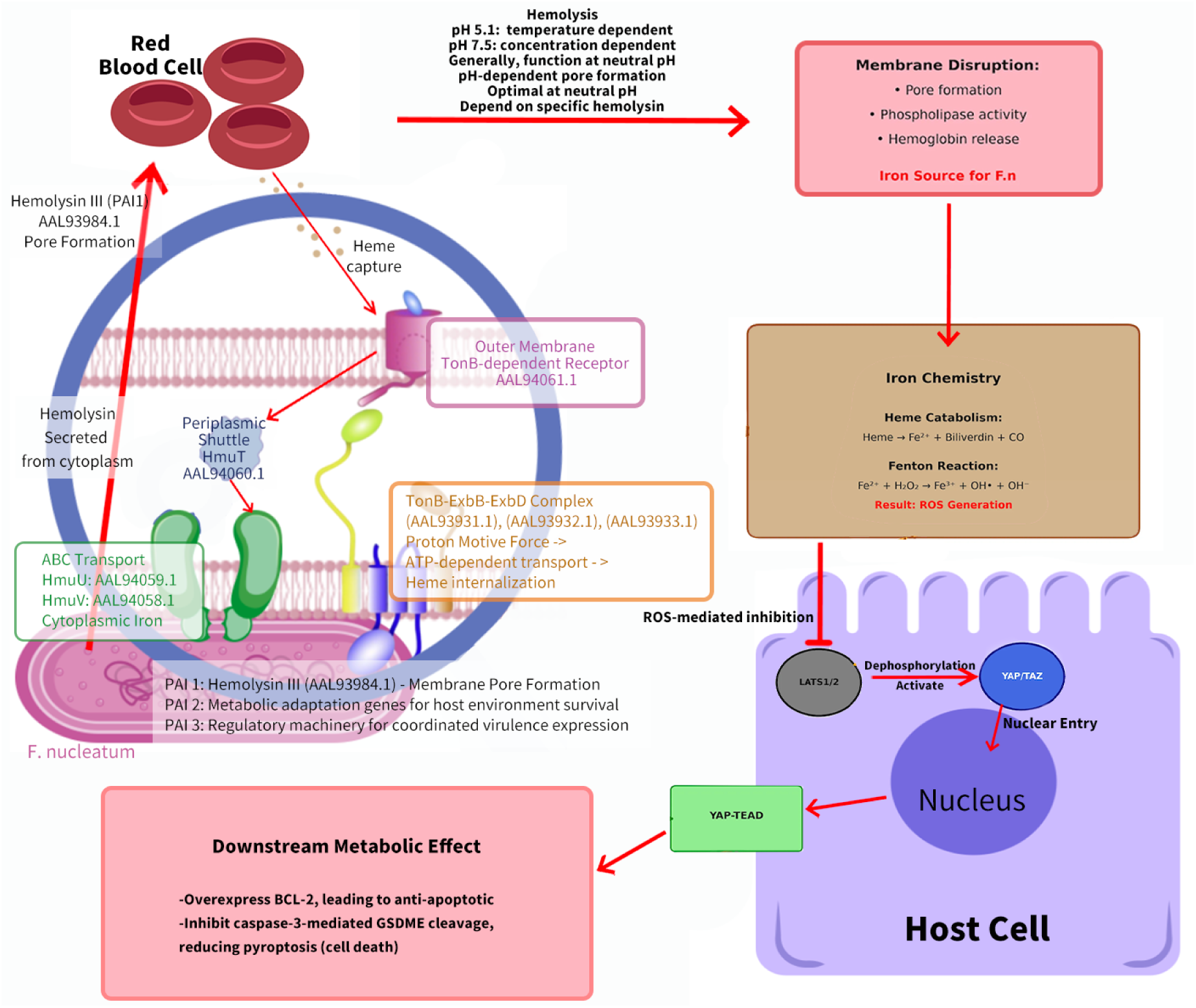
We propose that *F. nucleatum*’s iron acquisition through hemolysis has downstream effects on host cell signaling that promote cancer progression. The pathway produces reactive oxygen species (ROS), which could be associated with oxidative DNA damage (Albesa et al., 2004). Key proteins involved in this pathway, including TonB-dependent receptors, HmuTUV ABC transporters, and the TonB-ExbB-ExbD energy transduction system, with their respective protein IDs, are labeled (Braun et al., 2023; Richard et al., 2019; Silale and van den Berg, 2023).

Iron-induced oxidative stress inhibits LATS1/2 kinases, which dephosphorylate and nuclear translocate YAP and TAZ, the downstream effectors of the Hippo pathway. YAP/TAZ overactivation can promote pluripotency, tumor initiation, and chemotherapy resistance, which are common signs in cancer cells. They upregulate multidrug transporters (e.g., ABCB1, ABCC1, ABCG2) and anti-apoptotic proteins (e.g., Bcl-xL, cIAP1, Survivin), while suppressing p53 via MAPK/NF-κB signaling. Furthermore, YAP/TAZ activate TEAD-mediated transcription, and BCL-2 is a downstream target of the YAP-TEAD1 transcriptionally active complex (Amanda et al., 2024; Moroishi et al., 2015; Nguyen and Yi, 2019; Xuan et al., 2025). BCL-2 regulates apoptosis, and its overexpression is found in over 50% of oral squamous cell carcinoma (OSCC) tissues. The balance between pro-apoptotic (e.g., BAK, BAD) and anti-apoptotic (e.g., BCL-2, MCL-1) proteins determines tumor cell survival (Chen et al., 2017). *F. nucleatum*’s iron-mediated activation of YAP/TAZ may thus contribute to OSCC progression and resistance to anti-tumor therapies.

Further annotation of the Prokka-annotated genome was performed using eggNOG-mapper (see supplementary file FN.annotations.xlsx) (Huerta-Cepas et al., 2018). The result indicated the presence of the adhesin protein FadA (Fusobacterium adhesin A), but didn’t direct the identification of Fap2 or RadD in their classic forms. However, related proteins with similar functions were identified.

Two FadA proteins were found in the dataset (EJMLJHKB 00032 and EJMLJHKB 00869), both anno-tated as ”Adhesion protein FadA” with the FadA protein family domain (PFAMs: FadA); several other proteins with the FadA binding domain were also identified. FadA can activate the *β*-catenin and Wnt pathway via the binding to E-cadherin on the surface of CRC cells, causing relocation. Under stress conditions, FadA increase pathogenicity via conformational changes (Chen et al., 2022; Rubinstein et al., 2013).

While direct Fap2 was not found, several galactose-related proteins were found, including galac-tose kinase (EJMLJHKB 00595) and galactose-1-phosphate uridyl transferase (EJMLJHKB 00596), along with trimeric autotransporter adhesins (EJMLJHKB 01075 and EJMLJHKB 01330) which have structural homology with Fap2. Fibroblast activation protein 2 (Fap2), an outer membrane protein from F. nucleatum, is a galactose-sensitive hemagglutinin and adhesive protein that facilitate bacterial co-aggregation and host cell adhesion through binding to D-galactose-*β* (1-3)-N-acetyl-D-galactosamine (Gal-GalNAc) on host cell surfaces. Fap2 can bind to inhibitory receptors on immune cells, preventing immune recognition and facilitating immune evasion (Coppenhagen-Glazer et al., 2015; Ganesan et al., 2019). Although RadD is not directly present, multiple DNA repair and recombination proteins were identified, including RecG helicase (EJMLJHKB 00161), DNA topoisomerases (EJMLJHKB 00613, EJMLJHKB 00614), transcription-repair coupling factor (EJMLJHKB 00637), and Holliday junction re-solvase RuvC (EJMLJHKB 00821). RadD-related proteins function in DNA repair and recombination pathways. The identified protein also has a role in both of these pathways.

Our analysis identified one hemolysin cluster and a potential pathogenicity island, which could contribute to the proposed mechanism. PAI1 (366,847–372,320) may provide effector (hemolysin), initiating iron acquisition and membrane disruption. PAI2 (1,496,613–1,523,855), with metabolic adaptation genes (biotin synthesis, glycogen metabolism), may support survival in nutrient-limited host environments.

### Conserved Mechanism Across Pathogens

The hemolysin-ROS-genotoxicity pathway is not unique to *F. nucleatum*. Similar mechanisms have been documented in other cancer-associated bacteria, such as *E. coli* (Jamali et al., 2026; Nhu et al., 2019; Trouillon et al., 2023). *Uropathogenic Escherichia coli* (UPEC) originates from the gut, migrates to the urethra, is asscioated with bladder cancer. It secretes α-hemolysin (HlyA), a pore-forming toxin that disrupts mitochondrial function and promotes the generation of reactive oxygen species (ROS) in targeted host cells (Brito et al., 2018). ROS burst involves *E. coli* can lead to oxidative stress in blood, oxidative damage to macromolecules (proteins and lipids), oxidative DNA damage(Albesa et al., 2004; Baronetti et al., 2013). ROS can cause DNA damage during cell replication, and subsequently, mismatched or unpaired damaged DNA can cause the formation of mutations (Verma et al., 2020;

Waris and Ahsan, 2006).In specialized cells like spermatozoa and renal epithelial cells, exposure to hemolytic *E. coli* strains leads to increased intracellular ROS, which subsequently causes DNA strand breaks and genomic instability (Tvrdá et al., 2022).

The hemolysin-ROS pathway in bacteria can function as a mutagenic mechanism, particularly by inducing oxidative stress that causes DNA damage and subsequently increases mutation rates. This is not only proven in *E. coli*, but also shown in *Staphylococcus aureus*, which secretes α-toxin (Hla) leading to ROS production and resulting in DNA damage and apoptosis (Deplanche et al., 2019; Ha and Edwards, 2021; Menzies and Kourteva, 2000). Given the conserved mechanisms among pore-forming toxins, it is plausible that *F. nucleatum* hemolysin acts similarly.

However, not all bacteria containing hemolytic activity are associated with cancer, such as *Streptococcus pyogenes*, *Vibrio cholerae*, *Bacillus cereus*. S. pyogenes produces two major hemolysins, streptolysin O (SLO) and streptolysin S (SLS), which activate inflammasomes, induce pro-inflammatory cytokine release, and cause tissue destruction during acute infections such as pharyngitis, necrotizing fasciitis, and toxic shock syndrome (Brouwer et al., 2023; Langshaw et al., 2023; Pinho-Ribeiro et al., 2018; Richter et al., 2021; Shannon et al., 2024). Similarly, V. cholerae disrupts cell membranes and triggers inflammatory responses by β-barrel pore-forming hemolysin/cytolysin (VCC) (Gandhi et al., 2025; Singh et al., 2025). B. cereus produces multiple hemolysins, including hemolysin BL (Hbl), non-hemolytic enterotoxin (Nhe), and cytotoxin K, which cause food poisoning manifesting as diarrheal or emetic syndromes (Jovanovic et al., 2021). However, all are acute infections with a transient nature, which provides no opportunity for the persistent bacterial-epithelial interaction required for oncogenic transformation (Mager, 2006; Parsonnet, 1995).

Bacteria epidemiologically linked to cancer have the ability to establish chronic, persistent colonization of host tissues. The long persistence allows cumulative DNA damage from sustained oxidative stress and chronic immune activation. The proposed hemolysin-ROS-Hippo pathway alone is insufficient for cancer promotion; it requires sustained exposure to be oncogenically relevant (Jain et al., 2021; Mager, 2006; Parsonnet, 1995).

Different from acute pathogens, *F. nucleatum* establishes persistent colonization in the oral cavity and can migrate to other sites via the hematogenous route, such as colorectal cancer (Abed et al., 2020). The persistence is facilitated by the adhesins of FadA to E-cadherin on epithelial cells, while Fap2 recognizes tumor-displayed Gal-GalNAc (Alon-Maimon et al., 2022).

Hemolysin activity adds to the cancer-promoting potential of *F. nucleatum*, but it alone is insufficient. The temporal and spatial context of bacterial-host interaction is important. While the proposed pathway of hemolysin induces ROS, acting as genotoxic, is biologically plausible and fits current biology, experimental validation is required to establish causality. The computational framework only provides testable predictions for future mechanistic studies.

### Tumor Microenvironment Compatibility

Cancer cells generally create a more acidic microenvironment than normal tissues, which favors their progression (Tafech and Stéphanou, 2024). Hemolysin secretion and pore formation are pH-dependent, with optimal activity near neutral pH (Kasschau et al., 1995).

Based on experimental literature, *F. nucleatum* demonstrates optimal growth at pH 7.4, a relatively neutral pH. The bacterium exhibits significantly reduced metabolic activity, with glucose consumption decreasing to less than 50% and cell yield being halved at pH 6.8. Under moderate acidic conditions (pH 5.8), growth is substantially reduced, while under slightly alkaline conditions (pH 7.8), growth fails (Rogers et al., 1991). While Rogers’s team (1991) found a narrow optimal range, other studies indicate a broader survival capacity. *F. nucleatum* subsp. polymorphum demonstrates the ability to form biofilms at alkaline pH 8.2, indicating subspecies-specific alkaline adaptation capabilities(Chew et al., 2012). However, *F. nucleatum* ATCC 25586 (subsp. nucleatum), the strain commonly found in oral cavities, probably exhibits tolerance to alkaline conditions, as *F. nucleatum* is found to survive in root canal systems at pH 9.0 after Ca(OH)_2_ treatment (Zilm and Rogers, 2007). Furthermore, a key literature finding suggests *F. nucleatum* possesses extreme acid tolerance, due to its cell membrane containing erucic acid, which is regulated by the FnFabM protein, giving it the ability to resist the extreme acidic conditions of the stomach (pH 1.5) when translocated into the intestine (Li et al., 2025).

*F. nucleatum* ability to modulate the tumor microenvironment(TME) can also promote proliferation, metastasis, and chemoresistance. In CRC tissue, iron deposits within the macrophages were correlated with a poor prognosis for the *F. nucleatum*-positive patients (Dadgar-Zankbar et al., 2024). Considering the proposed mechanism of iron dependency and potential cancer activity, *F. nucleatum*’s activity within the TME may be with respect to iron availability. It is predicted that iron concentration evaluation in the TME may be a signal for *F. nucleatum* to induce a phenotypic switch, enabling the upregulation of its Pathogenicity Island (PAI) genes, allowing *F. nucleatum* to exploit the TME and facilitate cancer progression. The upregulation of these genes involved with *F. nucleatum*’s virulence will, in turn, increase its adhesive and invasive behavior to the host tissues, immune modulation, and ultimately may work with the TME by providing an environment conducive to the growth of tumors (Yamane et al., 2022).

While this study focuses on strain ATCC 25586, future analysis can be done on PAI and the rest of the unannotated genes across *F. nucleatum* subspecies, using phylogenetic analyses and positive selection, which may illuminate whether evolutionary pressures are shaping these virulence-associated functions and potentially characterize unknown proteins by evolutionary relatedness (Charlotte West, 2025).

This study has several limitations. All pathogenicity island predictions, protein functions, and interaction networks are computational and require experimental validation. The AlphaFold-Multimer interface with predictions showed low confidence (ipTM *<* 0.5) should be interpreted cautiously. Our co-expression analysis was limited by sample size (n=18). This study focused on strain ATCC 25586, and findings may not generalize to other *F. nucleatum* strains. The proposed Hemolysin-Iron-ROS-Hippo pathway remains a hypothesis requiring experimental testing.

## Conclusion

This work show how integrating bioinformatics and machine learning approaches enables us to provide further details into the pathogenic mechanisms of bacteria. By combining genomic analysis, metabolic modeling, protein structure prediction, and transcriptomic data, we have constructed a coherent framework pinpointing potential pathogenicity islands of *F. nucleatum* and connect to a proposed mechanism of cancer promotion through iron-mediated oxidative stress.

Our multi-omics approach identified complete hemolysin and iron acquisition machinery within the pathogenicity islands, established iron as metabolically essential through flux balance analysis, and revealed host-induced upregulation of hemolysin expression. Contextualized within the broader literature on ROS-mediated Hippo pathway oncologically relevant and distinction between acute hemolytic pathogens and chronic colonizers, this finding proposes a biologically plausible model for *F. nuclea-tum*-associated carcinogenesis.

However, one should be conscious of the interpretive boundaries of this work. The result in this study is predictive and contextual rather than causal. In complex biological systems, functional redundancy and compensatory pathways often mean that no single component is strictly determinative of growth or virulence. The current results are state-level predictions rather than mechanistic claims. The study suggests configurations of genomic programs, metabolic constraints, and environmental conditions that are consistent with pathogenic behavior, rather than proving that any individual gene or island is causative on its own.

Furthermore, many of the predicted proteins and genomic regions remain speculative without direct experimental or transcriptomic validation. Rather than being a weakness, it is a clear guide for the next stage of investigation—identifying which predicted states are actually realized under specific host or environmental conditions, and how perturbations shift the system between them.

As the amount of molecular sequence data of pathogens in the public domain grows, so does the range of bioinformatics and machine learning methods that operate on them through functional, structural, and evolutionary considerations. The methodology presented here provides a template for systematic investigation of bacterial virulence factors and their potential roles in human disease. However, advancement of the field is hampered by the discrepancy between bioinformatics and deep learning for many of the analysis methods.

## Supporting information

Supplementary_README

## Notes

### Competing Interest Statement

The authors have declared no competing interest.

### Summary of Updates

I have updated the structure and interaction section by scanning through the whole PAI2, inside of only the 8 unknown proteins. Later, adding in coexpression analysis by using RNAseq database. I revised the figure in the paper to make it more standardized. I updated the conclusion and abstract accordingly. I upload my supplmentary materials which contain my coded.

