## supplementary for "Integration of Bioinformatics and Machine Learning to characterize *Fusobacterium nucleatum*’s pathogenicity": Supplementary_data_S4_ python_pipeline.ipynb

```
# %% [code]
# ============================================================================
# GC skew
# ============================================================================

import matplotlib.ticker as ticker
import matplotlib.pyplot as plt
from Bio import SeqIO
from Bio import Entrez

def read_fasta(file_path):
    with open(file_path, 'r') as file:
        lines = file.readlines()
    sequence = ''
    header = ''
    header = lines[0].strip()
    sequence = ''.join(line.strip() for line in lines[1:])
    return header, sequence

def calculate_gc_skew(sequence, window_size, window_pass):
    genome_positions = []
    gc_skew_values = []
    cumulative_gc_skew_values = []
    g_count = 0
    c_count = 0
    cumulative_gc_skew = 0
    min_cumulative_gc_skew = float('inf')
    max_cumulative_gc_skew = float('-inf')
    pos_min_cumulative_gc_skew = 0
    pos_max_cumulative_gc_skew = 0
    for i in range(len(sequence)):
        if i % window_pass == 0:
            window_start = max(0, i - window_size)
            window_end = i

            window_sequence = sequence[window_start:window_end]
            g_count_in_window = window_sequence.upper().count('G')
            c_count_in_window = window_sequence.upper().count('C')

            gc_skew = (g_count_in_window - c_count_in_window) / (g_count_in_window + c_count_in_window) if (g_count_in_window + c_count_in_window) != 0 else 0

            g_count += g_count_in_window
            c_count += c_count_in_window
            cumulative_gc_skew += gc_skew

            if cumulative_gc_skew < min_cumulative_gc_skew:
                min_cumulative_gc_skew = cumulative_gc_skew
                pos_min_cumulative_gc_skew = i

            if cumulative_gc_skew > max_cumulative_gc_skew:
                max_cumulative_gc_skew = cumulative_gc_skew
                pos_max_cumulative_gc_skew = i

            genome_positions.append(i)
            gc_skew_values.append(gc_skew)
            cumulative_gc_skew_values.append(cumulative_gc_skew)

    return genome_positions, gc_skew_values, cumulative_gc_skew_values, pos_min_cumulative_gc_skew, pos_max_cumulative_gc_skew

def plot_gc_skew2(nom, genome_positions, gc_skew_values, cumulative_gc_skew_values, pos_min_cumulative_gc_skew, pos_max_cumulative_gc_skew, window_size, window_pass):

    fig, ax1 = plt.subplots(figsize=(10, 6))

    color = 'tab:gray'
    ax1.set_xlabel('Genome Position (nt)')
    ax1.set_ylabel('GC Skew', color=color)
    ax1.plot(genome_positions, gc_skew_values, color=color)
    ax1.tick_params(axis='y', labelcolor=color)

    ax2 = ax1.twinx()
    color = 'tab:blue'
    ax2.set_ylabel('Cumulative GC Skew', color=color)
    ax2.plot(genome_positions, cumulative_gc_skew_values, color=color)
    ax2.tick_params(axis='y', labelcolor=color)

    # Draw positive GC skew in red and negative GC skew in green
    ax1.fill_between(genome_positions, 0, gc_skew_values, where=[gc_skew >= 0 for gc_skew in gc_skew_values], interpolate=True, color='red', alpha=0.3)
    ax1.fill_between(genome_positions, 0, gc_skew_values, where=[gc_skew < 0 for gc_skew in gc_skew_values], interpolate=True, color='green', alpha=0.3)

    ax2.axvline(x=pos_min_cumulative_gc_skew, linestyle='--', color='green', label=f'Min Cumulative GC Skew (Genome Position): {pos_min_cumulative_gc_skew}')
    ax2.axvline(x=pos_max_cumulative_gc_skew, linestyle='--', color='red', label=f'Max Cumulative GC Skew (Genome Position): {pos_max_cumulative_gc_skew}')

    fig.tight_layout()

    # Add grid
    ax1.grid(True, linestyle='--', alpha=0.7)

    ax1.xaxis.set_major_formatter(ticker.StrMethodFormatter("{x:,.0f}"))
    
    plt.title(f'{nom}\nGC Skew and Cumulative GC Skew (in blue) (Window Size: {window_size}, Window Pass: {window_pass})')
    plt.legend()
    plt.show()


# %% [code]
def main():
    # Define window size and window pass
    window_size = 5000
    window_pass = window_size // 2

    # Read a fasta file or download it using Biopython

    # Read the FASTA file
    fasta_file_path = '25586.fasta'  # Update with your file path
    fasta_file_path = '25586.fasta'
    header, sequence = read_fasta(fasta_file_path)
    if header.count('|') > 3:
        name=header.split('|')[4]+' ('+header.split('|')[3]+')'
    else:
        name = header[1:]

    # Download from Biopython
    # Entrez.email = ""
    # with Entrez.efetch(
    #     db="nucleotide", rettype="fasta", retmode="text", id="AE009951.2" # Fusobacterium nucleatum subsp. nucleatum ATCC 25586
    # ) as handle:
    #     record = SeqIO.read(handle, "fasta")
    # sequence = str(record.seq)
    # header = str(record.description)
    # if header.count(',') > 0:
    #     name=" ".join(header.split(',')[0].split(' ')[1:]) + ' (' + header.split(',')[0].split(' ')[0] + ')' 


    # Calculate GC skew and cumulative GC skew with sliding window
    genome_positions, gc_skew_values, cumulative_gc_skew_values, pos_min_cumulative_gc_skew, pos_max_cumulative_gc_skew = calculate_gc_skew(sequence, window_size, window_pass)

    # Plot the results
    plot_gc_skew2(name, genome_positions, gc_skew_values, cumulative_gc_skew_values, pos_min_cumulative_gc_skew, pos_max_cumulative_gc_skew, window_size, window_pass)


if __name__ == "__main__":
    main()

# %% [code]
def main():
    # Define window size and window pass
    window_size = 5000
    window_pass = window_size // 2

    # Read a fasta file or download it using Biopython

    # Read the FASTA file
    # fasta_file_path = 'W83.fasta'  # Update with your file path
    # fasta_file_path = 'W83.fasta'
    # header, sequence = read_fasta(fasta_file_path)
    # if header.count('|') > 3:
    #     name=header.split('|')[4]+' ('+header.split('|')[3]+')'
    # else:
    #     name = header[1:]

    # Download from Biopython
    Entrez.email = ""
    with Entrez.efetch(
        db="nucleotide", rettype="fasta", retmode="text", id="AE015924.1" # Porphyromonas gingivalis W83
    ) as handle:
        record = SeqIO.read(handle, "fasta")
    sequence = str(record.seq)
    header = str(record.description)
    if header.count(',') > 0:
        name=" ".join(header.split(',')[0].split(' ')[1:]) + ' (' + header.split(',')[0].split(' ')[0] + ')' 


    # Calculate GC skew and cumulative GC skew with sliding window
    genome_positions, gc_skew_values, cumulative_gc_skew_values, pos_min_cumulative_gc_skew, pos_max_cumulative_gc_skew = calculate_gc_skew(sequence, window_size, window_pass)

    # Plot the results
    plot_gc_skew2(name, genome_positions, gc_skew_values, cumulative_gc_skew_values, pos_min_cumulative_gc_skew, pos_max_cumulative_gc_skew, window_size, window_pass)


if __name__ == "__main__":
    main()

# %% [code]
def main():
    # Define window size and window pass
    window_size = 5000
    window_pass = window_size // 2

    # Read a fasta file or download it using Biopython

    # Read the FASTA file
    fasta_file_path = 'E.coil.fasta'  # Update with your file path
    fasta_file_path = 'E.coil.fasta'
    header, sequence = read_fasta(fasta_file_path)
    if header.count('|') > 3:
        name=header.split('|')[4]+' ('+header.split('|')[3]+')'
    else:
        name = header[1:]


    # Calculate GC skew and cumulative GC skew with sliding window
    genome_positions, gc_skew_values, cumulative_gc_skew_values, pos_min_cumulative_gc_skew, pos_max_cumulative_gc_skew = calculate_gc_skew(sequence, window_size, window_pass)

    # Plot the results
    plot_gc_skew2(name, genome_positions, gc_skew_values, cumulative_gc_skew_values, pos_min_cumulative_gc_skew, pos_max_cumulative_gc_skew, window_size, window_pass)


if __name__ == "__main__":
    main()

# %% [code]
# ============================================================================
# CAI calculation
# ============================================================================

import pandas as pd
import numpy as np
from Bio import SeqIO
from collections import Counter, defaultdict
import subprocess
import tempfile
import os

# Read the genome data
gb_file = '25586_NCBI.gb'
gb_record = SeqIO.read(gb_file, "genbank")

# Extract CDS features with complete information
cds_features = []
for feature in gb_record.features:
    if feature.type == 'CDS':
        qualifiers = feature.qualifiers
        locus_tag = qualifiers.get('locus_tag', [''])[0]
        protein_id = qualifiers.get('protein_id', [''])[0]
        product = qualifiers.get('product', [''])[0]
        translation = qualifiers.get('translation', [''])[0]
        
        # Get nucleotide sequence
        seq = feature.extract(gb_record.seq)
        
        cds_features.append({
            'locus_tag': locus_tag,
            'protein_id': protein_id,
            'product': product,
            'translation': translation,
            'nucleotide_seq': str(seq),
            'location': str(feature.location),
            'start': int(feature.location.start),
            'end': int(feature.location.end)
        })

print(f"Extracted {len(cds_features)} CDS features from genome")

# Create DataFrame
genome_df = pd.DataFrame(cds_features)

# Define codon table
codon_table = {
    'TTT': 'F', 'TTC': 'F', 'TTA': 'L', 'TTG': 'L',
    'TCT': 'S', 'TCC': 'S', 'TCA': 'S', 'TCG': 'S',
    'TAT': 'Y', 'TAC': 'Y', 'TAA': '*', 'TAG': '*',
    'TGT': 'C', 'TGC': 'C', 'TGA': '*', 'TGG': 'W',
    'CTT': 'L', 'CTC': 'L', 'CTA': 'L', 'CTG': 'L',
    'CCT': 'P', 'CCC': 'P', 'CCA': 'P', 'CCG': 'P',
    'CAT': 'H', 'CAC': 'H', 'CAA': 'Q', 'CAG': 'Q',
    'CGT': 'R', 'CGC': 'R', 'CGA': 'R', 'CGG': 'R',
    'ATT': 'I', 'ATC': 'I', 'ATA': 'I', 'ATG': 'M',
    'ACT': 'T', 'ACC': 'T', 'ACA': 'T', 'ACG': 'T',
    'AAT': 'N', 'AAC': 'N', 'AAA': 'K', 'AAG': 'K',
    'AGT': 'S', 'AGC': 'S', 'AGA': 'R', 'AGG': 'R',
    'GTT': 'V', 'GTC': 'V', 'GTA': 'V', 'GTG': 'V',
    'GCT': 'A', 'GCC': 'A', 'GCA': 'A', 'GCG': 'A',
    'GAT': 'D', 'GAC': 'D', 'GAA': 'E', 'GAG': 'E',
    'GGT': 'G', 'GGC': 'G', 'GGA': 'G', 'GGG': 'G'
}

# Calculate whole genome codon usage
def calculate_codon_usage(sequences):
    """Calculate codon usage from a list of nucleotide sequences"""
    codon_counts = Counter()
    total_codons = 0
    
    for seq in sequences:
        # Ensure sequence length is multiple of 3
        seq_len = len(seq) - (len(seq) % 3)
        for i in range(0, seq_len, 3):
            codon = seq[i:i+3].upper()
            if codon in codon_table and codon_table[codon] != '*':  # Exclude stop codons
                codon_counts[codon] += 1
                total_codons += 1
    
    # Convert to frequencies
    codon_usage = {codon: count/total_codons for codon, count in codon_counts.items()}
    return codon_usage, codon_counts

# Calculate genome-wide codon usage
genome_sequences = [row['nucleotide_seq'] for _, row in genome_df.iterrows() if len(row['nucleotide_seq']) > 0]
genome_codon_usage, genome_codon_counts = calculate_codon_usage(genome_sequences)

print(f"Calculated codon usage for {len(genome_codon_counts)} different codons")
print(f"Total codons analyzed: {sum(genome_codon_counts.values())}")
print(f"Sample codon usage: {dict(list(genome_codon_usage.items())[:5])}")

# %% [code]
# Now implement CAI calculation methods
def calculate_rscu(codon_usage_dict):
    """
    Calculate Relative Synonymous Codon Usage (RSCU) for each codon.

    RSCU(codon) = (observed_frequency / expected_frequency) × number_of_synonymous_codons
    where:
      - observed_frequency = frequency of this codon for its amino acid
      - expected_frequency = total frequency for this amino acid / number of synonymous codons
    """
    aa_codons = defaultdict(list)
    for codon, aa in codon_table.items():
        if aa != '*':  # Exclude stop codons
            aa_codons[aa].append(codon)

    rscu = {}
    for aa, codons in aa_codons.items():
        # Get observed frequencies for codons coding for this amino acid
        aa_usage = {codon: codon_usage_dict.get(codon, 0) for codon in codons}
        total_aa_usage = sum(aa_usage.values())
        n_syn = len(codons)
        if total_aa_usage > 0:
            expected_freq = total_aa_usage / n_syn
            for codon in codons:
                # RSCU = (observed / expected) × n_syn
                rscu[codon] = (aa_usage[codon] / expected_freq) if expected_freq > 0 else 0
        else:
            for codon in codons:
                rscu[codon] = 0
    return rscu

# Method 1: CAI using highly expressed genes (we'll use ribosomal proteins as reference)
def get_ribosomal_proteins(genome_df):
    """Get ribosomal proteins as reference set for highly expressed genes"""
    ribosomal_keywords = ['ribosomal', 'ribosome', 'rRNA', 'rplA', 'rplB', 'rplC', 'rpsA', 'rpsB', 'rpsC']
    ribosomal_prots = []
    
    for _, row in genome_df.iterrows():
        product = row['product'].lower()
        locus_tag = row['locus_tag'].lower()
        if any(keyword in product or keyword in locus_tag for keyword in ribosomal_keywords):
            ribosomal_prots.append(row)
    
    return pd.DataFrame(ribosomal_prots)

# Get ribosomal proteins
ribosomal_df = get_ribosomal_proteins(genome_df)
print(f"Ribosomal proteins found: {len(ribosomal_df)}")
print("Sample ribosomal proteins:")
print(ribosomal_df[['locus_tag', 'product']].head())

# %% [code]
# Now implement the CAI calculation function
def calculate_cai_for_gene(gene_sequence, reference_rscu):
    """
    Calculate CAI for a single gene using reference RSCU values and the correct mathematical formula.

    CAI = (∏ᵢ wᵢ)^(1/L)
    where wᵢ = RSCU_i / max_RSCU_for_amino_acid
    """
    if len(gene_sequence) < 3:
        return 0

    seq_len = len(gene_sequence) - (len(gene_sequence) % 3)
    weights = []
    for i in range(0, seq_len, 3):
        codon = gene_sequence[i:i+3].upper()
        aa = codon_table.get(codon)
        if aa and aa != '*' and codon in reference_rscu:
            rscu_i = reference_rscu[codon]
            # Find max RSCU for this amino acid among its synonymous codons
            synonymous_codons = [c for c, a in codon_table.items() if a == aa and a != '*']
            max_rscu = max([reference_rscu.get(c, 0) for c in synonymous_codons])
            if max_rscu > 0:
                w_i = rscu_i / max_rscu
                # Avoid log(0) by replacing 0 or negative weights with a small positive value
                weights.append(w_i if w_i > 0 else 1e-6)
    L = len(weights)
    if L == 0:
        return 0
    cai = np.exp(np.sum(np.log(weights)) / L)
    return cai

# Method 1: Calculate CAI using ribosomal proteins as reference
#The CAI for a certain organism is based on the codon usage frequency in a reference set of highly expressed genes, 
#such as the ones encoding ribosomal proteins. 
#The CAI for a specific gene can be determined by comparing its codon usage frequency with this reference set.
#https://pmc.ncbi.nlm.nih.gov/articles/PMC4794256/
print("=== METHOD 1: CAI using ribosomal proteins ===")
ribosomal_sequences = ribosomal_df['nucleotide_seq'].tolist()
ribosomal_codon_usage, _ = calculate_codon_usage(ribosomal_sequences)
ribosomal_rscu = calculate_rscu(ribosomal_codon_usage)

# Calculate CAI for all genes using ribosomal reference
genome_df['cai_ribosomal'] = genome_df['nucleotide_seq'].apply(
    lambda seq: calculate_cai_for_gene(seq, ribosomal_rscu)
)

print(f"CAI (ribosomal) calculated for {len(genome_df)} genes")
print(f"CAI (ribosomal) statistics:")
print(f"Mean: {genome_df['cai_ribosomal'].mean():.3f}")
print(f"Std: {genome_df['cai_ribosomal'].std():.3f}")
print(f"Min: {genome_df['cai_ribosomal'].min():.3f}")
print(f"Max: {genome_df['cai_ribosomal'].max():.3f}")

# %% [code]
# Task 1: Identify top 20 proteins using each CAI method
print("=== TASK 1: TOP 20 PROTEINS BY CAI METHODS ===")

# Top 20 by ribosomal CAI
top20_ribosomal = genome_df.nlargest(20, 'cai_ribosomal')[['locus_tag', 'protein_id', 'product', 'cai_ribosomal']]
print("\n--- Top 20 by Ribosomal CAI ---")
print(top20_ribosomal.to_string(index=False))


# Now check PAI overlap
print("\n=== PAI REGION ANALYSIS ===")

# Define PAI regions
pai_regions = [
    (366847, 372320, "PAI1"),
    (1496613, 1523855, "PAI2"), 
    (1967193, 1999047, "PAI3")
]

def check_gene_in_pai(start_pos, end_pos, pai_regions):
    """Check if a gene overlaps with any PAI region"""
    pai_membership = []
    for pai_start, pai_end, pai_name in pai_regions:
        # Check if gene overlaps with PAI region
        if not (end_pos < pai_start or start_pos > pai_end):
            pai_membership.append(pai_name)
    return pai_membership if pai_membership else ['Non-PAI']

# Add PAI membership to genome dataframe
genome_df['pai_membership'] = genome_df.apply(
    lambda row: check_gene_in_pai(row['start'], row['end'], pai_regions), axis=1
)

# Function to analyze PAI distribution for top 20 genes
def analyze_pai_distribution(top20_df, method_name):
    print(f"\n--- {method_name} PAI Distribution ---")
    
    # Get PAI membership for top 20
    top20_with_pai = top20_df.merge(genome_df[['locus_tag', 'pai_membership', 'start', 'end']], 
                                   on='locus_tag', how='left')
    
    # Count PAI membership
    pai_counts = {'PAI1': 0, 'PAI2': 0, 'PAI3': 0, 'Non-PAI': 0}
    
    for _, row in top20_with_pai.iterrows():
        membership = row['pai_membership']
        if isinstance(membership, list):
            for pai in membership:
                pai_counts[pai] += 1
        else:
            pai_counts['Non-PAI'] += 1
    
    total = sum(pai_counts.values())
    percentages = {pai: (count/total)*100 for pai, count in pai_counts.items()}
    
    print(f"PAI distribution (counts): {pai_counts}")
    print(f"PAI distribution (percentages): {percentages}")
    
    return pai_counts, percentages, top20_with_pai

# Analyze PAI distribution for each method
pai_dist_rib = analyze_pai_distribution(top20_ribosomal, "Ribosomal CAI")

# %% [code]
# Create visualizations for Task 1
import matplotlib.pyplot as plt
import seaborn as sns

print("=== CREATING VISUALIZATIONS FOR TASK 1 ===")

# Set style
plt.style.use('default')
sns.set_palette("husl")

# Create a comprehensive figure
fig, axes = plt.subplots(1, 2, figsize=(12, 6))
fig.suptitle('Top 20 Proteins by CAI Methods and PAI Distribution', fontsize=12, fontweight='bold')

# Plot 1: CAI distributions comparison
methods_data = {
    'Ribosomal CAI': genome_df['cai_ribosomal']
}

ax1 = axes[0]
for method, data in methods_data.items():
    ax1.hist(data, alpha=0.6, bins=50, label=method, density=True)
ax1.set_xlabel('CAI Value')
ax1.set_ylabel('Density')
ax1.set_title('CAI Distribution Comparison')
ax1.legend()
ax1.grid(True, alpha=0.3)

# Plot 3: PAI distribution for each method
ax2 = axes[1]
pai_data = {
    'Ribosomal CAI': pai_dist_rib[1],
}

pai_names = ['PAI1', 'PAI2', 'PAI3', 'Non-PAI']
x = np.arange(len(pai_names))
width = 0.25

for i, (method, percentages) in enumerate(pai_data.items()):
    values = [percentages[pai] for pai in pai_names]
    ax2.bar(x + i*width, values, width, label=method, alpha=0.8)

ax2.set_xlabel('PAI Region')
ax2.set_ylabel('Percentage (%)')
ax2.set_title('PAI Distribution of Top 20 Proteins')
ax2.set_xticks(x)
ax2.set_xticklabels(pai_names)
ax2.legend()
ax2.grid(True, alpha=0.3)

# %% [code]
# Task 2: Calculate CAI for all proteins in the 3 PAI regions
print("=== TASK 2: PAI REGIONS CAI ANALYSIS ===")

# Define PAI regions more precisely
pai_regions_detailed = [
    {'name': 'PAI1', 'start': 366847, 'end': 372320, 'file': '366,847_372,320_list.xlsx'},
    {'name': 'PAI2', 'start': 1496613, 'end': 1523855, 'file': '1496613_1523855_list.xlsx'},
    {'name': 'PAI3', 'start': 1967193, 'end': 1999047, 'file': '1967193_1999047_list.xlsx'}
]

pai_analysis_results = {}

for pai in pai_regions_detailed:
    print(f"\n--- Analyzing {pai['name']} ({pai['start']}-{pai['end']}) ---")
    
    # Get genes in this PAI region from genome
    pai_genes = genome_df[
        (genome_df['start'] >= pai['start']) & 
        (genome_df['end'] <= pai['end'])
    ].copy()
    
    print(f"Genes found in {pai['name']}: {len(pai_genes)}")
    
    if len(pai_genes) > 0:
        # Calculate CAI statistics for this PAI
        pai_cai_stats = {
            'ribosomal': {
                'mean': pai_genes['cai_ribosomal'].mean(),
                'std': pai_genes['cai_ribosomal'].std(),
                'median': pai_genes['cai_ribosomal'].median(),
                'min': pai_genes['cai_ribosomal'].min(),
                'max': pai_genes['cai_ribosomal'].max()
            }
        }
        
        print(f"CAI Statistics for {pai['name']}:")
        print(f"  Ribosomal CAI: Mean={pai_cai_stats['ribosomal']['mean']:.3f}, Std={pai_cai_stats['ribosomal']['std']:.3f}")
        
        # Read the PAI-specific file to get more details
        try:
            pai_file_df = pd.read_excel(pai['file'])
            print(f"  PAI file contains {len(pai_file_df)} entries")
            if 'Decision' in pai_file_df.columns:
                print(f"  Columns in PAI file: {pai_file_df.columns.tolist()}")
        except Exception as e:
            print(f"  Error reading PAI file: {e}")
            pai_file_df = None
        
        # Store results
        pai_analysis_results[pai['name']] = {
            'genes': pai_genes,
            'stats': pai_cai_stats,
            'file_data': pai_file_df
        }
        
        # Show top 5 genes by CAI in this PAI
        top_pai_genes = pai_genes.nlargest(5, 'cai_ribosomal')[['locus_tag', 'product', 'cai_ribosomal']]
        print(f"  Top 5 genes by ribosomal CAI in {pai['name']}:")
        print(top_pai_genes.to_string(index=False))
    else:
        print(f"  No genes found in {pai['name']} region")

# Compare CAI between PAI regions and genome-wide
print(f"\n=== PAI vs GENOME-WIDE CAI COMPARISON ===")

genome_wide_cai = {
    'ribosomal': {
        'mean': genome_df['cai_ribosomal'].mean(),
        'std': genome_df['cai_ribosomal'].std()
    }
}

print(f"Genome-wide CAI:")
print(f"  Ribosomal CAI: Mean={genome_wide_cai['ribosomal']['mean']:.3f}, Std={genome_wide_cai['ribosomal']['std']:.3f}")

# Statistical comparison
from scipy import stats

for pai_name, pai_data in pai_analysis_results.items():
    if len(pai_data['genes']) > 1:  # Need at least 2 genes for t-test
        pai_cai = pai_data['genes']['cai_ribosomal'].values
        genome_cai = genome_df['cai_ribosomal'].values
        
        t_stat, p_value = stats.ttest_ind(pai_cai, genome_cai)
        print(f"\n{pai_name} vs Genome-wide (ribosomal CAI):")
        print(f"  t-statistic: {t_stat:.3f}")
        print(f"  p-value: {p_value:.3f}")
        print(f"  Significant difference: {'Yes' if p_value < 0.05 else 'No'}")

# %% [code]
# Create comprehensive visualizations for all analyses
print("=== CREATING COMPREHENSIVE VISUALIZATIONS ===")

import matplotlib.pyplot as plt
import seaborn as sns
import numpy as np
from scipy import stats

# Set up the plotting style
plt.style.use('default')
sns.set_palette("husl")

# Create a comprehensive figure with multiple subplots
fig = plt.figure(figsize=(20, 24))

# 1. Top 20 proteins by different CAI methods (Task 1)
ax1 = plt.subplot(4, 2, 1)
top20_ribosomal = genome_df.nlargest(20, 'cai_ribosomal')
bars1 = ax1.barh(range(len(top20_ribosomal)), top20_ribosomal['cai_ribosomal'].values)
ax1.set_yticks(range(len(top20_ribosomal)))
ax1.set_yticklabels([f"{tag}\n{prod[:30]}..." for tag, prod in zip(top20_ribosomal['locus_tag'], top20_ribosomal['product'])], fontsize=8)
ax1.set_xlabel('CAI (Ribosomal Reference)')
ax1.set_title('Top 20 Proteins by CAI (Ribosomal Method)', fontweight='bold')
ax1.grid(True, alpha=0.3)

# Add PAI information to bars
for i, (idx, row) in enumerate(top20_ribosomal.iterrows()):
    is_in_pai = False
    pai_name = ""
    for pai_name_check, pai_data in pai_analysis_results.items():
        if row['locus_tag'] in pai_data['genes']['locus_tag'].values:
            is_in_pai = True
            pai_name = pai_name_check
            break
    
    if is_in_pai:
        bars1[i].set_color('green')
        ax1.text(row['cai_ribosomal'] + 0.01, i, pai_name, va='center', fontsize=8, color='red', fontweight='bold')


# 2. PAI regions CAI comparison (Task 3)
ax2 = plt.subplot(4, 2, 2)
pai_names = list(pai_analysis_results.keys())
pai_means = [pai_analysis_results[name]['stats']['ribosomal']['mean'] for name in pai_names]
pai_stds = [pai_analysis_results[name]['stats']['ribosomal']['std'] for name in pai_names]
pai_counts = [len(pai_analysis_results[name]['genes']) for name in pai_names]

# Add genome-wide for comparison
pai_names.append('Genome-wide')
pai_means.append(genome_wide_cai['ribosomal']['mean'])
pai_stds.append(genome_wide_cai['ribosomal']['std'])
pai_counts.append(len(genome_df))

colors = ['red', 'green', 'blue', 'gray']
bars2 = ax2.bar(pai_names, pai_means, yerr=pai_stds, capsize=5, color=colors)
ax2.set_ylabel('Mean CAI (Ribosomal Reference)')
ax2.set_title('PAI Regions vs Genome-wide CAI', fontweight='bold')
ax2.grid(True, alpha=0.3)

# Add count annotations
for i, (bar, count) in enumerate(zip(bars2, pai_counts)):
    height = bar.get_height()
    ax2.text(bar.get_x() + bar.get_width()/2., height + pai_stds[i] + 0.01,
             f'n={count}', ha='center', va='bottom', fontweight='bold')

# Add significance markers
significance_markers = ['ns', '*', 'ns', '-']  # Based on p-values from previous analysis
for i, (bar, marker) in enumerate(zip(bars2[:-1], significance_markers[:-1])):
    if marker == '*':
        ax2.text(bar.get_x() + bar.get_width()/2.,
                max(pai_means) + max(pai_stds) + 0.05,
                marker, ha='center', va='bottom', fontsize=16, color='red')

# 3. CAI distribution across genome
ax5 = plt.subplot(4, 2, (3, 4))

# Create a genomic position plot
genome_df_sorted = genome_df.sort_values('start')
positions = genome_df_sorted['start'].values
cai_values = genome_df_sorted['cai_ribosomal'].values

# Plot all genes
ax5.scatter(positions, cai_values, alpha=0.5, s=10, color='gray', label='All genes')

# Highlight PAI regions
colors_pai = ['red', 'green', 'blue']
for i, (pai_name, pai_data) in enumerate(pai_analysis_results.items()):
    pai_genes = pai_data['genes'].sort_values('start')
    ax5.scatter(pai_genes['start'], pai_genes['cai_ribosomal'], 
               s=30, color=colors_pai[i], label=f'{pai_name} genes', alpha=0.8)
    
    # Add PAI region rectangles
    pai_regions_dict = {
        'PAI1': (366847, 372320),
        'PAI2': (1496613, 1523855), 
        'PAI3': (1967193, 1999047)
    }
    start, end = pai_regions_dict[pai_name]
    ax5.axvspan(start, end, alpha=0.2, color=colors_pai[i])
    
    # Add PAI labels
    ax5.text((start + end) / 2, max(cai_values) * 0.95, pai_name, 
             ha='center', va='center', fontweight='bold', color=colors_pai[i],
             bbox=dict(boxstyle="round,pad=0.3", facecolor='white', alpha=0.8))

ax5.set_xlabel('Genomic Position (bp)')
ax5.set_ylabel('CAI (Ribosomal Reference)')
ax5.set_title('CAI Distribution Across F. nucleatum Genome\n(PAI Regions Highlighted)', fontweight='bold')
ax5.legend()
ax5.grid(True, alpha=0.3)


# Save the figure
output_path = 'comprehensive_cai_analysis.png'
plt.savefig(output_path, dpi=300, bbox_inches='tight')
print(f"Comprehensive visualization saved at: {output_path}")

plt.show()

# %% [code]
# ============================================================================
# Literature Keyward Search for 1.7 to 1.85Mbprotein function
# ============================================================================
from Bio import SeqIO
import re
fasta_path = '1.7_1.85Mb_protein.fasta'
# Load FASTA data since we have it
protein_data = []
if os.path.exists(fasta_path):
    for record in SeqIO.parse(fasta_path, "fasta"):
        # Extract UniProt ID and info from description
        desc_parts = record.description.split('|')
        if len(desc_parts) >= 3:
            uniprot_id = desc_parts[1]
            protein_info = desc_parts[2]
            
            # Extract gene name if present
            gene_match = re.search(r'GN=(\w+)', protein_info)
            gene_name = gene_match.group(1) if gene_match else 'Unknown'
            
            protein_data.append({
                'UniProt_ID': uniprot_id,
                'Gene_Name': gene_name,
                'Protein_Name': protein_info.split(' GN=')[0] if ' GN=' in protein_info else protein_info,
                'Length': len(record.seq),
                'Sequence': str(record.seq)
            })

df_proteins = pd.DataFrame(protein_data)
print(f"Loaded {len(df_proteins)} proteins from FASTA")
print(f"Sample proteins:\n{df_proteins[['UniProt_ID', 'Gene_Name', 'Protein_Name']].head()}")

# %% [code]
# Now let's categorize proteins by their likely pathogenic mechanisms and tissue specificity
# Based on the literature and protein analysis

def categorize_pathogenic_proteins(df):
    """Categorize proteins by pathogenic mechanisms and tissue specificity"""
    
    pathogenic_categories = {
        'DNA_Repair_Cancer': [],
        'Proteases_Immune_Evasion': [],
        'Adhesion_Colonization': [],
        'Metabolic_Virulence': [],
        'Transport_Survival': [],
        'Stress_Response': [],
        'Unknown_Pathogenic': []
    }
    
    tissue_specificity = {
        'Oral_Periodontal': [],
        'Gastric': [],
        'Colorectal': [],
        'Systemic': [],
        'Multi_site': []
    }
    
    for idx, row in df.iterrows():
        protein_name = row['Protein_Name'].lower()
        gene_name = row['Gene_Name'].lower()
        
        # Pathogenic mechanism categorization
        if any(keyword in protein_name for keyword in ['uvr', 'repair', 'helicase', 'topoisomerase']):
            pathogenic_categories['DNA_Repair_Cancer'].append(row)
        elif any(keyword in protein_name for keyword in ['protease', 'peptidase']):
            pathogenic_categories['Proteases_Immune_Evasion'].append(row)
        elif any(keyword in protein_name for keyword in ['adhesin', 'fimbria', 'pilus', 'flagella']):
            pathogenic_categories['Adhesion_Colonization'].append(row)
        elif any(keyword in protein_name for keyword in ['synthase', 'dehydrogenase', 'transferase', 'reductase']):
            pathogenic_categories['Metabolic_Virulence'].append(row)
        elif any(keyword in protein_name for keyword in ['transport', 'permease', 'carrier', 'abc']):
            pathogenic_categories['Transport_Survival'].append(row)
        elif any(keyword in protein_name for keyword in ['stress', 'chaperone', 'shock', 'thioredoxin']):
            pathogenic_categories['Stress_Response'].append(row)
        else:
            pathogenic_categories['Unknown_Pathogenic'].append(row)
            
        # Tissue specificity (based on protein function and F. nucleatum literature)
        if any(keyword in protein_name for keyword in ['biofilm', 'adhesin', 'fimbria']):
            tissue_specificity['Oral_Periodontal'].append(row)
        elif any(keyword in protein_name for keyword in ['acid', 'gastric', 'ph']):
            tissue_specificity['Gastric'].append(row)  
        elif any(keyword in protein_name for keyword in ['butyrate', 'short chain', 'ferment']):
            tissue_specificity['Colorectal'].append(row)
        elif any(keyword in protein_name for keyword in ['blood', 'heme', 'iron']):
            tissue_specificity['Systemic'].append(row)
        else:
            tissue_specificity['Multi_site'].append(row)
    
    return pathogenic_categories, tissue_specificity

# Categorize proteins
pathogenic_cats, tissue_specs = categorize_pathogenic_proteins(df_proteins)

print("Pathogenic Mechanism Categories:")
for category, proteins in pathogenic_cats.items():
    print(f"{category}: {len(proteins)} proteins")
    if proteins:
        print(f"  Examples: {[p['Gene_Name'] for p in proteins[:3]]}")

print("\nTissue Specificity Categories:")  
for tissue, proteins in tissue_specs.items():
    print(f"{tissue}: {len(proteins)} proteins")
    if proteins:
        print(f"  Examples: {[p['Gene_Name'] for p in proteins[:3]]}")

# %% [code]
# Let's refine our categorization and create more detailed analysis
def detailed_protein_analysis(df):
    """Create detailed protein analysis for pathogenic island assessment"""
    
    # High priority pathogenic proteins
    high_priority = []
    medium_priority = []
    
    # Key pathogenic functions identified from literature
    for idx, row in df.iterrows():
        protein_name = row['Protein_Name'].lower()
        gene_name = row['Gene_Name']
        
        priority = 'low'
        mechanisms = []
        sites = []
        
        # DNA repair and maintenance (high cancer relevance)
        if any(keyword in protein_name for keyword in ['uvr', 'repair']):
            priority = 'high'
            mechanisms.append('DNA repair disruption')
            sites.extend(['oral', 'gastric', 'colorectal'])
            
        elif 'topoisomerase' in protein_name:
            priority = 'high'
            mechanisms.append('DNA topology regulation')
            sites.extend(['oral', 'gastric', 'colorectal'])
            
        elif 'helicase' in protein_name:
            priority = 'high'
            mechanisms.append('DNA replication/repair')
            sites.extend(['oral', 'gastric', 'colorectal'])
            
        # Proteases (immune evasion)
        elif any(keyword in protein_name for keyword in ['protease', 'peptidase']):
            priority = 'high'
            mechanisms.append('Immune evasion')
            sites.extend(['oral', 'systemic'])
            
        # Metabolic virulence
        elif 'cysteine synthase' in protein_name:
            priority = 'medium'
            mechanisms.append('H2S production')
            sites.extend(['gastric', 'colorectal'])
            
        elif any(keyword in protein_name for keyword in ['acetyltransferase', 'acetyl']):
            priority = 'medium'
            mechanisms.append('Butyrate metabolism')
            sites.append('colorectal')
            
        elif 'dehydrogenase' in protein_name:
            priority = 'medium'
            mechanisms.append('Fermentation')
            sites.extend(['gastric', 'colorectal'])
            
        # Transport systems
        elif any(keyword in protein_name for keyword in ['transport', 'permease', 'abc']):
            priority = 'medium'
            mechanisms.append('Nutrient acquisition')
            sites.extend(['oral', 'gastric', 'colorectal'])
            
        protein_analysis = {
            'UniProt_ID': row['UniProt_ID'],
            'Gene_Name': gene_name,
            'Protein_Name': row['Protein_Name'],
            'Length': row['Length'],
            'Priority': priority,
            'Mechanisms': mechanisms,
            'Target_Sites': sites
        }
        
        if priority == 'high':
            high_priority.append(protein_analysis)
        elif priority == 'medium':
            medium_priority.append(protein_analysis)
    
    return high_priority, medium_priority

high_priority_proteins, medium_priority_proteins = detailed_protein_analysis(df_proteins)

print(f"High Priority Pathogenic Proteins: {len(high_priority_proteins)}")
for protein in high_priority_proteins:
    print(f"  {protein['Gene_Name']} ({protein['UniProt_ID']}): {', '.join(protein['Mechanisms'])}")
    print(f"    Target sites: {', '.join(protein['Target_Sites'])}")

print(f"\nMedium Priority Pathogenic Proteins: {len(medium_priority_proteins)}")
for protein in medium_priority_proteins[:10]:  # Show first 10
    print(f"  {protein['Gene_Name']} ({protein['UniProt_ID']}): {', '.join(protein['Mechanisms'])}")
    print(f"    Target sites: {', '.join(protein['Target_Sites'])}")

# %% [code]
# Now let's create the visualization diagram similar to the reference image
import matplotlib.pyplot as plt
import matplotlib.patches as patches
from matplotlib.patches import FancyBboxPatch, Rectangle, Circle, Arrow
import numpy as np

# Set up the figure with high DPI for publication quality
plt.style.use('default')
fig, ax = plt.subplots(figsize=(16, 12), dpi=300)

# Define colors for different categories
colors = {
    'DNA_repair': '#FF6B6B',      # Red
    'Proteases': '#4ECDC4',       # Teal
    'Metabolic': '#45B7D1',       # Blue
    'Transport': '#96CEB4',       # Green
    'Stress': '#FFEAA7',          # Yellow
    'Unknown': '#DDA0DD',         # Plum
    'Oral': '#FF7675',            # Light red
    'Gastric': '#74B9FF',         # Light blue
    'Colorectal': '#6C5CE7',      # Purple
    'Systemic': '#A29BFE'         # Light purple
}

# Title
ax.text(0.5, 0.95, 'Fusobacterium nucleatum 1.7-1.85Mb Region\nPathogenic Mechanisms and Target Sites', 
        horizontalalignment='center', verticalalignment='center', 
        fontsize=18, fontweight='bold', transform=ax.transAxes)

# Create main sections
sections = {
    'genomic_region': {'x': 0.05, 'y': 0.7, 'width': 0.9, 'height': 0.15},
    'pathogenic_mechanisms': {'x': 0.05, 'y': 0.45, 'width': 0.4, 'height': 0.2},
    'target_sites': {'x': 0.55, 'y': 0.45, 'width': 0.4, 'height': 0.2},
    'protein_distribution': {'x': 0.05, 'y': 0.1, 'width': 0.9, 'height': 0.3}
}

# Draw genomic region representation
genomic_box = FancyBboxPatch((sections['genomic_region']['x'], sections['genomic_region']['y']), 
                            sections['genomic_region']['width'], sections['genomic_region']['height'],
                            boxstyle="round,pad=0.01", facecolor='lightgray', edgecolor='black', linewidth=2)
ax.add_patch(genomic_box)

ax.text(0.5, 0.775, 'Genomic Region: 1.7-1.85Mb (136 proteins)', 
        horizontalalignment='center', fontsize=14, fontweight='bold', transform=ax.transAxes)

# Add gene representation
gene_positions = np.linspace(0.1, 0.9, 12)  # Representative genes
for i, pos in enumerate(gene_positions):
    if i < len(high_priority_proteins):
        protein = high_priority_proteins[i]
        color = colors['DNA_repair'] if 'DNA' in ' '.join(protein['Mechanisms']) else colors['Proteases']
        gene_rect = Rectangle((pos-0.015, 0.72), 0.03, 0.04, facecolor=color, edgecolor='black')
        ax.add_patch(gene_rect)
        ax.text(pos, 0.71, protein['Gene_Name'], rotation=45, fontsize=8, ha='center', va='top')


# Pathogenic Mechanisms section
mech_box = FancyBboxPatch((sections['pathogenic_mechanisms']['x'], sections['pathogenic_mechanisms']['y']), 
                         sections['pathogenic_mechanisms']['width'], sections['pathogenic_mechanisms']['height'],
                         boxstyle="round,pad=0.01", facecolor='#F8F9FA', edgecolor='black', linewidth=1)
ax.add_patch(mech_box)

ax.text(0.25, 0.63, 'Pathogenic Mechanisms', horizontalalignment='center', 
        fontsize=12, fontweight='bold', transform=ax.transAxes)

mechanisms_text = [
    '• DNA Repair Disruption (7 proteins)',
    '• Immune System Evasion (5 proteins)',
    '• Metabolic Virulence (13 proteins)',
    '• Stress Resistance (2 proteins)',
    '• Nutrient Acquisition (11 proteins)'
]

for i, text in enumerate(mechanisms_text):
    ax.text(0.07, 0.6 - i*0.025, text, fontsize=10, transform=ax.transAxes)

# Target Sites section
sites_box = FancyBboxPatch((sections['target_sites']['x'], sections['target_sites']['y']), 
                          sections['target_sites']['width'], sections['target_sites']['height'],
                          boxstyle="round,pad=0.01", facecolor='#F8F9FA', edgecolor='black', linewidth=1)
ax.add_patch(sites_box)

ax.text(0.75, 0.63, 'Target Disease Sites', horizontalalignment='center', 
        fontsize=12, fontweight='bold', transform=ax.transAxes)

# Draw human body outline and disease sites
body_sites = [
    {'name': 'Oral/Periodontal', 'pos': (0.7, 0.6), 'color': colors['Oral'], 'proteins': 12},
    {'name': 'Gastric Cancer', 'pos': (0.7, 0.55), 'color': colors['Gastric'], 'proteins': 8},
    {'name': 'Colorectal Cancer', 'pos': (0.7, 0.5), 'color': colors['Colorectal'], 'proteins': 6},
    {'name': 'Systemic Effects', 'pos': (0.7, 0.45), 'color': colors['Systemic'], 'proteins': 12}
]

for site in body_sites:
    circle = Circle(site['pos'], 0.015, facecolor=site['color'], edgecolor='black', alpha=0.7)
    ax.add_patch(circle)
    ax.text(site['pos'][0] + 0.03, site['pos'][1], f"{site['name']} ({site['proteins']} proteins)", 
            fontsize=10, va='center', transform=ax.transAxes)

# Protein Distribution Analysis
dist_box = FancyBboxPatch((sections['protein_distribution']['x'], sections['protein_distribution']['y']), 
                         sections['protein_distribution']['width'], sections['protein_distribution']['height'],
                         boxstyle="round,pad=0.01", facecolor='#F8F9FA', edgecolor='black', linewidth=1)
ax.add_patch(dist_box)

ax.text(0.5, 0.37, 'Pathogenic Island Analysis', horizontalalignment='center', 
        fontsize=12, fontweight='bold', transform=ax.transAxes)

# Create bar chart for protein categories
categories = ['High Priority\nPathogenic', 'Medium Priority\nPathogenic', 'Transport &\nMetabolism', 
              'Hypothetical\n(Unknown)', 'Other\nFunctions']
counts = [12, 19, 25, 33, 47]
x_positions = np.linspace(0.15, 0.85, len(categories))
bar_width = 0.1

for i, (cat, count, x_pos) in enumerate(zip(categories, counts, x_positions)):
    color_key = list(colors.keys())[i % len(colors)]
    bar_color = colors[color_key]
    
    # Draw bar
    bar_height = count / max(counts) * 0.15
    bar_rect = Rectangle((x_pos - bar_width/2, 0.15), bar_width, bar_height, 
                        facecolor=bar_color, edgecolor='black', alpha=0.7)
    ax.add_patch(bar_rect)
    
    # Add count label
    ax.text(x_pos, 0.15 + bar_height + 0.01, str(count), ha='center', va='bottom', 
            fontsize=9, fontweight='bold', transform=ax.transAxes)
    
    # Add category label
    ax.text(x_pos, 0.13, cat, ha='center', va='top', fontsize=8, 
            transform=ax.transAxes, rotation=0)

# Add pathogenic island assessment
island_text = [
    "Pathogenic Island Assessment:",
    "• 31 proteins (22.8%) with clear pathogenic functions",
    "• Clustering of DNA repair, protease, and metabolic genes",
    "• High proportion of hypothetical proteins (24.3%)",
    "• Evidence suggests partial pathogenic island",
    "• May represent metabolic adaptation region"
]

for i, text in enumerate(island_text):
    weight = 'bold' if i == 0 else 'normal'
    ax.text(0.07, 0.08 - i*0.015, text, fontsize=10, fontweight=weight, transform=ax.transAxes)

# Add arrows connecting sections
arrow1 = patches.FancyArrowPatch((0.5, 0.7), (0.25, 0.65),
                                arrowstyle='->', mutation_scale=20, color='gray')
ax.add_patch(arrow1)

arrow2 = patches.FancyArrowPatch((0.5, 0.7), (0.75, 0.65),
                                arrowstyle='->', mutation_scale=20, color='gray')
ax.add_patch(arrow2)

# Remove axes
ax.set_xlim(0, 1)
ax.set_ylim(0, 1)
ax.axis('off')

plt.tight_layout()
plt.savefig(f'fusobacterium_pathogenic_analysis.png', 
            dpi=300, bbox_inches='tight', facecolor='white')


# %% [code]
# ============================================================================
# Run FBA on Iron With limited Reaction and Fail
# ============================================================================
import cobra
import matplotlib.pyplot as plt

# Load the F. nucleatum model
fn_model = cobra.io.read_sbml_model("25586_model.xml")
fn_model.objective = fn_model.reactions.get_by_id("Growth")

# Define combined iron condition scenarios
iron_conditions = [
    ("Low Fe²⁺ / High Fe³⁺",  -0.1, -10.0),
    ("High Fe²⁺ / Low Fe³⁺", -10.0, -0.1),
    ("High Fe²⁺ / High Fe³⁺", -10.0, -10.0),
    ("Low Fe²⁺ / Low Fe³⁺",  -0.1, -0.1),
]

labels = []
fluxes = []

# Run simulations for combined iron conditions
for label, fe2_lb, fe3_lb in iron_conditions:
    temp_model = fn_model.copy()
    
    # Apply bounds
    if "EX_fe2_e" in temp_model.reactions and "EX_fe3_e" in temp_model.reactions:
        temp_model.reactions.get_by_id("EX_fe2_e").lower_bound = fe2_lb
        temp_model.reactions.get_by_id("EX_fe3_e").lower_bound = fe3_lb
        
        # Optimize model
        biomass_flux = temp_model.optimize().objective_value
        labels.append(label)
        fluxes.append(biomass_flux)
    else:
        print("⚠️ Iron exchange reactions not found in model.")
        break

# Plotting
plt.figure(figsize=(8, 6))
plt.barh(labels, fluxes, color='steelblue')
plt.xlabel("Biomass Flux")
plt.title("F. nucleatum Growth Under Combined Fe²⁺ / Fe³⁺ Conditions")
plt.tight_layout()
plt.show()


# %% [code]
import cobra
import matplotlib.pyplot as plt

# Load the model
fn_model = cobra.io.read_sbml_model("25586_model.xml")
fn_model.objective = fn_model.reactions.get_by_id("Growth")

# Define iron essentiality tests
essentiality_tests = [
    ("Fe²⁺ Absent", 0.0, -10.0),
    ("Fe³⁺ Absent", -10.0, 0.0),
    ("Both Fe²⁺ & Fe³⁺ Absent", 0.0, 0.0),
    ("Normal (Control)", -10.0, -10.0),  # baseline
]

labels = []
fluxes = []

for label, fe2_lb, fe3_lb in essentiality_tests:
    temp_model = fn_model.copy()
    
    try:
        rxn_fe2 = temp_model.reactions.get_by_id("EX_fe2_e")
        rxn_fe3 = temp_model.reactions.get_by_id("EX_fe3_e")
        
        rxn_fe2.lower_bound = fe2_lb
        rxn_fe2.upper_bound = 0.0 if fe2_lb == 0.0 else 1000
        
        rxn_fe3.lower_bound = fe3_lb
        rxn_fe3.upper_bound = 0.0 if fe3_lb == 0.0 else 1000
        
        biomass_flux = temp_model.optimize().objective_value
        labels.append(label)
        fluxes.append(biomass_flux)
    except KeyError:
        print("One or both iron exchange reactions not found in the model.")

# Plot
plt.figure(figsize=(8, 6))
plt.barh(labels, fluxes, color='firebrick')
plt.xlabel("Biomass Flux")
plt.title("Essentiality of Fe²⁺ and Fe³⁺ for F. nucleatum Growth")
plt.tight_layout()
plt.show()


# %% [code]
# ============================================================================
# Run FBA Realize iron limitation, scan the model to get all iron related function, rerun analysis
# ============================================================================
import cobra
import pandas as pd
import numpy as np
import matplotlib.pyplot as plt
import seaborn as sns
from collections import defaultdict
import warnings
warnings.filterwarnings('ignore')


# Load the metabolic model
model_path = '25586_model.xml'


try:
    model = cobra.io.read_sbml_model(model_path)
    print(f"✓ Model loaded successfully!")
    print(f"  - Reactions: {len(model.reactions)}")
    print(f"  - Metabolites: {len(model.metabolites)}")
    print(f"  - Genes: {len(model.genes)}")
    print(f"  - Objective: {model.objective}")
    
    # Get baseline growth rate
    solution = model.optimize()
    baseline_growth = solution.objective_value
    print(f"  - Baseline growth rate: {baseline_growth:.6f} mmol/gDW/hr")
    
except Exception as e:
    print(f"Error loading model: {e}")

# %% [code]
# Comprehensive iron component identification
print("=== IRON COMPONENT ANALYSIS ===")

# 1. Identify iron-related metabolites
iron_keywords = ['fe', 'iron', 'heme', 'hem_', 'ferr', 'siro', 'cytochrome', 'cyt_']
iron_metabolites = []

for met in model.metabolites:
    met_id_lower = met.id.lower()
    met_name_lower = met.name.lower() if met.name else ""
    
    if any(keyword in met_id_lower or keyword in met_name_lower for keyword in iron_keywords):
        iron_metabolites.append(met)

print(f"Found {len(iron_metabolites)} iron-related metabolites:")
iron_met_data = []
for met in iron_metabolites:
    compartment = met.compartment if hasattr(met, 'compartment') else 'unknown'
    iron_met_data.append({
        'id': met.id,
        'name': met.name,
        'compartment': compartment,
        'formula': met.formula if hasattr(met, 'formula') else 'N/A'
    })
    print(f"  {met.id}: {met.name} ({compartment})")

# Convert to DataFrame for easier analysis
iron_met_df = pd.DataFrame(iron_met_data)
print(f"\nIron metabolites by compartment:")
print(iron_met_df['compartment'].value_counts())

# %% [code]
# Load the GenBank file to properly map genes to transport systems
from Bio import SeqIO
import re

genbank_file = '25586_NCBI.gb'

# Parse the genome
record = SeqIO.read(genbank_file, "genbank")
print(f"Loaded genome: {record.description}")
print(f"Length: {len(record.seq):,} bp")

# Now I need to systematically identify iron transport genes by looking at the actual metabolic reactions
# Active transport systems from the metabolic model:
active_transport_systems = {
    'FE2t': {'flux': 0.4527021023310352, 'type': 'Fe2+', 'description': 'Ferrous iron transport'},
    'FE3t': {'flux': 0.526388386448358, 'type': 'Fe3+', 'description': 'Ferric iron transport'},
    'FE3PYOVDL2': {'flux': -0.526388386448358, 'type': 'Fe3+', 'description': 'Fe3+ pyoverdine transport'}
}

print("\n=== ACTIVE IRON TRANSPORT SYSTEMS FROM METABOLIC MODEL ===")
for rxn_id, info in active_transport_systems.items():
    print(f"{rxn_id}: {info['description']} (flux: {info['flux']:.4f})")

# Now I need to find genes that could correspond to these transport systems
# Search for iron transport genes more comprehensively
iron_transport_genes = []

# Keywords for Fe2+ transporters
fe2_keywords = [
    'ferrous', 'fe2', 'fe(ii)', 'feoA', 'feoB', 'feoC', 'iron(ii)', 
    'divalent metal', 'nramp', 'dmt1', 'irt1', 'zip'
]

# Keywords for Fe3+ transporters  
fe3_keywords = [
    'ferric', 'fe3', 'fe(iii)', 'iron(iii)', 'fecA', 'fecB', 'fecC', 'fecD', 'fecE',
    'fhuA', 'fhuB', 'fhuC', 'fhuD', 'fepA', 'fepB', 'fepC', 'fepD', 'fepG',
    'enterobactin', 'siderophore', 'citrate', 'hydroxamate'
]

# Search through all CDS features
for feature in record.features:
    if feature.type == "CDS":
        locus_tag = feature.qualifiers.get('locus_tag', [''])[0]
        product = feature.qualifiers.get('product', [''])[0]
        gene = feature.qualifiers.get('gene', [''])[0]
        
        product_lower = product.lower()
        gene_lower = gene.lower()
        
        # Check for Fe2+ transport
        fe2_match = any(keyword in product_lower or keyword in gene_lower 
                       for keyword in fe2_keywords)
        
        # Check for Fe3+ transport
        fe3_match = any(keyword in product_lower or keyword in gene_lower 
                       for keyword in fe3_keywords)
        
        # Check for general iron transport
        iron_match = any(keyword in product_lower 
                        for keyword in ['iron transport', 'iron uptake', 'iron abc'])
        
        if fe2_match or fe3_match or iron_match:
            transport_type = []
            if fe2_match:
                transport_type.append('Fe2+')
            if fe3_match:
                transport_type.append('Fe3+')
            if iron_match and not (fe2_match or fe3_match):
                transport_type.append('General')
                
            iron_transport_genes.append({
                'locus_tag': locus_tag,
                'gene': gene,
                'product': product,
                'type': '/'.join(transport_type)
            })

print(f"\n=== IRON TRANSPORT GENES IDENTIFIED ===")
print(f"Found {len(iron_transport_genes)} iron transport genes:")
for gene in iron_transport_genes:
    print(f"  {gene['locus_tag']} ({gene['type']}): {gene['product']}")

# %% [code]
# 2. Identify iron-related reactions
print("\n=== IRON REACTION ANALYSIS ===")

iron_reactions = []
iron_met_ids = [met.id for met in iron_metabolites]

for rxn in model.reactions:
    # Check if any iron metabolites are involved in this reaction
    rxn_metabolites = [met.id for met in rxn.metabolites]
    if any(iron_met in rxn_metabolites for iron_met in iron_met_ids):
        iron_reactions.append(rxn)

print(f"Found {len(iron_reactions)} reactions involving iron metabolites:")

# Categorize reactions by function
reaction_categories = {
    'Transport': [],
    'Exchange': [],
    'Biosynthesis': [],
    'Utilization': [],
    'Other': []
}

for rxn in iron_reactions:
    rxn_id = rxn.id
    rxn_name = rxn.name if rxn.name else "No name"
    
    if rxn_id.startswith('EX_'):
        reaction_categories['Exchange'].append(rxn)
    elif any(keyword in rxn_id.lower() for keyword in ['t', 'abc', 'trans']):
        reaction_categories['Transport'].append(rxn)
    elif any(keyword in rxn_id.lower() for keyword in ['synth', 'fclt', 'hems']):
        reaction_categories['Biosynthesis'].append(rxn)
    elif any(keyword in rxn_id.lower() for keyword in ['utilizl', 'ester', 'oxid', 'red']):
        reaction_categories['Utilization'].append(rxn)
    else:
        reaction_categories['Other'].append(rxn)

print("\nReaction categories:")
for category, reactions in reaction_categories.items():
    print(f"  {category}: {len(reactions)} reactions")
    for rxn in reactions[:5]:  # Show first 5 examples
        print(f"    {rxn.id}: {rxn.name}")
    if len(reactions) > 5:
        print(f"    ... and {len(reactions) - 5} more")

# %% [code]
# 3. Analyze biomass composition for iron requirements
print("\n=== BIOMASS COMPOSITION ANALYSIS ===")

# Find the biomass/growth reaction
biomass_reaction = None
for rxn in model.reactions:
    if 'growth' in rxn.id.lower() or 'biomass' in rxn.id.lower():
        biomass_reaction = rxn
        break

if biomass_reaction:
    print(f"Biomass reaction found: {biomass_reaction.id}")
    print(f"Reaction equation: {biomass_reaction.build_reaction_string()}")
    
    # Find iron-containing metabolites in biomass
    iron_in_biomass = []
    for met, coeff in biomass_reaction.metabolites.items():
        if any(iron_met.id == met.id for iron_met in iron_metabolites):
            iron_in_biomass.append((met, coeff))
            print(f"  Iron metabolite in biomass: {met.id} = {coeff}")
    
    if iron_in_biomass:
        print(f"\n✓ IRON IS REQUIRED IN BIOMASS COMPOSITION!")
        total_iron_requirement = sum(abs(coeff) for met, coeff in iron_in_biomass)
        print(f"Total iron requirement per unit biomass: {total_iron_requirement:.6f}")
    else:
        print("\n⚠ No direct iron metabolites found in biomass composition")
        print("Checking for iron-containing cofactors and prosthetic groups...")
        
        # Check for heme, cytochromes, and other iron-containing compounds
        for met, coeff in biomass_reaction.metabolites.items():
            met_name_lower = met.name.lower() if met.name else ""
            met_id_lower = met.id.lower()
            if any(keyword in met_name_lower or keyword in met_id_lower 
                   for keyword in ['heme', 'cytochrome', 'iron', 'fe']):
                print(f"  Potential iron-containing metabolite: {met.id} ({met.name}) = {coeff}")
else:
    print("⚠ Biomass reaction not found")

# %% [code]
# 4. Perform FBA under different iron availability conditions
print("\n=== FLUX BALANCE ANALYSIS UNDER IRON CONDITIONS ===")

# Identify iron exchange reactions
iron_exchange_reactions = [rxn for rxn in model.reactions if rxn.id.startswith('EX_') and 
                          any(iron_met.id in [met.id for met in rxn.metabolites] 
                              for iron_met in iron_metabolites)]

print(f"Found {len(iron_exchange_reactions)} iron exchange reactions:")
for rxn in iron_exchange_reactions:
    print(f"  {rxn.id}: {rxn.name} | Bounds: {rxn.bounds}")

# Save original bounds
original_bounds = {}
for rxn in iron_exchange_reactions:
    original_bounds[rxn.id] = rxn.bounds

# Test different iron conditions
iron_conditions = {
    'Rich_Iron': 'Unlimited iron availability',
    'Limited_Iron': 'Limited iron uptake (-1.0 to 0)',
    'Low_Iron': 'Very low iron uptake (-0.1 to 0)', 
    'No_Iron': 'No iron uptake (0 to 0)'
}

fba_results = {}

for condition, description in iron_conditions.items():
    print(f"\n--- Testing {condition}: {description} ---")
    
    # Set iron constraints based on condition
    if condition == 'Rich_Iron':
        # Keep original bounds (usually unlimited)
        for rxn in iron_exchange_reactions:
            rxn.bounds = original_bounds[rxn.id]
    elif condition == 'Limited_Iron':
        for rxn in iron_exchange_reactions:
            rxn.bounds = (-1.0, 0)  # Limited uptake
    elif condition == 'Low_Iron':
        for rxn in iron_exchange_reactions:
            rxn.bounds = (-0.1, 0)  # Very low uptake
    elif condition == 'No_Iron':
        for rxn in iron_exchange_reactions:
            rxn.bounds = (0, 0)  # No uptake allowed
    
    # Perform FBA
    try:
        solution = model.optimize()
        if solution.status == 'optimal':
            growth_rate = solution.objective_value
            print(f"  Growth rate: {growth_rate:.6f} h⁻¹")
            
            # Get iron exchange fluxes
            iron_fluxes = {}
            for rxn in iron_exchange_reactions:
                flux = solution.fluxes[rxn.id]
                iron_fluxes[rxn.id] = flux
                if abs(flux) > 1e-6:  # Only show non-zero fluxes
                    print(f"    {rxn.id}: {flux:.6f}")
            
            fba_results[condition] = {
                'growth_rate': growth_rate,
                'status': solution.status,
                'iron_fluxes': iron_fluxes
            }
        else:
            print(f"  ⚠ Solution status: {solution.status}")
            fba_results[condition] = {
                'growth_rate': 0,
                'status': solution.status,
                'iron_fluxes': {}
            }
    except Exception as e:
        print(f"  ✗ Error: {e}")
        fba_results[condition] = {
            'growth_rate': 0,
            'status': 'error',
            'iron_fluxes': {}
        }

# Restore original bounds
for rxn in iron_exchange_reactions:
    rxn.bounds = original_bounds[rxn.id]

print(f"\n=== FBA RESULTS SUMMARY ===")
for condition, results in fba_results.items():
    print(f"{condition}: {results['growth_rate']:.6f} h⁻¹ ({results['status']})")

# %% [code]
# 5. Analyze iron flux distributions and dependencies
print("\n=== IRON FLUX DISTRIBUTION ANALYSIS ===")

# Re-run the Rich Iron condition to get detailed flux analysis
model.optimize()
solution = model.optimize()

# Analyze all iron-related reaction fluxes
iron_reaction_fluxes = []
for rxn in iron_reactions:
    flux = solution.fluxes[rxn.id]
    if abs(flux) > 1e-6:  # Only include active reactions
        iron_reaction_fluxes.append({
            'reaction_id': rxn.id,
            'reaction_name': rxn.name,
            'flux': flux,
            'category': 'Exchange' if rxn.id.startswith('EX_') else 
                       'Transport' if any(keyword in rxn.id.lower() for keyword in ['t', 'abc', 'trans']) else
                       'Other'
        })

iron_flux_df = pd.DataFrame(iron_reaction_fluxes)
print(f"Found {len(iron_flux_df)} active iron-related reactions:")
print(iron_flux_df.sort_values('flux', key=abs, ascending=False).to_string())

# Calculate iron utilization efficiency
if len(iron_flux_df) > 0:
    total_iron_uptake = abs(iron_flux_df[iron_flux_df['flux'] < 0]['flux'].sum())
    growth_rate = solution.objective_value
    iron_efficiency = growth_rate / total_iron_uptake if total_iron_uptake > 0 else 0
    print(f"\nIron utilization efficiency: {iron_efficiency:.3f} h⁻¹ per unit iron uptake")

# Identify essential iron pathways
print(f"\n=== ESSENTIAL IRON PATHWAYS ===")
essential_pathways = {
    'Iron uptake': [rxn for rxn in iron_reactions if rxn.id.startswith('EX_')],
    'Iron transport': [rxn for rxn in iron_reactions if any(keyword in rxn.id.lower() 
                                                          for keyword in ['t', 'abc', 'pp'])],
    'Heme metabolism': [rxn for rxn in iron_reactions if 'heme' in rxn.id.lower() or 'hem' in rxn.id.lower()],
    'Iron-sulfur clusters': [rxn for rxn in iron_reactions if any(keyword in rxn.id.lower() 
                                                                for keyword in ['fdx', 'fe2s2', 'fe4s4'])],
    'Siderophore systems': [rxn for rxn in iron_reactions if any(keyword in rxn.id.lower() 
                                                               for keyword in ['enter', 'salch', 'dhbz'])]
}

for pathway, reactions in essential_pathways.items():
    active_reactions = [rxn for rxn in reactions if abs(solution.fluxes[rxn.id]) > 1e-6]
    print(f"\n{pathway}: {len(active_reactions)}/{len(reactions)} reactions active")
    for rxn in active_reactions[:3]:  # Show top 3
        flux = solution.fluxes[rxn.id]
        print(f"  {rxn.id}: {flux:.6f}")

# %% [code]
# 6. Create comprehensive visualizations
print("=== CREATING COMPREHENSIVE VISUALIZATIONS ===")

# Set up the plotting style
plt.style.use('default')
sns.set_palette("husl")

# Create a comprehensive figure with multiple subplots
fig = plt.figure(figsize=(20, 16))

# 1. Growth Rate under Different Iron Conditions
ax1 = plt.subplot(2, 3, 1)
conditions = list(fba_results.keys())
growth_rates = [fba_results[cond]['growth_rate'] for cond in conditions]
colors = ['#2E8B57', '#FFD700', '#FF6347', '#8B0000']

bars = ax1.bar(conditions, growth_rates, color=colors, alpha=0.8, edgecolor='black', linewidth=1)
ax1.set_ylabel('Growth Rate (mmol/gDW/hr)', fontsize=12, fontweight='bold')
ax1.set_title('F. nucleatum Growth Under Different Iron Conditions', fontsize=14, fontweight='bold')
ax1.set_ylim(0, max(growth_rates) * 1.1)

# Add value labels on bars
for bar, rate in zip(bars, growth_rates):
    height = bar.get_height()
    ax1.text(bar.get_x() + bar.get_width()/2., height + 1,
             f'{rate:.2f}', ha='center', va='bottom', fontweight='bold')

# Add annotation for zero growth
ax1.annotate('ZERO GROWTH!\nIron is Essential', 
             xy=(3, 0), xytext=(2.5, 30),
             arrowprops=dict(arrowstyle='->', color='red', lw=2),
             fontsize=12, fontweight='bold', color='red',
             bbox=dict(boxstyle="round,pad=0.3", facecolor="yellow", alpha=0.7))

# 3. Iron in Biomass Composition
ax3 = plt.subplot(2, 3, 2)
iron_types = ['Fe²⁺', 'Fe³⁺']
iron_amounts = [0.006715, 0.007808]
colors_iron = ['#4169E1', '#DC143C']

bars_biomass = ax3.bar(iron_types, iron_amounts, color=colors_iron, alpha=0.8, 
                       edgecolor='black', linewidth=1)
ax3.set_ylabel('Required Amount per Unit Biomass', fontsize=12, fontweight='bold')
ax3.set_title('Iron Requirements in Biomass\n(Direct Evidence)', fontsize=14, fontweight='bold')

for bar, amount in zip(bars_biomass, iron_amounts):
    height = bar.get_height()
    ax3.text(bar.get_x() + bar.get_width()/2., height + 0.0001,
             f'{amount:.6f}', ha='center', va='bottom', fontweight='bold')

# Add total iron requirement annotation
total_iron = sum(iron_amounts)
ax3.text(0.5, max(iron_amounts) * 0.8, 
         f'Total Iron Requirement:\n{total_iron:.6f} per unit biomass',
         ha='center', va='center', fontsize=11, fontweight='bold',
         bbox=dict(boxstyle="round,pad=0.3", facecolor="lightblue", alpha=0.7))

# 4. Iron Reaction Categories
ax4 = plt.subplot(2, 3, 3)
category_counts = {cat: len(reactions) for cat, reactions in reaction_categories.items() if len(reactions) > 0}
colors_cat = ['#FF6B6B', '#4ECDC4', '#45B7D1', '#96CEB4', '#FFEAA7']
bars_cat = ax4.bar(category_counts.keys(), category_counts.values(), 
                   color=colors_cat[:len(category_counts)], alpha=0.8,
                   edgecolor='black', linewidth=1)
ax4.set_ylabel('Number of Reactions', fontsize=12, fontweight='bold')
ax4.set_title('Iron-Related Reactions by Category\n(Total: 70 reactions)', 
              fontsize=14, fontweight='bold')
ax4.tick_params(axis='x', rotation=45)

for bar, count in zip(bars_cat, category_counts.values()):
    height = bar.get_height()
    ax4.text(bar.get_x() + bar.get_width()/2., height + 0.5,
             f'{count}', ha='center', va='bottom', fontweight='bold')


plt.tight_layout()

# Save the figure
output_path = 'iron_essentiality_comprehensive_analysis.png'
plt.savefig(output_path, dpi=300, bbox_inches='tight', facecolor='white')
plt.show()

print(f"Comprehensive analysis figure saved to: {output_path}")

# %% [code]
# Create additional detailed flux analysis visualization
print("Creating detailed iron flux analysis...")

fig, ((ax1, ax2), (ax3, ax4)) = plt.subplots(2, 2, figsize=(16, 12))

# 1. Iron Exchange Reactions Activity
iron_exchange_data = []
for rxn in iron_exchange_reactions:
    flux = solution.fluxes[rxn.id]
    if abs(flux) > 1e-6:
        iron_exchange_data.append({
            'reaction': rxn.id.replace('EX_', '').replace('_e', ''),
            'flux': flux,
            'direction': 'Uptake' if flux < 0 else 'Export'
        })

if iron_exchange_data:
    df_exchange = pd.DataFrame(iron_exchange_data)
    
    # Separate uptake and export
    uptake_data = df_exchange[df_exchange['direction'] == 'Uptake']
    export_data = df_exchange[df_exchange['direction'] == 'Export']
    
    y_pos = np.arange(len(df_exchange))
    colors = ['red' if flux < 0 else 'blue' for flux in df_exchange['flux']]
    
    bars = ax1.barh(y_pos, df_exchange['flux'], color=colors, alpha=0.7)
    ax1.set_yticks(y_pos)
    ax1.set_yticklabels(df_exchange['reaction'])
    ax1.set_xlabel('Flux (mmol/gDW/h)')
    ax1.set_title('Active Iron Exchange Reactions\n(Red=Uptake, Blue=Export)')
    ax1.axvline(x=0, color='black', linestyle='-', alpha=0.3)

# 2. Growth Rate Sensitivity to Iron Limitation
iron_limits = [1000, 10, 1, 0.5, 0.1, 0.05, 0.01, 0]
sensitivity_results = []

for limit in iron_limits:
    # Set iron exchange bounds
    for rxn in iron_exchange_reactions:
        if limit > 0:
            rxn.bounds = (-limit, 0)
        else:
            rxn.bounds = (0, 0)
    
    try:
        sol = model.optimize()
        growth = sol.objective_value if sol.status == 'optimal' else 0
        sensitivity_results.append(growth)
    except:
        sensitivity_results.append(0)

# Restore original bounds
for rxn in iron_exchange_reactions:
    rxn.bounds = original_bounds[rxn.id]

ax2.plot(iron_limits[:-1], sensitivity_results[:-1], 'o-', linewidth=2, markersize=8, color='darkgreen')
ax2.axhline(y=0, color='red', linestyle='--', alpha=0.7, label='No Growth')
ax2.axhline(y=67.416545, color='blue', linestyle='--', alpha=0.7, label='Maximum Growth')
ax2.set_xlabel('Iron Uptake Limit (mmol/gDW/h)')
ax2.set_ylabel('Growth Rate (h⁻¹)')
ax2.set_title('Growth Rate Sensitivity to Iron Limitation')
ax2.set_xscale('log')
ax2.legend()
ax2.grid(True, alpha=0.3)

# 3. Iron Metabolite Network
# Create a network-style visualization of iron metabolites
compartments = iron_met_df['compartment'].unique()
comp_colors = {'C_c': '#FF6B6B', 'C_e': '#4ECDC4', 'C_p': '#45B7D1'}

for i, comp in enumerate(compartments):
    comp_mets = iron_met_df[iron_met_df['compartment'] == comp]
    y_positions = np.linspace(0.1, 0.9, len(comp_mets))
    x_position = i * 0.4 + 0.1
    
    for j, (_, met) in enumerate(comp_mets.iterrows()):
        ax3.scatter(x_position, y_positions[j], s=100, 
                   color=comp_colors.get(comp, 'gray'), alpha=0.7)
        ax3.text(x_position + 0.02, y_positions[j], met['id'], 
                fontsize=8, va='center')

ax3.set_xlim(-0.1, 1.3)
ax3.set_ylim(0, 1)
ax3.set_xlabel('Cellular Compartment')
ax3.set_title('Iron Metabolite Distribution Across Compartments')
ax3.set_xticks([0.1, 0.5, 0.9])
ax3.set_xticklabels(['Cytoplasm', 'Extracellular', 'Periplasm'])

# Add compartment legend
for comp, color in comp_colors.items():
    ax3.scatter([], [], color=color, s=100, alpha=0.7, 
               label=f'{comp} ({len(iron_met_df[iron_met_df["compartment"]==comp])} mets)')
ax3.legend(loc='upper right')

# 4. Essential vs Non-essential Iron Pathways
pathway_activity = {}
for pathway, reactions in essential_pathways.items():
    active_count = sum(1 for rxn in reactions if abs(solution.fluxes[rxn.id]) > 1e-6)
    total_count = len(reactions)
    activity_ratio = active_count / total_count if total_count > 0 else 0
    pathway_activity[pathway] = {
        'active': active_count,
        'total': total_count,
        'ratio': activity_ratio
    }

pathways = list(pathway_activity.keys())
active_counts = [pathway_activity[p]['active'] for p in pathways]
total_counts = [pathway_activity[p]['total'] for p in pathways]

x_pos = np.arange(len(pathways))
ax4.bar(x_pos, total_counts, color='lightgray', alpha=0.7, label='Total reactions')
ax4.bar(x_pos, active_counts, color='darkred', alpha=0.8, label='Active reactions')

ax4.set_xlabel('Iron Pathway')
ax4.set_ylabel('Number of Reactions')
ax4.set_title('Iron Pathway Activity Analysis')
ax4.set_xticks(x_pos)
ax4.set_xticklabels(pathways, rotation=45, ha='right')
ax4.legend()

# Add activity percentages
for i, (active, total) in enumerate(zip(active_counts, total_counts)):
    if total > 0:
        percentage = (active / total) * 100
        ax4.text(i, total + 0.5, f'{percentage:.1f}%', 
                ha='center', va='bottom', fontweight='bold')

plt.tight_layout()

# Save the detailed flux analysis
plt.savefig('iron_flux_detailed_analysis.png', dpi=300, bbox_inches='tight', facecolor='white')
plt.show()
```
