## supplementary for "Integration of Bioinformatics and Machine Learning to characterize *Fusobacterium nucleatum*’s pathogenicity": Supplementary_data_S5_ bnlearn_analysis.R

```
# ============================================================================
# BAYESIAN NETWORK ANALYSIS OF PAI2 PROTEIN INTERACTIONS
# Using bnlearn R Package
# Data Source: GSE161360 (GEO) - F. nucleatum ATCC 25586 RNA-seq
#
# Bayesian network structure learning to identify co-expression relationships among 32 proteins in the PAI2 genomic island.
#
# ============================================================================

# ----------------------------------------------------------------------------
# STEP 1: INSTALL AND LOAD REQUIRED PACKAGES
# ----------------------------------------------------------------------------

# Install if not already installed, delete the "#" in front of the code to run
# if (!require("bnlearn")) {
#     install.packages("bnlearn", repos = "https://cloud.r-project.org/")
# }

# library(bnlearn)

# cat("bnlearn version:", as.character(packageVersion("bnlearn")), "\n")

# ----------------------------------------------------------------------------
# STEP 2: LOAD AND PREPARE EXPRESSION DATA
# ----------------------------------------------------------------------------
# The expression data was preprocessed from GSE161360:
# - WIG coverage files were parsed to extract per-gene coverage
# - Coverage was averaged across each gene's coordinates
# - Log2 transformation was applied (with pseudocount of 1)
#
# Data structure:
# - Rows: 18 samples (3 growth phases × 2 treatments × 3 replicates)
# - Columns: 32 PAI2 genes (FN0834-FN0865)

expr_data <- read.csv("pai2_expression_log2.csv", row.names = 1)

cat("\nData dimensions:\n")
cat("  Samples:", nrow(expr_data), "\n")
cat("  Genes:", ncol(expr_data), "\n")

# Convert to data frame (required by bnlearn)
expr_df <- as.data.frame(expr_data)

# Check data summary
cat("\nExpression value summary:\n")
print(summary(as.vector(as.matrix(expr_df))))

# ----------------------------------------------------------------------------
# STEP 3: BAYESIAN NETWORK STRUCTURE LEARNING
# ----------------------------------------------------------------------------
# We use the Hill-Climbing (hc) algorithm with BIC score.
#
# WHY HILL-CLIMBING?
# - Score-based algorithm that searches for the DAG maximizing a score
# - Efficient for moderate-sized networks
# - Works well with continuous (Gaussian) data
#
# WHY BIC SCORE?
# - Bayesian Information Criterion penalizes model complexity
# - Helps prevent overfitting, especially important with small sample sizes
# - "bic-g" is the Gaussian version for continuous data
#
# The algorithm:
# 1. Starts with an empty network (no edges)
# 2. Iteratively adds, removes, or reverses edges
# 3. Accepts changes that improve the BIC score
# 4. Stops when no improvement is possible

set.seed(42)  # For reproducibility

cat("\nLearning network structure...\n")
cat("  Algorithm: Hill-Climbing (hc)\n")
cat("  Score: BIC-Gaussian (bic-g)\n")

bn_structure <- hc(expr_df, score = "bic-g")

cat("\nNetwork learned!\n")
cat("  Nodes:", length(nodes(bn_structure)), "\n")
cat("  Edges:", narcs(bn_structure), "\n")

# View the learned edges
cat("\nLearned edges (directed):\n")
print(arcs(bn_structure))

# ----------------------------------------------------------------------------
# STEP 4: BOOTSTRAP ANALYSIS FOR EDGE CONFIDENCE
# ----------------------------------------------------------------------------
# A single network may be unstable due to sampling variability.
# Bootstrap analysis assesses edge robustness:
#
# 1. Resample the data with replacement (500 times)
# 2. Learn a network from each bootstrap sample
# 3. Count how often each edge appears
#
# Output metrics:
# - STRENGTH: Proportion of bootstrap samples containing this edge
#   (higher = more robust, range 0-1)
# - DIRECTION: Proportion where edge has this specific direction
#   (higher = more confident about causality direction)
#
# Significance thresholds (commonly used):
# - strength > 0.5: Edge appears in majority of bootstrap samples
# - direction > 0.5: Direction is consistent in majority of samples

cat("\nRunning bootstrap analysis (500 iterations)...\n")
cat("This may take a few minutes...\n")

set.seed(42)
boot_strength <- boot.strength(
    expr_df, 
    R = 500,                              # Number of bootstrap samples
    algorithm = "hc",                      # Same algorithm as above
    algorithm.args = list(score = "bic-g") # Same score
)

cat("Bootstrap complete!\n")

# Filter for significant edges
significant_edges <- boot_strength[
    boot_strength$strength > 0.5 & boot_strength$direction > 0.5, 
]
significant_edges <- significant_edges[order(-significant_edges$strength), ]

cat("\nSignificant edges (strength > 0.5, direction > 0.5):\n")
cat("Found", nrow(significant_edges), "significant edges\n\n")
print(head(significant_edges, 20))

# ----------------------------------------------------------------------------
# STEP 5: SAVE RESULTS
# ----------------------------------------------------------------------------

# Save all bootstrap edge strengths
write.csv(boot_strength, "bnlearn_all_edges.csv", row.names = FALSE)

# Save significant edges only
write.csv(significant_edges, "bnlearn_significant_edges.csv", row.names = FALSE)

cat("\nResults saved:\n")
cat("  - bnlearn_all_edges.csv (all pairwise edge strengths)\n")
cat("  - bnlearn_significant_edges.csv (significant edges only)\n")

# ----------------------------------------------------------------------------
# STEP 6: INTERPRETATION GUIDE
# ----------------------------------------------------------------------------
cat("\n")
cat("============================================================================\n")
cat("INTERPRETATION GUIDE\n")
cat("============================================================================\n")
cat("\n")
cat("WHAT THE EDGES MEAN:\n")
cat("- An edge A -> B suggests that expression of gene A is predictive of\n")
cat("  expression of gene B, given all other genes in the network.\n")
cat("- This could indicate: regulatory relationship, co-regulation, or\n")
cat("  membership in the same functional pathway.\n")
cat("\n")
cat("IMPORTANT CAVEATS:\n")
cat("1. CORRELATION ≠ CAUSATION: Edges represent statistical dependencies,\n")
cat("   not necessarily direct physical interactions or causal relationships.\n")
cat("\n")
cat("2. SAMPLE SIZE: With only 18 samples, the network may miss true edges\n")
cat("   (false negatives) or include spurious edges (false positives).\n")
cat("   Bootstrap analysis helps, but cannot fully compensate.\n")
cat("\n")
cat("3. DIRECTION: Edge directions should be interpreted cautiously.\n")
cat("   They reflect conditional dependencies, not necessarily causal flow.\n")
cat("\n")
cat("4. COMPLEMENTARY EVIDENCE: Edges supported by BOTH bnlearn AND STRING\n")
cat("   are more reliable, as they have both expression and database support.\n")
cat("\n")
cat("============================================================================\n")

# ----------------------------------------------------------------------------
# OPTIONAL: VISUALIZE THE NETWORK
# ----------------------------------------------------------------------------
# Uncomment to create a network plot (requires Rgraphviz from Bioconductor)
#
# if (require("Rgraphviz")) {
#     # Create averaged network from bootstrap
#     avg_net <- averaged.network(boot_strength, threshold = 0.5)
#     
#     # Plot
#     pdf("bnlearn_network.pdf", width = 12, height = 10)
#     graphviz.plot(avg_net, main = "PAI2 Bayesian Network (bootstrap threshold = 0.5)")
#     dev.off()
# }

cat("\nAnalysis complete!\n")
```
