## supplementary for "Integration of Bioinformatics and Machine Learning to characterize *Fusobacterium nucleatum*’s pathogenicity": Supplementary_data_S5_ bnlearn_analysis_README.md

# Bayesian Network Analysis of PAI2 Protein Interactions

[![DOI](10.18637/jss.v035.i03.) (Nagarajan R, Scutari M (2013). Bayesian Networks in R with Applications in Systems Biology. Springer, New York. doi:10.1007/978-1-4614-6446-4, ISBN 978-1-4614-6445-7, 978-1-4614-6446-4.)

## Overview

This repository contains the code and data processing pipeline for Bayesian network analysis of protein-protein interactions within the PAI2 genomic island of *Fusobacterium nucleatum* ATCC 25586. The analysis integrates RNA-seq expression data with computational interaction predictions to identify co-expression relationships among 32 PAI2-encoded proteins.

## Citation

If you use this code, please cite:

> DOI: https://doi.org/10.1101/2025.08.21.671586

>Several scripts and functions in this project were generated with the help of AI tools

## Requirements

### Software Dependencies

| Software | Version | Purpose |
|----------|---------|---------|
| Python | ≥3.8 | Data preprocessing |
| R | ≥4.0 | Bayesian network learning |
| bnlearn | ≥4.7 | R package for network analysis |

### Python Packages

```
numpy>=1.20.0
pandas>=1.3.0
```

### R Packages

```r
bnlearn (≥4.7)
```

## Data Source

Expression data was obtained from NCBI GEO:

- **Accession**: [GSE161360](https://www.ncbi.nlm.nih.gov/geo/query/acc.cgi?acc=GSE161360)
- **Title**: RNA landscape of the emerging cancer-bug Fusobacterium nucleatum
- **Reference**: Ponath F, et al. (2021) *Nature Communications*. PMID: 34239075
- **Organism**: *Fusobacterium nucleatum* subsp. nucleatum ATCC 25586
- **Samples**: 18 RNA-seq samples (3 growth phases × 2 treatments × 3 replicates)

## Repository Structure

```
├── README.md                          # This file
├── convert_wig_to_expression.ipynb       # Python script for data preprocessing
├── bnlearn_analysis_annotated.R       # R script for Bayesian network analysis
├── data/
│   ├── pai2_gene_coordinates.csv      # Gene coordinates (FN0834-FN0865)
│   ├── pai2_expression_log2.csv       # Preprocessed expression matrix
│   └── sample_metadata.csv            # Sample annotations
└── results/
    ├── bnlearn_all_edges.csv          # All pairwise edge strengths
    ├── bnlearn_significant_edges.csv  # Significant edges (filtered)
    └── bnlearn_string_overlap.csv     # Edges overlapping with STRING
```

## Quick Start

### Step 1: Download Raw Data from GEO

Go to https://www.ncbi.nlm.nih.gov/geo/query/acc.cgi?acc=GSE161360
Download Supplementary file "GSE161360_RAW.tar". The file contains multiples organisms, only GSM4905285 to GSM4905302 is the organisms of interest, thus used for this analysis.

### Step 2: Run Python to Converted wig to Gene Expression Matrix

Ensure the "convert_wig_to_expression.ipynb" file is in the same folder as the needed to processed wig.gz files.

Open the folder in VScode and run.

**Output files:**
- `pai2_expression_matrix.csv` - Raw expression values
- `pai2_expression_log2.csv` - Log2-transformed (input for bnlearn)
- `pai2_gene_coordinates.csv` - Gene coordinates used


### Step 3: Run Bayesian Network Analysis

Ensure all previous output file is in the same folder as bnlearn_analysis_annotated.R.
Open folder with VScode.

**Output files:**
- `bnlearn_all_edges.csv` - All pairwise edge bootstrap strengths (992 possible edges with their bootstrp scores)
- `bnlearn_significant_edges.csv` - Significant edges (strength > 0.5, direction > 0.5) (only 56 edges passed the filtering and use to futher analysis)


## Detailed Methods

### Data Preprocessing (Python)

The raw data from GSE161360 is provided as WIG (wiggle) format files containing per-nucleotide coverage values. The preprocessing script:

1. **Parses WIG files** for the 18 samples of interested (forward and reverse strands)
2. **Extracts gene-level expression** by averaging coverage across gene coordinates
3. **Applies strand-specific calculation** (positive strand → forward coverage; negative strand → reverse coverage)
4. **Log2-transforms** values with pseudocount of 1 to normalize distribution

Gene coordinates for PAI2 (FN0834-FN0865) were obtained from NCBI GenBank accession AE009951.2.

### Bayesian Network Learning (R)

The bnlearn analysis uses:

- **Algorithm**: Hill-Climbing (hc)
- **Score**: BIC-Gaussian (bic-g) - appropriate for continuous data with small sample sizes
- **Bootstrap**: 500 iterations for edge confidence assessment

**Significance thresholds:**
- Edge strength > 0.5 (appears in >50% of bootstrap samples)
- Direction > 0.5 (consistent direction in >50% of samples)

### Output Interpretation

| Column | Description |
|--------|-------------|
| `from` | Source gene in directed edge |
| `to` | Target gene in directed edge |
| `strength` | % of bootstrap samples containing this edge (0-1) |
| `direction` | % of samples with this specific direction (0-1) |

**Note**: Edges represent statistical dependencies. potentially suggests co-expression, but not necessarily direct physical interactions.

## Reproducibility

To ensure reproducibility:

1. Random seed is set to 42 in both Python and R scripts
2. All package versions are documented
3. Raw data is publicly available from GEO

## Limitations

1. **Sample size**: 18 samples is below the recommended 30-50+ for robust Bayesian network learning
2. **Co-expression ≠ interaction**: Edges represent statistical dependencies, not direct physical protein-protein interactions
3. **Condition-specific effects**: Results may be influenced by the specific growth conditions in GSE161360

## Troubleshooting

### Python Issues

```bash
# If missing packages:
pip install numpy pandas

# If permission error:
pip install --user numpy pandas
```

### R Issues

```r
# If bnlearn installation fails:
install.packages("bnlearn", dependencies = TRUE)

# If Bioconductor dependencies needed:
if (!requireNamespace("BiocManager", quietly = TRUE))
    install.packages("BiocManager")
BiocManager::install(c("graph", "Rgraphviz"))
```

## Acknowledgments

- Expression data from Ponath et al. (2021), GEO accession GSE161360
- bnlearn package developed by Marco Scutari
- STRING database for interaction validation
