## supplementary for "Integration of Bioinformatics and Machine Learning to characterize *Fusobacterium nucleatum*’s pathogenicity": Supplementary_Table_S2_PePPER.pdf

university of groningen

### ProPr: Prokaryote Promoter Prediction v2.0

including Transcription Start Sites (TSSs), Transcription Terminators, and Operons

Promoters as Table | Promoters as GFF | Transcription Terminators as Table | Transcription Terminators as GFF | Bookmark results here | New Sessic

Predicted Operons as GFF | Comprehensive GFF

Show Graphics | Download table

| chrom | ID | TSS | locus_tag | min10seq | min10score | strand | score | class | promoterSeq |
|---|---|---|---|---|---|---|---|---|---|
| AE009951.2 | p_0001 | 999 | p_0001 | TACTATAAT | 4 | - | 0.99 | intergenic | TTGTCCAAAAAATTAATATAGTTTTTATTAAATGC**TTTACA** |
| AE009951.2 | p_0002 | 2471 | p_0002 | AATAATATA | 3 | + | 1.0 | intergenic | ATTCTCTTGATTTAAGCTAGAAATTTAAAAATAAT**TGTTGA** |
| AE009951.2 | p_0003 | 2670 | p_0003 | ATTCTATAA | 3 | + | 0.92 | intergenic | GAAAAGAAATTTAAGTTTTTTAACTTGTTTTCTAC**ATTTTA** |
| AE009951.2 | p_0004 | 2543 | p_0004 | CAAATCAAA | 2 | - | 0.33 | intergenic | CTTAAATTTCTTTTCTTCTCTCATCCAGACTCTAC**TGTCGG** |
| AE009951.2 | p_0005 | 2383 | p_0005 | AGTTATAAT | 4 | - | 0.97 | intergenic | TTTTGTGAATTTGTCAACAATTATTTTTAAATTTC**TAGCTT** |
| AE009951.2 | p_0006 | 3760 | p_0006 | AATGTTTAA | 2 | + | 0.21 | intergenic | ATAATTAGTTATTTTATATAATATTTGTATTTGTT**GTGTGT** |
| AE009951.2 | p_0007 | 10275 | p_0007 | TTTTATATA | 3 | + | 1.0 | intergenic | TATTTAGCAAATAAAAACAATTAGATAAAAATATT**TAAAAG** |
| AE009951.2 | p_0008 | 10431 | p_0008 | TGAAATTCC | 2 | + | 0.86 | intergenic | AGGGTGAAATTCCCGACCGGTGGTACAGTCCACGA**AAGCAT** |
| AE009951.2 | p_0009 | 10462 | p_0009 | GAGTATAAT | 4 | - | 0.97 | intergenic | ATATTTAAAATTTTTGTTTTAATAAAAAAATGCTC**AAACAA** |
| AE009951.2 | p_0010 | 10197 | p_0010 | TGCTAAATA | 3 | - | 1.0 | intergenic | TATATAAAAATTAGTTTGATGTTCTTTTAAATATT**TTTATC** |
| AE009951.2 | p_0011 | 14357 | p_0011 | TAATATAAT | 5 | + | 1.0 | intergenic | TTTTTAATTTAAAAAAGTATATTACATAAATTTTT**TAGAAA** |
| AE009951.2 | p_0012 | 15947 | p_0012 | TTTTATTAA | 3 | + | 1.0 | intergenic | TCCAAAAATCAAAATGGAAGTCAAAATAGAATGAA**CTTCTA** |
| AE009951.2 | p_0013 | 26239 | p_0013 | TGTTAAATA | 4 | + | 0.89 | intergenic | TTTCCCCTTTTTATTTTTCAGTATTTAAAAATTAT**AACATA** |
| AE009951.2 | p_0014 | 26368 | p_0014 | TGATAAAAT | 4 | + | 1.0 | intergenic | ACATAAATTTATTAAGTTAAGCAAAATAAATTTTA**TTGTAT** |
| AE009951.2 | p_0015 | 26472 | p_0015 | TAGTAAAAT | 3 | + | 0.98 | intergenic | TATTAAATTTTGAGCTAGTCAATATTTTATTAATA**CTAATT** |
| AE009951.2 | p_0016 | 26339 | p_0016 | TCATATAAT | 4 | - | 0.73 | intergenic | AATTTAATATATTAAATATATTAATTCTATTTAGT**AAACAT** |
| AE009951.2 | p_0017 | 26191 | p_0017 | TGTTATAAT | 5 | - | 1.0 | intergenic | TGAAAAATAAAATATAATTCTATTTCATAATATTT**AACACA** |
| AE009951.2 | p_0018 | 43684 | p_0018 | ATTTATAAT | 5 | + | 0.78 | intergenic | TATTAGTATATTTTTAGATTAAAATAATCTATTTA**TAGATA** |
| AE009951.2 | p_0019 | 43646 | p_0019 | AAATATAAT | 5 | - | 0.14 | intergenic | TTGTAAGAAACAAGAGTTAGTTCTTTTTAATTTAT**TAAAAA** |
| AE009951.2 | p_0020 | 43507 | p_0020 | TGATATAAT | 5 | - | 1.0 | intergenic | AGCATATAAAAATGAATATCAGATGAATATTTTTG**ATGACA** |
| AE009951.2 | p_0021 | 49912 | p_0021 | TGTTAAATT | 3 | - | 1.0 | intergenic | TAATAGAAAAATATATAAATAAATATAAAAAAAGT**TTGAAA** |
| AE009951.2 | p_0022 | 52496 | p_0022 | AATTATAAT | 5 | + | 0.98 | intergenic | TCATATTTTAACATAATAAGCAATATTTAAGATTT**TTTAAA** |
| AE009951.2 | p_0023 | 52422 | p_0023 | TGTTAAAAT | 4 | - | 1.0 | intergenic | ATATATTATAATTATTTGTCAAAAATATTTAAAAA**ATCTTA** |
| AE009951.2 | p_0024 | 61734 | p_0024 | TATAAAAAA | 3 | - | 1.0 | intergenic | TTATTCGTGATAAAGGTACAAAAAATAAAAATAAA**CATTGA** |
| AE009951.2 | p_0025 | 63475 | p_0025 | TTTAAAATT | 3 | + | 1.0 | intergenic | TCATATTTTCTTTAAAATCATCAATCTTTTTATAC**TTTTTT** |
| AE009951.2 | p_0026 | 63402 | p_0026 | TGATTTTAA | 3 | - | 1.0 | intergenic | ATTGTAAATAAAATTTTAAAAATGTTTAAAAAAAG**TATAAA** |
| AE009951.2 | p_0027 | 68645 | p_0027 | ACTTATAAT | 4 | - | 1.0 | intergenic | TCTTTACTTTTTATAATAATTAAAATAAAAATAAA**TTGAAA** |
| AE009951.2 | p_0028 | 76106 | p_0028 | TAGTATAAT | 4 | - | 1.0 | intergenic | ATTTGGAATTATTTTAAAAATAATATATTTAGATA**TTGTAT** |
| AE009951.2 | p_0029 | 76002 | p_0029 | AATAAAAAA | 3 | - | 0.98 | intergenic | ATAAAAAAGGAATTTTTATATTTTTCTTTTTTTAT**TCTTAT** |
| AE009951.2 | p_0030 | 75909 | p_0030 | TATTATAAA | 3 | - | 0.96 | intergenic | ATTATTTTATAAGTAAATTTCAAATAGCTAGATTA**TTTTAT** |
| AE009951.2 | p_0031 | 80682 | p_0031 | TTTTTATAT | 3 | + | 0.93 | intergenic | ATCCTCCTTTCTTATTTTTTATTATTTTTATAATT**ATAACT** |
| AE009951.2 | p_0032 | 80803 | p_0032 | TGCTATAAT | 5 | + | 0.95 | intergenic | TAGTTTATTACTAAGTAGATTTTTTTATTAATTAC**TATAAA** |
| AE009951.2 | p_0033 | 80792 | p_0033 | ATAATTATT | 3 | - | 0.98 | intergenic | CTAAGTCAAAATTTTTAATTTCTTTTAAATTCATT**TTAAAC** |
| AE009951.2 | p_0034 | 80650 | p_0034 | GTTTATAAT | 4 | - | 1.0 | intergenic | GAAATAAAAATTTTTACTAAGTAAAGAAATTATGG**TTGACA** |
| AE009951.2 | p_0035 | 93547 | p_0035 | TGTAAAATA | 4 | - | 1.0 | intergenic | AACTTTTTTATTTTTAAAAAATAAAAATAAATCAT**AGAAAT** |
| AE009951.2 | p_0036 | 93474 | p_0036 | TTTTAAAAT | 3 | - | 0.65 | intergenic | AAATGTGAGATTAATATAAAAATAAAGTTTTTCTT**GAAATA** |
| AE009951.2 | p_0037 | 98696 | p_0037 | TGATCTATT | 2 | + | 1.0 | intergenic | TCCACACAAAACCACAGACTCTCACCGTGATTTTG**CTGGGA** |
| AE009951.2 | p_0038 | 98867 | p_0038 | TACTAAAAT | 3 | + | 1.0 | intergenic | TCTTTATTTTTATAACTAAAAATAAACATTTTATT**ATATAT** |
| AE009951.2 | p_0039 | 98775 | p_0039 | TAAATAATA | 3 | - | 0.98 | intergenic | AAAAATAATATATATAATAAAATGTTTATTTTTAG**TTATAA** |

Background image is our builing Linnaeusborg on the Zernike campus. Image by Anne de Jong

2025-04 259
2025-05 227
2025-06 245
2025-07 205
2025-08 82
Successful Sessions 2025: 3450

University of Groningen
Molecular Genetics
the Netherlands

Anne de Jong and Auke van Heel
Co-authors: Hung Chu, Jan Kok, Oscar Kuipers
Master students: Max Luppus, Dan Kaptijn, Inge Hendriks
Bachelor students: Tomas Vogels, Michael van Dijk, Marlon van Es, Ezvin Herdic

MolGen

| | | | | | | | | | |
|---|---|---|---|---|---|---|---|---|---|
| AE009951.2 | p_0045 | 109061 | p_0045 | AATGAAAAT | 3 | - | 1.0 | intergenic | GGAAAATATACTAATTTTAATCTTTTTTTATTGAC**AAATTA** |
| AE009951.2 | p_0046 | 111024 | p_0046 | TGCTATAAT | 5 | - | 1.0 | intergenic | TTTCTTTGTTTATAACAGCTTCTCTTGTTTTTCTA**TGGTAT** |
| AE009951.2 | p_0047 | 111626 | p_0047 | TTTTAAAAT | 3 | - | 0.99 | intergenic | GGTAAAAAAGTTGAAGTAACTGAGAAAGATGAGCA**TAGACA** |
| AE009951.2 | p_0048 | 119732 | p_0048 | TGGTATAAT | 5 | - | 1.0 | intergenic | GATTTTTTTACTTAAAGATTATAACAAAATAATAA**AGACTA** |
| AE009951.2 | p_0049 | 125239 | p_0049 | TGCTACAAT | 2 | + | 0.99 | intergenic | GGAATGTAATGAGCTATTTTCTATATTATTTATTC**TTTACT** |
| AE009951.2 | p_0050 | 125143 | p_0050 | TCTTAAAAT | 3 | - | 0.52 | intergenic | AAAAAAGTAAAGAATAAATAATATAGAAAATAGCT**CATTAC** |
| AE009951.2 | p_0051 | 127921 | p_0051 | TGGTAGAAT | 2 | + | 1.0 | intergenic | TAACATAAAAAGAATACTATTACAAGAGAGAATAA**TTGACA** |
| AE009951.2 | p_0052 | 127839 | p_0052 | TGTTAAAAT | 4 | - | 1.0 | intergenic | TACCATATTTTTCAGACTTTGTCAATTATTCTCTC**TTGTAA** |
| AE009951.2 | p_0053 | 141978 | p_0053 | TAAAAATTT | 3 | + | 1.0 | intergenic | GAGTATAACATACTTTTTAAAAAAATAAAAGAAAA**ATTTTT** |
| AE009951.2 | p_0054 | 142015 | p_0054 | AGAATTTTA | 3 | - | 1.0 | intergenic | TCCAAAAATAAAATAATTTGTAAAATTTTAAATAT**TTAAAT** |
| AE009951.2 | p_0055 | 141906 | p_0055 | TTTAAAAAG | 3 | - | 1.0 | intergenic | AGTAGTATAAAAAAATTTTTAATGATTTAAAAAAT**TTTTCT** |
| AE009951.2 | p_0056 | 147666 | p_0056 | TTAATAAAA | 3 | - | 1.0 | intergenic | AAAAATTGCTTTATAAAAAAAGTTTAGCATATTAA**TTATTT** |
| AE009951.2 | p_0057 | 149154 | p_0057 | ATATATAAT | 5 | - | 1.0 | intergenic | ACTAATTTTTTAACAGTTTCTTTTTATTTTTGCTA**ATATTA** |
| AE009951.2 | p_0058 | 155153 | p_0058 | TGATAAAAT | 4 | + | 1.0 | intergenic | CCAATCTTCTTGTAATTATTTTTTAACAGAAAACC**TTTATA** |
| AE009951.2 | p_0059 | 155094 | p_0059 | TGTTAAAAA | 4 | - | 0.19 | intergenic | GTATTTTATTTATACTTCAATTTTATCATCTTTTT**TAAATA** |
| AE009951.2 | p_0060 | 155011 | p_0060 | TAGAATTAT | 3 | - | 0.99 | intergenic | GATATAGAATATGATATAATTAAATATTTTAGTTG**GAAAAG** |
| AE009951.2 | p_0061 | 158828 | p_0061 | TGTTAAAAT | 4 | - | 1.0 | intergenic | TTGAATGGTATGTGAAAATTTAATAAAATTAAAAC**TAGAAT** |
| AE009951.2 | p_0062 | 162093 | p_0062 | TTAAAAAAT | 3 | + | 1.0 | intergenic | CTATTTAGAACTTTTATAGTTTAATTACCAAATAA**AAATTT** |
| AE009951.2 | p_0063 | 162245 | p_0063 | TGATAAAAT | 4 | + | 1.0 | intergenic | TGAATATATATTGTTGTTAATAAAGGATATTTTTA**TGGAAA** |
| AE009951.2 | p_0064 | 161894 | p_0064 | AACATATAT | 3 | - | 1.0 | intergenic | GCTAAGGATTACTTTTATTTTTTATTTATATAAAA**ATCTTG** |
| AE009951.2 | p_0065 | 164621 | p_0065 | ATAAATAAG | 3 | - | 0.99 | intergenic | AAACTAAAATTTATTAGTTATTTAAAAATTTATAA**TTAATT** |
| AE009951.2 | p_0066 | 168380 | p_0066 | TGTTATAAT | 5 | - | 0.94 | intergenic | AAACCCACTATGTAGATAGTTTTCTCTTTAAACTA**TTGACT** |
| AE009951.2 | p_0067 | 172695 | p_0067 | TTATATAAT | 5 | + | 1.0 | intergenic | TATAAAATATTTATCTATACAACCTCCTTTCATAG**TTTATT** |
| AE009951.2 | p_0068 | 172607 | p_0068 | TTATATATT | 3 | - | 0.2 | intergenic | AAATATTAAATAAAATAAACTATGAAAGGAGGTTG**TATAGA** |
| AE009951.2 | p_0069 | 174590 | p_0069 | ATATTAATA | 3 | - | 0.36 | intergenic | TCTTTTCTGTAAATTTTCTTCTTCTATGATTAGAA**TTATTT** |
| AE009951.2 | p_0070 | 174425 | p_0070 | TGTAAAATA | 4 | - | 1.0 | intergenic | TAACATAATTAAGAGAATTTTTGAGAATTAAAAAC**TTAAAA** |
| AE009951.2 | p_0071 | 175850 | p_0071 | ACTTAAAAT | 3 | - | 1.0 | intergenic | AATATTTTTAATACCTTCAATAATTATAATTGATA**TTTTTA** |
| AE009951.2 | p_0072 | 195217 | p_0072 | ATTTATAAT | 5 | - | 1.0 | intergenic | TGAAAAAGGGAAGGTAAAAAATTATTAAGAAAATA**GAAACT** |
| AE009951.2 | p_0073 | 194952 | p_0073 | GTAGAATTA | 2 | - | 1.0 | intergenic | TGATATAGCTATTGATTTAAGGGAAAATAGTGAGA**CTTTTG** |
| AE009951.2 | p_0074 | 199207 | p_0074 | AATTAAAAA | 3 | + | 1.0 | intergenic | TAAAGTAAAAATTTATTTTATTTTATATTATATCA**TACAAT** |
| AE009951.2 | p_0075 | 199155 | p_0075 | TGATATAAT | 5 | - | 0.97 | intergenic | ACCACAAATTTTCTAAATATTAATAATTTTTAATT**ATGATT** |
| AE009951.2 | p_0076 | 212589 | p_0076 | TTGTATAAT | 4 | + | 0.97 | intergenic | GCTCACAATTTCAATTATAGTCGAAATGATAAAAT**TTAAAT** |
| AE009951.2 | p_0077 | 231141 | p_0077 | TATATAAAA | 3 | - | 0.31 | intergenic | TGAATGGAATTTTCAAGAAAAATTAATTTTTATTT**AAAAGT** |
| AE009951.2 | p_0078 | 232195 | p_0078 | TGTTATAAT | 5 | - | 1.0 | intergenic | GAAAGGATTTTTTTAAATCTTCAATCAGCTTTTTT**TGTTAA** |
| AE009951.2 | p_0079 | 240938 | p_0079 | GAGTATAAT | 4 | - | 1.0 | intergenic | TTTTTAGATATATAAGATAAAAAACAAAATTTTTG**GTTTTA** |
| AE009951.2 | p_0080 | 244324 | p_0080 | TAGTATATT | 3 | + | 0.29 | intergenic | ATAAATAAAATTTTTTTTGTTTTTTGCTTATGTAT**TTCAGC** |
| AE009951.2 | p_0081 | 244460 | p_0081 | AGTTAAAAT | 3 | + | 0.99 | intergenic | CTTTAAAAAAATAATGTTAATTTTATTTTTTATTT**GACTCT** |
| AE009951.2 | p_0082 | 244335 | p_0082 | TATTAAAAA | 3 | - | 1.0 | intergenic | ACATTATTTTTTTAAAGAATACAGAAAATATAAAA**AATAGT** |
| AE009951.2 | p_0083 | 255205 | p_0083 | TGCTATTCC | 2 | - | 1.0 | intergenic | AGCGCAGCCCGGTAGCGCACCTGCCTTGGGAGCAG**GGGGTC** |
| AE009951.2 | p_0084 | 256695 | p_0084 | TGTAAAATT | 3 | - | 1.0 | intergenic | GAGACTACTTTTAGAGCAGTCTCTTTTTTTTATTGT**TGTAAA** |
| AE009951.2 | p_0085 | 264700 | p_0085 | TTCTAAAAT | 3 | + | 1.0 | intergenic | AATCTATCTAATTTATAGTATTTTTTAGTAAATTC**ATAAAA** |
| AE009951.2 | p_0086 | 264640 | p_0086 | TAAAAAATA | 3 | - | 1.0 | intergenic | TTGTCAATAATATTTTAAATTTTAGAAAAAAAATA**AAAAAA** |
| AE009951.2 | p_0087 | 266879 | p_0087 | TGCTATAAT | 5 | + | 1.0 | intergenic | TTACCATAAAAAATTATATTTTACCATTTATAAAT**ATTTTA** |
| AE009951.2 | p_0088 | 266798 | p_0088 | TGGTAAAAT | 3 | - | 1.0 | intergenic | ATAGCATAAAATATAAAACTTAAAATATTTATAAA**TGGTAA** |
| AE009951.2 | p_0089 | 273405 | p_0089 | TGTATTATA | 3 | - | 1.0 | intergenic | TGTATACCAATATAATTGATTTTGAAAAAAAGTCT**TGATTT** |
| AE009951.2 | p_0090 | 275728 | p_0090 | AAAAAAACT | 2 | - | 1.0 | intergenic | AATAGAAAAAAGATAATATTTTAATCATAAATTTA**ATTGAC** |
| AE009951.2 | p_0091 | 278567 | p_0091 | ATAGTATAT | 3 | + | 0.99 | intergenic | TCTAAATTACTAATACATTTAAAAATATTTATTTT**TTATTG** |
| AE009951.2 | p_0092 | 278530 | p_0092 | AATTAAATA | 3 | - | 0.93 | intergenic | AAATTGTGATAGTAAAATAATAATTTTAAAGTATC**TATATG** |
| AE009951.2 | p_0093 | 278943 | p_0093 | TGGTAATTT | 3 | + | 1.0 | intergenic | ATAAGGTAAAGTCTTATTGAAAATGTTGAGATTTA**ATGATA** |
| AE009951.2 | p_0094 | 284076 | p_0094 | TGTATAGTT | 2 | + | 0.76 | intergenic | ATAGCATTTTTATATAAAATTAGCAATAAATATAA**ATAGTA** |
| AE009951.2 | p_0095 | 284181 | p_0095 | TATCTTTTT | 2 | + | 0.08 | intergenic | AAATAATATAAATAATTTTTATATAGAAAAGTTTC**ATAAAT** |
| AE009951.2 | p_0096 | 284290 | p_0096 | TTTTATAAT | 5 | + | 1.0 | intergenic | AATATATATTATATATATAATTTTTAAAAAAATAA**TTGACA** |
| AE009951.2 | p_0097 | 284279 | p_0097 | TTTTATAAT | 5 | - | 0.97 | intergenic | TAACCCCTCTTATATTTTAATGATAATTTATGTAT**AAAAAA** |
| AE009951.2 | p_0098 | 284110 | p_0098 | ATAAAAATT | 3 | - | 0.98 | intergenic | TATAAAGAAAAAGATAAGAACGAAAGAAAAATTTA**TGAAAC** |
| AE009951.2 | p_0099 | 283990 | p_0099 | TGATTTATT | 3 | - | 0.98 | intergenic | AAAATTATCTATACATACTATTTATATTTATTGCT**AATTTT** |
| AE009951.2 | p_0100 | 290089 | p_0100 | TCTTAAAAT | 3 | - | 0.88 | intergenic | GATCATTTCACTCAAACACAGCAAGATTTGCTCGG**TTCATT** |
| AE009951.2 | p_0101 | 292057 | p_0101 | TAGAATAAA | 3 | - | 0.98 | intergenic | TTAATTTTTTATCTAAAATTTAGAATGCAATTCAC**TTATTT** |
| AE009951.2 | p_0102 | 295575 | p_0102 | TGATAAAAT | 4 | - | 0.99 | intergenic | TTCTCTTTTTTTATTGTTAAATCGCATTATTACAT**TAAAAA** |
| AE009951.2 | p_0103 | 295449 | p_0103 | AAATATAAT | 5 | - | 1.0 | intergenic | AAAAGTTTTTTTTATTTTCAATTTTATTCTATTAA**GCAAAG** |
| AE009951.2 | p_0104 | 296145 | p_0104 | TAATATAAT | 5 | - | 1.0 | intergenic | TGTTGCAAATTAAATAGTATACTACATAAATTTTT**TAGAAA** |
| AE009951.2 | p_0105 | 314102 | p_0105 | AAGAAAAAA | 3 | + | 0.46 | intergenic | TACATTTTTAAAATTTTTTAAACCTTTACTAAAAA**GACAAA** |
| AE009951.2 | p_0106 | 314022 | p_0106 | TTTTAAAAA | 3 | - | 1.0 | intergenic | TTTTCTTATATTATATTAGAATTTGTCTTTTTAGT**AAAGGT** |

| | | | | | | | | | |
|---|---|---|---|---|---|---|---|---|---|
| AE009951.2 | p_0107 | 327520 | p_0107 | AACTTTTTA | 2 | + | 0.92 | intergenic | AGTTTGTTTAATTTTTTTACAAAAAATAAGAAATA**GAAGAT** |
| AE009951.2 | p_0108 | 327613 | p_0108 | TGTTATAAT | 5 | + | 1.0 | intergenic | GTTTTATATAATTACAATTTTTTCTAATTAGCTCT**CTTTAT** |
| AE009951.2 | p_0109 | 327369 | p_0109 | TGTTATAAT | 5 | - | 1.0 | intergenic | TTTTTGAAGTCTAAGGTAAATAATTAATCTAAAAA**TTGTAA** |
| AE009951.2 | p_0110 | 333696 | p_0110 | TGTTATAAT | 5 | + | 1.0 | intergenic | TTTCAATAAAGAATAAATTTAATAAAATATTAATT**TTTTAT** |
| AE009951.2 | p_0111 | 341705 | p_0111 | TATTATAAA | 3 | + | 1.0 | intergenic | AATATTTTTTTTAAGGATAATTTGATTTAAAAAAT**AATTGA** |
| AE009951.2 | p_0112 | 342231 | p_0112 | TGTTATAAT | 5 | + | 1.0 | intergenic | ATAAGAATCCCTTTAAAAAAGAGGGATTTTTATTT**TACTTT** |
| AE009951.2 | p_0113 | 348395 | p_0113 | TTAAAATAA | 3 | - | 1.0 | intergenic | CTAAAATATTGTACTAAAAAAGTAAGGATTTAGTT**GACTTT** |
| AE009951.2 | p_0114 | 349906 | p_0114 | TATTAAAAT | 3 | - | 1.0 | intergenic | CTTAAAAAGTTTCTCCTTAGTTTAAAAATAAAATA**AAAATT** |
| AE009951.2 | p_0115 | 362734 | p_0115 | ATTTTATAT | 3 | - | 1.0 | intergenic | TAAGAGAATTTTTTGAGAGCAAAAACTTAAAAATT**CTCTTT** |
| AE009951.2 | p_0116 | 365225 | p_0116 | TACTATAAA | 3 | - | 0.88 | intergenic | ATAAGGAAAAAGTCTTATTTAATGGTTTATAACTA**TTTTTT** |
| AE009951.2 | p_0117 | 366726 | p_0117 | TATTAATAT | 3 | + | 0.96 | intergenic | AACTAACAAAAAATCTTTTATAAAATCTTGTTTTA**TCTATC** |
| AE009951.2 | p_0118 | 366817 | p_0118 | TAGTATAAA | 3 | + | 0.89 | intergenic | TGATACATTTTTATTATATCACAATAATATTATCA**TTGACA** |
| AE009951.2 | p_0119 | 366751 | p_0119 | TGATATAAT | 5 | - | 1.0 | intergenic | ATCTTGTAGGTATTTATACTATTTTTATTTGGTTT**TGTCAA** |
| AE009951.2 | p_0120 | 367649 | p_0120 | TAATATAAT | 5 | + | 1.0 | intergenic | TCAAAAGTAAAGGAGTTTTTAAACTCCTTTTTCTC**TTGTAA** |
| AE009951.2 | p_0121 | 368820 | p_0121 | TTATATAAT | 5 | + | 1.0 | intergenic | TATAAAATATTTATCTATACAACCTCCTTTCATAG**TTTATT** |
| AE009951.2 | p_0122 | 370761 | p_0122 | AACTTAATT | 3 | + | 1.0 | intergenic | TTATTAATTTTATAAACTTTTATTTTTTTCTTGAT**TTTTTC** |
| AE009951.2 | p_0123 | 370647 | p_0123 | TGATACTAT | 2 | - | 1.0 | intergenic | AAAAAATAAAAGTTTATAAAATTAATAAAAAATAG**TTGACA** |
| AE009951.2 | p_0124 | 373519 | p_0124 | TGATATAAA | 4 | - | 1.0 | intergenic | TGTTGCTATTGCTTTTTTAGATAAAATAATATATA**ATTGAA** |
| AE009951.2 | p_0125 | 374341 | p_0125 | ATAAATAAT | 3 | - | 0.81 | intergenic | TACATTCTTTTATTATACAATCAAAAAATTGACAA**TATACT** |
| AE009951.2 | p_0126 | 381725 | p_0126 | TGTTATTAT | 3 | - | 1.0 | intergenic | CTTTTTTATTTCTTTATAAAAAAATAAAAAATTAT**TTGACA** |
| AE009951.2 | p_0127 | 381624 | p_0127 | ATTAAAAAA | 3 | - | 0.98 | intergenic | AAAAAGTCTGTTTTTTTCCTAAATGTTTGGTATAA**AGGTAA** |
| AE009951.2 | p_0128 | 382539 | p_0128 | TAAAATATA | 3 | - | 0.1 | intergenic | AGATTTGTAAATTTTAATATTCTATGGTATAATAC**ATCAGT** |
| AE009951.2 | p_0129 | 388133 | p_0129 | TGTTATTAT | 3 | + | 1.0 | intergenic | ATACCTCCTCCTAATAATAAAAACTGTTTATTTGT**TTATCA** |
| AE009951.2 | p_0130 | 388338 | p_0130 | GATTATAAT | 4 | + | 0.98 | intergenic | ATTAGATACATATTATTTATATTTTTTTAATAATA**CTTGAA** |
| AE009951.2 | p_0131 | 388258 | p_0131 | ATCTAATAT | 3 | - | 1.0 | intergenic | TATAATCATAAAAAAAATTTTTTTCAAGTATTATTA**AAAAAA** |
| AE009951.2 | p_0132 | 388114 | p_0132 | ATATATAAT | 5 | - | 0.76 | intergenic | CGACACTAAAAAAATATAAATATTATAAAAAGATA**TTGACT** |
| AE009951.2 | p_0133 | 390681 | p_0133 | TTCTAATTT | 3 | - | 1.0 | intergenic | AAAGAAGCAAAAATATTTTAGAATGAAAATTCATT**TATTTT** |
| AE009951.2 | p_0134 | 393230 | p_0134 | ATAGTTATA | 2 | - | 0.56 | intergenic | TTGATAAAAAAGTAAATACGGACAAAAAAGACTGT**ACTTTA** |
| AE009951.2 | p_0135 | 398323 | p_0135 | CTTTATAAT | 4 | + | 0.97 | intergenic | ATTTAAAGAACAATATACGCTCTTTATCCTCCCTT**TTCTAA** |
| AE009951.2 | p_0136 | 398472 | p_0136 | TGATATATT | 3 | + | 1.0 | intergenic | TATTTATTAATAAAAGCTTAAAATTTTAAAATAAG**TTGAAA** |
| AE009951.2 | p_0137 | 398383 | p_0137 | AATAAAACA | 2 | - | 0.48 | intergenic | TTATGTTTAATATTTCAACTTATTTTAAAATTTTA**AGCTTT** |
| AE009951.2 | p_0138 | 398306 | p_0138 | TGCTATATT | 3 | - | 1.0 | intergenic | TAAAGTATGACAAAATATATTTTAAAAATATAATT**TGACGA** |
| AE009951.2 | p_0139 | 410022 | p_0139 | CGTTATAAT | 4 | - | 1.0 | intergenic | TCAATATCATTTTTTATTAAAAATTAATATATTTC**TTGTAT** |
| AE009951.2 | p_0140 | 425401 | p_0140 | AGGTATAAT | 4 | - | 1.0 | intergenic | GTGATAGCTATATTTTTTAAATAGTTTTCTTAAAT**TTGCAA** |
| AE009951.2 | p_0141 | 434215 | p_0141 | ATCATAATA | 3 | - | 0.96 | intergenic | TTTTCTTTTACTAAAAATTTTTAATCAAACCTTAA**TTTTAA** |
| AE009951.2 | p_0142 | 441258 | p_0142 | TGATATATA | 4 | + | 1.0 | intergenic | AATCTAAAAAAAATTTTTAATTTTTTAAAAAAACTC**TTGACA** |
| AE009951.2 | p_0143 | 441160 | p_0143 | TTAATAAAA | 3 | - | 1.0 | intergenic | AAATGTCAAGAGTTTTTTTAAAAAATTAAAAATTT**TTTTAG** |
| AE009951.2 | p_0144 | 444926 | p_0144 | ATTAAAATA | 3 | + | 0.99 | intergenic | ATTTTTATTATGGAATATATTTTTTTAGAATAGTT**TTTTCG** |
| AE009951.2 | p_0145 | 444850 | p_0145 | ATAATAAAA | 3 | - | 1.0 | intergenic | TTAAATAATATTTTAATCTTTAAAACGAAAAAACT**ATTCTA** |
| AE009951.2 | p_0146 | 446731 | p_0146 | AGCTATTTT | 2 | + | 1.0 | intergenic | GGCTGTTGCATTTTTATTATTTACCCCTCTTTAAT**ATATAA** |
| AE009951.2 | p_0147 | 455032 | p_0147 | TATAAAATA | 3 | - | 1.0 | intergenic | AATAGTTTCTTTTTTATTATTCCTAAAAATAGATA**TGGACT** |
| AE009951.2 | p_0148 | 458306 | p_0148 | TGTTATATT | 3 | - | 1.0 | intergenic | TCTAAAAATGAAGGTGTTATTTAAAAAAATTATAC**TTGAAA** |
| AE009951.2 | p_0149 | 459182 | p_0149 | ATGTATAAT | 4 | - | 0.8 | intergenic | CTTATGCCTTTGGACCAAAAGTTACAGATAAAGTT**AGATTT** |
| AE009951.2 | p_0150 | 460130 | p_0150 | GAGTATAAT | 4 | - | 1.0 | intergenic | TTTTATTTTTTGAACAAATAGTATAAAAATATAAT**ATTGAA** |
| AE009951.2 | p_0152 | 472184 | p_0152 | GTTTATAAT | 4 | - | 0.98 | intergenic | AGTTTTTTGTGTCTAAATTAACAAGAGAAGATAAA**TTGACA** |
| AE009951.2 | p_0153 | 475716 | p_0153 | TGGTATAAT | 5 | + | 1.0 | intergenic | TTTACCACATATTTTCTAATTAAATTAATATTTTA**TATTGT** |
| AE009951.2 | p_0154 | 475636 | p_0154 | TGGTAAAAT | 3 | - | 1.0 | intergenic | TATACCACAAAATATTGAATTACAATATAAAATAT**TAATTT** |
| AE009951.2 | p_0155 | 483540 | p_0155 | TGATATTAT | 3 | + | 0.9 | intergenic | AAATCGCACGGAATTTTAAAGTTCCTAAATCTTCT**CCTAAT** |
| AE009951.2 | p_0156 | 485618 | p_0156 | TAGTAAATT | 3 | + | 0.12 | intergenic | TTGATATTTTTAGCAATATAGAATAAAATTGAATT**AAATAT** |
| AE009951.2 | p_0157 | 488866 | p_0157 | TGAAAAAAA | 4 | + | 1.0 | intergenic | ATTTATTTTATTTACAATAATTTTTATTTTTTTAAG**GTATAG** |
| AE009951.2 | p_0158 | 488817 | p_0158 | TACTTAAAA | 3 | - | 1.0 | intergenic | CTCCTCCATAAAATTTTTGTTAACTTATCATATCA**TTTTTT** |
| AE009951.2 | p_0159 | 491010 | p_0159 | TACTTAATT | 3 | + | 1.0 | intergenic | TTTACTAATTTAAAACACTATCAAAATAATTTGAC**AAATAA** |
| AE009951.2 | p_0160 | 490799 | p_0160 | TGTTAAATA | 4 | - | 1.0 | intergenic | TTTTTAGTTTTATAAGTGGAAGAAATTTCATTTTT**TTATTA** |
| AE009951.2 | p_0161 | 492563 | p_0161 | TATTAAAAT | 3 | + | 0.21 | intergenic | GAGAGGGCTAACCTCTCCTTTTTTTAAAAATTATA**TTATGA** |
| AE009951.2 | p_0162 | 494009 | p_0162 | TGGTATAAT | 5 | + | 1.0 | intergenic | TTATACATTTTTATTTGTGATAGTCTTTTTTTATT**TGTAAA** |
| AE009951.2 | p_0163 | 498249 | p_0163 | AATCCAATA | 2 | + | 0.92 | intergenic | TTAAAAAACTAGACTTGGGGTGCAAGTCCCCTTTT**TTATTG** |
| AE009951.2 | p_0164 | 499935 | p_0164 | ATAATATAT | 3 | + | 1.0 | intergenic | TCTTTTTATTTTGGATAATATATTATCTAAATATT**GACTTT** |
| AE009951.2 | p_0165 | 503859 | p_0165 | TGATATAAT | 5 | + | 1.0 | intergenic | AACATATAATTATTAAGATTTGGTTAATTTTTAAT**AGACAC** |
| AE009951.2 | p_0166 | 503776 | p_0166 | TGTTATAAT | 5 | - | 0.94 | intergenic | ATCAGAAATTAGGAATAAGTGTCTATTAAAAATTA**ACCAAA** |
| AE009951.2 | p_0167 | 506169 | p_0167 | ATCTAAAAT | 3 | + | 1.0 | intergenic | ACTCACTTCATTCAAACACAGCGAGATTTGCTCGG**CTCATT** |
| AE009951.2 | p_0168 | 507582 | p_0168 | TGTGAAAAA | 3 | - | 0.98 | intergenic | ATTTTGAGATTTTAGATGAAGAGAAAGTTTAGAAA**AACTTA** |

| | | | | | | | | | |
|---|---|---|---|---|---|---|---|---|---|
| AE009951.2 | p_0169 | 510434 | p_0169 | TACTTAAAT | 3 | - | 1.0 | intergenic | TGTAAATTTGCAATAGCCCTTATTTTTTGTTTTAC**AAATTA** |
| AE009951.2 | p_0170 | 526835 | p_0170 | TATTAAATA | 3 | - | 1.0 | intergenic | TAATTTTTAAAATTAGAATTTAATTGTATTTTAAT**TTTTAA** |
| AE009951.2 | p_0171 | 534003 | p_0171 | TTATATAAT | 5 | - | 1.0 | intergenic | AGTTTTTAAATTATAAAAAAATTTTCAAAAAAAAT**TTCTCA** |
| AE009951.2 | p_0172 | 546640 | p_0172 | TGATAAAAA | 4 | - | 0.01 | intergenic | TTATAAATAATTTTTAATATCTTAAAAATAGTTTA**TTGCTA** |
| AE009951.2 | p_0173 | 549957 | p_0173 | ACACATGCA | 1 | - | 1.0 | intergenic | ACACATGCAGGTCAATGGCTCAATTGGTAGAGCAT**CGGTCT** |
| AE009951.2 | p_0174 | 551479 | p_0174 | TGTTATAAT | 5 | - | 1.0 | intergenic | TTTATAGAGTATTATTTTCTAACAAAAAAGAAGGT**TGTTTT** |
| AE009951.2 | p_0175 | 561103 | p_0175 | TAGAATATT | 3 | - | 0.98 | intergenic | ATAAAAAATTATTGGTACTTAAATAAAAAATGGTTG**ATTTTA** |
| AE009951.2 | p_0176 | 562654 | p_0176 | TGATATAAT | 5 | + | 0.96 | intergenic | TTTAATTAAATCTATCATAATATTCTTATTTAGTC**AAGCAA** |
| AE009951.2 | p_0177 | 562587 | p_0177 | AGAATATTA | 3 | - | 1.0 | intergenic | AATTTTTCTAAATTATATCATTTTTTTACAAATAT**TGCTTG** |
| AE009951.2 | p_0178 | 565723 | p_0178 | TGATATAAT | 5 | - | 1.0 | intergenic | ACAAAAATAAACTCTCTTTTGATAGAAATATCAAA**GGAGAG** |
| AE009951.2 | p_0179 | 575203 | p_0179 | AAGAAAAAA | 3 | - | 1.0 | intergenic | AAGTTTTTTTAAACTTTATTTATCTAAAATTTCGT**TTATTT** |
| AE009951.2 | p_0180 | 575939 | p_0180 | TTTTATAAA | 3 | + | 1.0 | intergenic | ATAATATAACTTTATATTTTTATTTTGTCAAATAT**TTTTTT** |
| AE009951.2 | p_0181 | 575866 | p_0181 | AAAATATAA | 3 | - | 1.0 | intergenic | TATATTTTATAAAAAGTTTGTAAAAACTAAAAAAA**TATTTG** |
| AE009951.2 | p_0182 | 580499 | p_0182 | TGATATAAT | 5 | - | 1.0 | intergenic | TTATACATATTGATATAAAATTTTAGACTAAAATC**TTTGAG** |
| AE009951.2 | p_0183 | 582091 | p_0183 | TGATATAAT | 5 | - | 1.0 | intergenic | GATTTGAAAGTAATTCTTTTTTATTTTTTAAAAAT**AAAATA** |
| AE009951.2 | p_0184 | 582573 | p_0184 | TTCTAAAAT | 3 | + | 1.0 | intergenic | GATATTTTATCACTTTTTAACGAAAAAATATACTG**TTTTTT** |
| AE009951.2 | p_0185 | 582498 | p_0185 | TGATAAAAT | 4 | - | 1.0 | intergenic | TAAATTTTAGAAAATAAAAAAAGTTAAAAAAACAG**TATATT** |
| AE009951.2 | p_0186 | 586438 | p_0186 | ATATATAAT | 5 | - | 0.72 | intergenic | TAATGTAATTATTGTGGAAGGTTATTTTTACAGAT**AGTATC** |
| AE009951.2 | p_0187 | 587997 | p_0187 | AATTAATAT | 3 | + | 0.72 | intergenic | TTAAAATATATTAATTTTTATTATGAAAGATATAA**AATATA** |
| AE009951.2 | p_0188 | 593180 | p_0188 | TTTTTTTAA | 3 | - | 1.0 | intergenic | TTCCTACAAAGTTTTTTAAATAATTTTATATAAAG**TATTGA** |
| AE009951.2 | p_0189 | 595821 | p_0189 | TGATATAAT | 5 | + | 1.0 | intergenic | GCCTATCAAAATTTATCTATATATTATCACAAATT**TTAAAT** |
| AE009951.2 | p_0190 | 595777 | p_0190 | TAAAAAATA | 3 | - | 1.0 | intergenic | TGTCTCTCCCTTCTTTATAAAATATGCTTTTAAAA**TTATAT** |
| AE009951.2 | p_0191 | 597408 | p_0191 | TGATATAAT | 5 | - | 1.0 | intergenic | AAAAACCACAAGTCTCTCCCCTCATTTCTTGTGGT**TTTTTG** |
| AE009951.2 | p_0192 | 597262 | p_0192 | TGTTATAAA | 4 | - | 0.88 | intergenic | TTTTTTTCCCATAAAATATATAACAAATGTTTATC**TTTAAT** |
| AE009951.2 | p_0193 | 598872 | p_0193 | AAAATAATA | 3 | - | 0.78 | intergenic | GTAAATTCAATTTTACAATTTCTTGCTGAAAGCTA**TGATTT** |
| AE009951.2 | p_0194 | 607357 | p_0194 | GGCTATAAT | 4 | - | 1.0 | intergenic | TATTTAATAAATAGGAGGAAAAAATGAAAATATAT**TAGCTT** |
| AE009951.2 | p_0195 | 607113 | p_0195 | TTATATAAT | 5 | - | 1.0 | intergenic | TATTATATTTTTTACAAAATATTTGTTTTTTTATT**TGACAA** |
| AE009951.2 | p_0196 | 608361 | p_0196 | ACATATAAT | 4 | + | 0.98 | intergenic | CAGTATATGATTGTCAATCTAAACAAAATATTTTC**TTGAAA** |
| AE009951.2 | p_0197 | 608266 | p_0197 | AACTATTAA | 3 | - | 0.99 | intergenic | TTTAAATTTCAAGAAAATATTTTGTTTAGATTGAC**AATCAT** |
| AE009951.2 | p_0198 | 608842 | p_0198 | TGTTATAAT | 5 | + | 0.99 | intergenic | AAAATAGTATCTATGGCAGTAGATACTATTTTTAA**TTTTAA** |
| AE009951.2 | p_0199 | 614365 | p_0199 | TTTTTAACA | 2 | + | 0.78 | intergenic | AAATAGGGCTGTTGCAAATCAATAGTTATATTTGC**AATAAG** |
| AE009951.2 | p_0200 | 616422 | p_0200 | ATTTATAAA | 3 | + | 1.0 | intergenic | AGAAAAAGGATTAATTTTTCTAATAAAGTTAAAAT**TAATCC** |
| AE009951.2 | p_0201 | 616578 | p_0201 | ATATATAAT | 5 | + | 0.98 | intergenic | AAATAGAATAAATAGTTATAATAATAAGTTTTTTA**TTGATT** |
| AE009951.2 | p_0202 | 618259 | p_0202 | ATATATAAT | 5 | + | 1.0 | intergenic | AATTTAAAATTTTAGATTAAAGATACAGTTTTTTA**TTTATT** |
| AE009951.2 | p_0203 | 625012 | p_0203 | AGATAATAA | 3 | - | 1.0 | intergenic | TATAATAGTGATGATGGAATAAATTTTCTATCATC**ATTTTT** |
| AE009951.2 | p_0204 | 628378 | p_0204 | ATAAATAAT | 3 | - | 0.43 | intergenic | TTTAAATAAAATTTAGAGGGGATTATTTTTATAAT**CCCATT** |
| AE009951.2 | p_0205 | 628164 | p_0205 | GAATATAAT | 4 | - | 0.95 | intergenic | TTAAATATAATTATTTTATAATAGATTTTTACCAA**ATATTA** |
| AE009951.2 | p_0206 | 648115 | p_0206 | TGCTATAAT | 5 | + | 0.99 | intergenic | TTTTTTTATATATTCTTATTGTCAAGTTTATTGTC**TTGTCA** |
| AE009951.2 | p_0207 | 652328 | p_0207 | TGATATAAT | 5 | + | 1.0 | intergenic | GTATAGCAATTTTTTCCTAAAAATACAATTATAAT**TTATAA** |
| AE009951.2 | p_0208 | 652250 | p_0208 | AAAAAATTG | 2 | - | 1.0 | intergenic | ATTATATCACAAAATATTAAATATTATAAATTATA**ATTGTA** |
| AE009951.2 | p_0209 | 653742 | p_0209 | ATATATAAT | 5 | + | 1.0 | intergenic | AAGAGTTTTATAAAACAAACAATATGTTTGAAAAT**AATTGA** |
| AE009951.2 | p_0210 | 653609 | p_0210 | TGTTATAAT | 5 | - | 1.0 | intergenic | AAAACTCTTTACTTAACCATTTTGACTTTTAAATT**TCAGTA** |
| AE009951.2 | p_0211 | 654655 | p_0211 | TGATAAATT | 3 | + | 0.8 | intergenic | ACCAAGATATAGAGAAGATGAATATAATAAATTAT**TTAATA** |
| AE009951.2 | p_0212 | 654754 | p_0212 | ATAAAAATA | 3 | + | 1.0 | intergenic | GAAAATAATAAATACAAAAAAGTTTCAGGAAGTTT**TTATTT** |
| AE009951.2 | p_0213 | 656668 | p_0213 | TGCTATAAT | 5 | + | 0.9 | intergenic | ATATTTTGATTATATCATTATATAAAAATAAATAA**TAGAAA** |
| AE009951.2 | p_0214 | 656598 | p_0214 | TGATATAAT | 5 | - | 0.95 | intergenic | TAATTTTAATTATAGCAATTATAATGAGATATTTC**TATTAT** |
| AE009951.2 | p_0215 | 659742 | p_0215 | TAACTAAAA | 3 | + | 0.98 | intergenic | AAAAATATTACATTAATTTAAAGAAAGTATTTGAC**ATTATA** |
| AE009951.2 | p_0216 | 661302 | p_0216 | TGTAATAAA | 4 | + | 1.0 | intergenic | TTAATATGGATTTAAGATTAAAAAAGTCTTAAATC**CTTTTT** |
| AE009951.2 | p_0217 | 663002 | p_0217 | TCATATAAT | 4 | + | 1.0 | intergenic | ATTGTATAAAATAAAAAAAATATTTAATAATTTTT**ACCAAA** |
| AE009951.2 | p_0218 | 665053 | p_0218 | TAATATAAT | 5 | + | 1.0 | intergenic | TTTTTAAATTTAAAAAGTATATTACATAAATTTTT**TAGAAA** |
| AE009951.2 | p_0219 | 667237 | p_0219 | TATTATAAT | 5 | + | 1.0 | intergenic | TAAATGAAAAATTTTCAATATAATTTAAAATGATG**TTGAAA** |
| AE009951.2 | p_0220 | 667149 | p_0220 | TGCTAAAAT | 3 | - | 0.99 | intergenic | AATCTATTGTAATTTTCAACATCATTTTAAATTAT**ATTGAA** |
| AE009951.2 | p_0221 | 670045 | p_0221 | TTTTATTAT | 3 | - | 1.0 | intergenic | AGTTTTTTCACTCTTTCACCTCCTTTAAAAAGGGA**TGTTAG** |
| AE009951.2 | p_0222 | 675429 | p_0222 | TTAAAAATA | 3 | - | 1.0 | intergenic | CATAGGTGTTGTTTTTATTTTCTCTTAAAAAATAA**GAAAAA** |
| AE009951.2 | p_0223 | 676241 | p_0223 | TGATATAAT | 5 | - | 1.0 | intergenic | ACATCTATGTCTGTGGATGTTGTTTTTATTTTTAT**ATTGAT** |
| AE009951.2 | p_0224 | 678179 | p_0224 | TGTTAAAAT | 4 | - | 1.0 | intergenic | GTTTAGCGACTCCATTTTTTATTATATAAAAGTAA**TTGTTT** |
| AE009951.2 | p_0225 | 682300 | p_0225 | TGATAAAAT | 4 | + | 1.0 | intergenic | AAAAATTAATAATTTATATTATAACATATTTTATG**GTTACA** |
| AE009951.2 | p_0226 | 682247 | p_0226 | ATAAAATAT | 3 | - | 1.0 | intergenic | ACCTTTTTATTTAATTATCTTTTTTATTTTATCAT**TTTTTT** |
| AE009951.2 | p_0227 | 684547 | p_0227 | AATATTAAT | 3 | + | 0.99 | intergenic | TGCTTCAATTTTTCATCTAAAATTTAGAATACAAT**TTCACT** |
| AE009951.2 | p_0228 | 686252 | p_0228 | CTTTATAAT | 4 | + | 1.0 | intergenic | ATAAATCAAGTAATTTTTAGGCTTAAATTTGACAA**GTAACA** |
| AE009951.2 | p_0229 | 686434 | p_0229 | AATTATTAA | 3 | + | 0.95 | intergenic | AACAAACTATTAAAAAAAACTTAAACAATACTAAA**AACAAA** |
| AE009951.2 | p_0230 | 686343 | p_0230 | TATTTTTAT | 3 | - | 0.98 | intergenic | TTAATCTGATTTTGTTTTTAGTATTGTTTAAGTTT**TTTTTA** |

| | | | | | | | | | |
|---|---|---|---|---|---|---|---|---|---|
| AE009951.2 | p_0231 | 686244 | p_0231 | AAGATAAAA | 3 | - | 1.0 | intergenic | GTTCTAAATTTTGCTCTTAAGAATTATCTCCAATT**TTGTCC** |
| AE009951.2 | p_0232 | 687102 | p_0232 | TGAAATATT | 3 | + | 0.99 | intergenic | AACATCATTTTTTTTCTGTAAATATAAATAATATT**CATTGA** |
| AE009951.2 | p_0233 | 688957 | p_0233 | TGGTATAAT | 5 | + | 0.99 | intergenic | TGTTGTAAGTAATATTTTCAACAACTTCTTTTTTA**TTGTTA** |
| AE009951.2 | p_0234 | 692274 | p_0234 | TGATATAAT | 5 | + | 0.46 | intergenic | TAGTACTTGCTAGGCAGGTACTATTTTTCTTTTTG**TTGTTT** |
| AE009951.2 | p_0235 | 697125 | p_0235 | TGTTATAAT | 5 | + | 1.0 | intergenic | TTATTTATTGATTGATTTACAATAACCTTATTTTT**TAAATA** |
| AE009951.2 | p_0236 | 708909 | p_0236 | TTCGCTAAA | 2 | + | 1.0 | intergenic | TAGTTCGTTACTAGCCAGATTTCTTAACAATAAAA**AATCAA** |
| AE009951.2 | p_0237 | 709004 | p_0237 | TTATATAAT | 5 | + | 0.68 | intergenic | CGGCTCACTCTATTTGATTTTTTATTTAAAATCTG**GAATGT** |
| AE009951.2 | p_0238 | 711802 | p_0238 | AGCATTTAA | 2 | + | 0.25 | intergenic | AAAAAGCAGAACTCACTTCGTTCAAACAACTGCTT**TTTTTG** |
| AE009951.2 | p_0239 | 723404 | p_0239 | AAAATAAGG | 2 | + | 1.0 | intergenic | CCTAAAATCTGGAATGTAACTCACTTATTTTTGTA**AGTTTT** |
| AE009951.2 | p_0240 | 729265 | p_0240 | TCTTATAAT | 4 | + | 0.77 | intergenic | GATTTCAGTAGCTTTTTATTTATGTGACTATGTCA**CTGAAA** |
| AE009951.2 | p_0241 | 736003 | p_0241 | AAATATAAT | 5 | + | 0.99 | intergenic | ACTATAAAAGAAATATTTTATAAATAATATTTAAC**AATAAA** |
| AE009951.2 | p_0242 | 736080 | p_0242 | TTAGATAAT | 3 | + | 1.0 | intergenic | TAAAAGTAAAATATATTTTTTATATATTAAACAAA**AAAGAT** |
| AE009951.2 | p_0243 | 735913 | p_0243 | TTTTTTTAT | 3 | - | 1.0 | intergenic | ATTTAGCTTTTTTTATTGTTAAATATTATTTATAA**AATATT** |
| AE009951.2 | p_0244 | 740129 | p_0244 | TTCTCTAAA | 2 | + | 0.99 | intergenic | ATCTTCACTACTTTTGATTTTTAATTCTTCAAGCC**AGCCTT** |
| AE009951.2 | p_0245 | 741036 | p_0245 | CTTTATAAT | 4 | + | 1.0 | intergenic | TTTAATCCCCATCTGTTTTTTATTATAAGGAAAAT**ATATTT** |
| AE009951.2 | p_0246 | 741601 | p_0246 | ACCTATAAT | 4 | + | 1.0 | intergenic | ATAGGAAAGTAAAAGTAATCTAATAAAAAATTCTA**TTGACA** |
| AE009951.2 | p_0247 | 746666 | p_0247 | TTTTATATA | 3 | + | 1.0 | intergenic | AAATATATAGAAAAATTTTTTTAAGAAATTTTGAA**AAAATA** |
| AE009951.2 | p_0248 | 746575 | p_0248 | AAATATAAT | 5 | - | 1.0 | intergenic | TTTTGTCAACTATTTTTTCAAAATTTCTTAAAAAA**ATTTTT** |
| AE009951.2 | p_0249 | 750361 | p_0249 | TCTGATTTT | 2 | + | 0.78 | intergenic | GGATAAAAATTAAGAATTCGCTGCAAAACAGAGAA**ACTCGC** |
| AE009951.2 | p_0250 | 753587 | p_0250 | ATTTTATAT | 3 | + | 0.88 | intergenic | AAAATTTTAAATATTAAAAAACTAGTGTTATTTTA**TATAGA** |
| AE009951.2 | p_0251 | 753711 | p_0251 | TTATATAAT | 5 | + | 0.06 | intergenic | TCTTAAATAATTCAACACTTTATATAAAATATAAT**TTGAAT** |
| AE009951.2 | p_0252 | 753612 | p_0252 | TATTAATAA | 3 | - | 0.85 | intergenic | TAATTCAAATTATATTTTATATAAAGTGTTGAATT**ATTTAA** |
| AE009951.2 | p_0253 | 753512 | p_0253 | ATTTAAAAT | 3 | - | 1.0 | intergenic | CAAATATAAAATATTTCTTTAAAATTTCTATATAA**AATAAC** |
| AE009951.2 | p_0254 | 756520 | p_0254 | TAAAATAAT | 3 | + | 1.0 | intergenic | ATAAAAATTGTAACATTTGTTACAGAAAAAACTTT**ACAAGA** |
| AE009951.2 | p_0255 | 756423 | p_0255 | TAAAATATA | 3 | - | 1.0 | intergenic | TAATTCTTGTAAAGTTTTTTCTGTAACAAATGTTA**CAATTT** |
| AE009951.2 | p_0256 | 758604 | p_0256 | ATCTAAAAT | 3 | + | 0.55 | intergenic | ACAAGTCAGCTTCAAACACACCGAGATTTGCTCGG**CTCACT** |
| AE009951.2 | p_0257 | 759649 | p_0257 | ATATATAAT | 5 | + | 1.0 | intergenic | CTTTTTTGTTATTAATAAAAAAATATAAAAAGTAC**TTGATT** |
| AE009951.2 | p_0258 | 761557 | p_0258 | TGATAATAT | 4 | + | 0.84 | intergenic | TAAATGAGTTTAGCGAATATAATATTGATGAAAAG**GGACAA** |
| AE009951.2 | p_0259 | 761843 | p_0259 | GGGAATATA | 2 | + | 0.67 | intergenic | AGCAGGTAAAGTTAATTTAAAAGATGGAGATTTCA**TTGGAA** |
| AE009951.2 | p_0260 | 770280 | p_0260 | TGTATCATA | 2 | + | 1.0 | intergenic | GGAAAAAGCTAAAAAAATAAAGAGAATTTTTTTAAA**GAAGAT** |
| AE009951.2 | p_0261 | 773388 | p_0261 | AATTATAAT | 5 | + | 0.98 | intergenic | GGTTGTATAGATAAATATTTTATATATTTATCATA**CAAGGA** |
| AE009951.2 | p_0262 | 773270 | p_0262 | TTATATAAT | 5 | - | 1.0 | intergenic | TATAAAATATTTATCTATACAACCTCCTTTCATAG**TTTATT** |
| AE009951.2 | p_0263 | 790034 | p_0263 | AAAATATTA | 3 | + | 0.93 | intergenic | GGTTATAACTTATAATCCAATTTTTTTCTTTTCAC**TAGACA** |
| AE009951.2 | p_0264 | 793549 | p_0264 | TGATTTAAT | 3 | + | 0.98 | intergenic | ATATGATTAATTATTGAGCTATTGTAAAAATAAAT**TTGCAA** |
| AE009951.2 | p_0265 | 794637 | p_0265 | AGACAAAAA | 2 | + | 1.0 | intergenic | TTTACTTTTTCAGATGATAAAATTTTAAATTTTGC**CTGTGT** |
| AE009951.2 | p_0266 | 796904 | p_0266 | ATTTTTAAA | 3 | + | 1.0 | intergenic | GCTTATTTTCTATTTAATATCAATAAATTTTTATC**ACAAAA** |
| AE009951.2 | p_0267 | 797016 | p_0267 | ATAATGATA | 2 | + | 1.0 | intergenic | TTCTCTCCCCCCAGAAAGAAAAAAAGCCCTCCTTG**AAAAAG** |
| AE009951.2 | p_0268 | 797119 | p_0268 | TGATAAAAT | 4 | + | 1.0 | intergenic | TTAAAAAGTCCACCTCAAAAACGGACTTTTTTTTA**TTTTCA** |
| AE009951.2 | p_0269 | 805436 | p_0269 | ATGTATAAT | 4 | - | 0.99 | intergenic | GAGGAGATTTTTTATTATTATATATTAGATACTAC**TTGAAA** |
| AE009951.2 | p_0270 | 810795 | p_0270 | AAATATAAT | 5 | + | 0.99 | intergenic | ATAATTAAATTCAAGTTTTTTATAAATAGATTCTA**TTATAT** |
| AE009951.2 | p_0271 | 810977 | p_0271 | TTTAAAATA | 3 | + | 0.54 | intergenic | TTTAAAAGTTATTTAAATATTTTTTAAGAAAAATT**TGTGAT** |
| AE009951.2 | p_0272 | 810706 | p_0272 | TTTTATTTA | 3 | - | 1.0 | intergenic | TTTTTTTTACAAATATAATAGAATCTATTTATAAA**AAACTT** |
| AE009951.2 | p_0273 | 815865 | p_0273 | TAAAAATAA | 3 | + | 1.0 | intergenic | GTTTTAAACTGGGTTCTTTATTTTTTATAAGAAAA**TGAAAA** |
| AE009951.2 | p_0274 | 817441 | p_0274 | CATTATAAT | 4 | + | 0.98 | intergenic | TTCTTTATTTTTTATAAAATACATAAATTTGTAAT**TGACAT** |
| AE009951.2 | p_0275 | 823792 | p_0275 | TAAATATTT | 3 | + | 0.99 | intergenic | ATAAAAAAAAGCTAGAAAAATCTATAAAAAAAATTG**TATTAT** |
| AE009951.2 | p_0276 | 837121 | p_0276 | TGTTACAAT | 2 | + | 1.0 | intergenic | AGGTGCAGTACAAAATTCTTTCTTATAAAAAATAAG**TTGAAA** |
| AE009951.2 | p_0278 | 845311 | p_0278 | TTGTATAAT | 4 | + | 1.0 | intergenic | TTCAAGTCATTTTTTTTTACATTTTATTCCAAAAAA**TTTAAA** |
| AE009951.2 | p_0279 | 845254 | p_0279 | TGGAATAAA | 3 | - | 0.99 | intergenic | TTGTTTAATTTCTTTAATATTATTATACAATTTTT**TTATTC** |
| AE009951.2 | p_0280 | 846901 | p_0280 | TGTTACAAT | 2 | + | 1.0 | intergenic | AGAACTAGGAGTATTTTTTGCATTGGAATTTTTCT**TTTCTT** |
| AE009951.2 | p_0281 | 848520 | p_0281 | TATATAAAA | 3 | + | 1.0 | intergenic | AATATATAAAAATTAAATTTTAAGAAAAACTTATT**TAAAAA** |
| AE009951.2 | p_0282 | 856322 | p_0282 | TGTATATTT | 3 | - | 1.0 | intergenic | TTTTATTACTATTTTGTATCAGTCTATTTTTTTATT**TGACTT** |
| AE009951.2 | p_0283 | 857721 | p_0283 | TAAAAAATG | 3 | - | 0.02 | intergenic | TGTGATTCAAAAACTCTATATAAAAGAATAATTGG**AACAAA** |
| AE009951.2 | p_0284 | 857583 | p_0284 | CTAAATTTT | 3 | - | 1.0 | intergenic | GAGGCAGGACTAGCAAGTTTTTGTGATGCTTCTGT**TGTAGA** |
| AE009951.2 | p_0285 | 859642 | p_0285 | TACTTTATT | 3 | - | 1.0 | intergenic | TGCAGTTCATTTCTGACAGCTTCTTTTCTATTTAT**CTTTTT** |
| AE009951.2 | p_0286 | 863507 | p_0286 | AACTATAAT | 4 | + | 0.07 | intergenic | AGAAAATCAATATTTAAAAAATATTAATATTTTTT**ACCAAA** |
| AE009951.2 | p_0287 | 863419 | p_0287 | TATAAAAAA | 3 | - | 0.98 | intergenic | GTAATTATTAAGATTTGGTAAAAAATATTAATATT**TTTTAA** |
| AE009951.2 | p_0288 | 864008 | p_0288 | TGGTATAAT | 5 | + | 1.0 | intergenic | GTATTTAGCCATCAAAATAACTTCTCAAAAATTAC**CTCATT** |
| AE009951.2 | p_0289 | 868188 | p_0289 | GAGTATAAT | 4 | + | 1.0 | intergenic | TTTAAGAATAAAAGTTTAGTATTTTAGTAAAAAAT**TTTGAC** |
| AE009951.2 | p_0290 | 872741 | p_0290 | GGGTATAAT | 4 | + | 1.0 | intergenic | TAGCTTAAATATTTGATATATATTTTATAATTTTT**TAGTGT** |
| AE009951.2 | p_0291 | 881597 | p_0291 | TGATACAAT | 2 | + | 0.72 | intergenic | CCATTTTTATATAAGTATATTTTAAACTTTCTATC**TTTACT** |
| AE009951.2 | p_0292 | 882314 | p_0292 | TGTTATAAT | 5 | + | 1.0 | intergenic | CAAGAGTTAGAGAAAATAATGATATGATTTCATGG**ATAAAA** |

| | | | | | | | | | |
|---|---|---|---|---|---|---|---|---|---|
| AE009951.2 | p_0293 | 882619 | p_0293 | TATTATAAA | 3 | + | 1.0 | intergenic | ATAGATTTTTTTTAATATAAATTTCTAATAAAAAT**GATTGA** |
| AE009951.2 | p_0294 | 886771 | p_0294 | AATTATAAT | 5 | - | 1.0 | intergenic | ATTTTAATTCTTTATTAAAAAAATAAAAAAGTACT**TGACAG** |
| AE009951.2 | p_0295 | 889986 | p_0295 | TATTAATAT | 3 | + | 0.96 | intergenic | ATTTATATAATAAAATTTTTTGTTAATAAACTATA**TTGATA** |
| AE009951.2 | p_0296 | 890123 | p_0296 | AAAAATAAA | 3 | + | 1.0 | intergenic | TTAATAAATAATAAAATTGATAAAAAAAATACTCTA**AATAAG** |
| AE009951.2 | p_0297 | 890010 | p_0297 | TTAAATCAT | 2 | - | 1.0 | intergenic | ATTTTTTTATCAATTTTATTATTTATTAAAATTTT**AATTGA** |
| AE009951.2 | p_0298 | 889911 | p_0298 | TATTATATA | 3 | - | 0.87 | intergenic | AATATATTAATAAAATAAATTTTTATTATCAATAT**AGTTTA** |
| AE009951.2 | p_0299 | 890987 | p_0299 | TATTATAAT | 5 | + | 0.99 | intergenic | AAAACATTGTAATTTTTTAAAATTTTATGTATAAT**TGTCAA** |
| AE009951.2 | p_0300 | 891484 | p_0300 | ATTTATAAA | 3 | + | 1.0 | intergenic | ATGAAATAAAACCTTTTAAGTTTATATTTTTTTAAT**ATAAAT** |
| AE009951.2 | p_0301 | 897888 | p_0301 | TGATATAAT | 5 | + | 1.0 | intergenic | TGAATAAAAGTGAAAATTTTGAAAGCTCTTTTTTA**AATTGA** |
| AE009951.2 | p_0302 | 901752 | p_0302 | TTATATAAT | 5 | - | 1.0 | intergenic | GCTTTTACAAGTAAAATTTGTAAAAGCTTTTTTAT**ATTTTA** |
| AE009951.2 | p_0303 | 915884 | p_0303 | TGTTAAAAT | 4 | - | 0.96 | intergenic | AATCTTAATTAAAGGTTTGAGTGTAGTTTTTTATC**TTGATA** |
| AE009951.2 | p_0304 | 919195 | p_0304 | TGATATATT | 3 | - | 0.99 | intergenic | CTGTTGAAAATGCAAAAAATATAAAGGAATTAACA**GATTTT** |
| AE009951.2 | p_0305 | 919096 | p_0305 | TGCTGGATT | 1 | - | 1.0 | intergenic | ATGGGGCAAAAGCAACAATAGCAGTAGCAGATACA**AACTAT** |
| AE009951.2 | p_0312 | 929927 | p_0312 | TTCTATTAA | 3 | + | 1.0 | intergenic | ATTTCATTAAAATTTTGTTTCAATTTTTTCACTTC**CTTTTC** |
| AE009951.2 | p_0313 | 929846 | p_0313 | TGAAATCAA | 2 | - | 0.86 | intergenic | ATAGAAAAATTAAATATTTCGAAAAGGAAGTGAAA**AAATTG** |
| AE009951.2 | p_0314 | 934445 | p_0314 | TACAATAAT | 3 | + | 1.0 | intergenic | ACCGTTTTTTTATTGTTAAAAAACTATTTTACTTG**TAAAAA** |
| AE009951.2 | p_0315 | 943121 | p_0315 | TGATATGTT | 2 | + | 1.0 | intergenic | AAGTGGTTAAAAAAAGGTCAATTTAATCTTATAAA**TTGACT** |
| AE009951.2 | p_0316 | 947247 | p_0316 | ATTTAAAAT | 3 | + | 0.99 | intergenic | AAGTGCTACAATGATTGAAGATATAAAAATTTTTT**TATAGA** |
| AE009951.2 | p_0317 | 950034 | p_0317 | ATAATAAAA | 3 | + | 0.6 | intergenic | ATTATTCAATAACTTTTTTATTTATTTGTGCTAAC**ATATAT** |
| AE009951.2 | p_0318 | 957039 | p_0318 | TACTATATT | 3 | + | 0.99 | intergenic | TGTTTAAAAGAAGTATAACATTAATTTTATTTATT**GACAAA** |
| AE009951.2 | p_0319 | 956974 | p_0319 | AATGTTATA | 2 | - | 0.99 | intergenic | AATCTTTACTTATAATATAGTATATCATTTAATTT**CTTTGT** |
| AE009951.2 | p_0320 | 957723 | p_0320 | TGGTATATA | 3 | + | 1.0 | intergenic | AAGTCTAACCAATCATTTTTTACTTTTATTTTTTA**TTGACA** |
| AE009951.2 | p_0321 | 959686 | p_0321 | AAAAAATAT | 3 | + | 1.0 | intergenic | TTGATACAAATTCAAGCAATTTAAGAAAATTGAAA**AATATC** |
| AE009951.2 | p_0322 | 962334 | p_0322 | AAAATAATG | 3 | - | 0.99 | intergenic | CTTGACAAGTTCTAAAAAATACTATATACTTATAT**GGTATT** |
| AE009951.2 | p_0323 | 963889 | p_0323 | TTTTAAAAT | 3 | + | 0.96 | intergenic | AGGTCTTTATTAAGACCTCTGAAATTAAAAAAGAA**TTATAC** |
| AE009951.2 | p_0324 | 964200 | p_0324 | TAATATAAT | 5 | + | 1.0 | intergenic | AATAAGAAAAATTTAGATAACATAAATTTTATTTA**TTGACA** |
| AE009951.2 | p_0325 | 964095 | p_0325 | TGATAATAT | 4 | - | 1.0 | intergenic | AATAAATAAAATTTATGTTATCTAAATTTTTCTTA**TTGACA** |
| AE009951.2 | p_0326 | 965212 | p_0326 | TGATATAAT | 5 | + | 1.0 | intergenic | ATATTTCCCACAGAATTTTCTGTGGGATTTTTATT**TTAAAA** |
| AE009951.2 | p_0327 | 969165 | p_0327 | TATTAAAAT | 3 | - | 1.0 | intergenic | TAAAGCTTTAGACTAAGAGAAAGTGCTGATTTTAC**TCAAAC** |
| AE009951.2 | p_0328 | 971093 | p_0328 | TGTTAAAAT | 4 | - | 1.0 | intergenic | TCTAAAAAATATCAAATTAGAATTTTTTTTTATAA**TTGAAC** |
| AE009951.2 | p_0329 | 971659 | p_0329 | TACTTTTTA | 2 | + | 1.0 | intergenic | TTAATGATTTTTTATTATATAATATTTTTTTATTT**TTTACA** |
| AE009951.2 | p_0330 | 971594 | p_0330 | TTATATAAT | 5 | - | 1.0 | intergenic | TATCACAAATTTTTAAAAAGTATACCTGATAAAAT**TTGTAA** |
| AE009951.2 | p_0331 | 981845 | p_0331 | TGATATAAT | 5 | - | 1.0 | intergenic | ATATATAAAGTTCACTTATAAAAACAAATTAATGT**TTTGGA** |
| AE009951.2 | p_0332 | 985203 | p_0332 | TGTTAAAAT | 4 | + | 0.85 | intergenic | TACGGGGTCATCCTTTAAGATTATACAATATTTTA**TTTAAG** |
| AE009951.2 | p_0333 | 987491 | p_0333 | TGTTAAAAT | 4 | - | 1.0 | intergenic | GAGGCTATTATTTGAAGTAGCCTCATTTTTTAATA**TTGACA** |
| AE009951.2 | p_0334 | 997045 | p_0334 | ATAAAAATT | 3 | - | 1.0 | intergenic | AATTAAAAAAAATAATACAAATAAAAAATAAAAGA**GTTACA** |
| AE009951.2 | p_0335 | 1000038 | p_0335 | TTATATAAT | 5 | + | 1.0 | intergenic | TACCATACTTTTTCACTTTAAATTATATTTTATTG**ATAAAA** |
| AE009951.2 | p_0336 | 1000051 | p_0336 | TAAAAATAT | 3 | - | 1.0 | intergenic | CAATATTTTATACTTTATTACTATAACATATATTT**TTAATT** |
| AE009951.2 | p_0337 | 1002646 | p_0337 | AATTAAAAA | 3 | + | 1.0 | intergenic | ATTATATTAAATTTTGAAGAAAATTTCTAAAAAAT**TTATGT** |
| AE009951.2 | p_0338 | 1002568 | p_0338 | TAATATAAT | 5 | - | 1.0 | intergenic | TTTTTAATTTAAAAAAGTATATTACATAAATTTTT**TAGAAA** |
| AE009951.2 | p_0339 | 1010359 | p_0339 | TCATATAAT | 4 | + | 1.0 | intergenic | TTTTATTTCCTCCTTTAATATAAATTGTTTAATTT**TTTGAC** |
| AE009951.2 | p_0340 | 1010571 | p_0340 | TGCTAAAAT | 3 | + | 1.0 | intergenic | TTGAATTAAATTTTTCATAATTTTTAAAAAAAAAC**TTGAAA** |
| AE009951.2 | p_0341 | 1010501 | p_0341 | TGAAAAATT | 3 | - | 1.0 | intergenic | ACTTTATAATTTTAGCAAACACATGTTATTTTTTC**AAGTTT** |
| AE009951.2 | p_0342 | 1027604 | p_0342 | TTTTAAAAT | 3 | + | 1.0 | intergenic | ATTATACTCCTTATAAAAAAATATTTCTATATAAA**AGTTTT** |
| AE009951.2 | p_0343 | 1027580 | p_0343 | TTTTAAAAA | 3 | - | 1.0 | intergenic | TTATTATTTATAATAACATATTTTTATTTTAAAAA**ACAAAC** |
| AE009951.2 | p_0344 | 1028723 | p_0344 | TGTTATAAT | 5 | + | 0.91 | intergenic | GAAACCTACTTTAAATATTTATAATTTCATTATTA**TAGAAA** |
| AE009951.2 | p_0345 | 1028583 | p_0345 | TGGTAAAAT | 3 | - | 0.96 | intergenic | TCTATTTGGTAACTAAAATTCAAAAAAGTGCCTTC**TTAAAA** |
| AE009951.2 | p_0346 | 1034297 | p_0346 | TAATATAAT | 5 | + | 0.99 | intergenic | ATATATTTTTTATATGTATATTAGAGATAATTAAT**TTGACA** |
| AE009951.2 | p_0347 | 1035950 | p_0347 | TTAATAAAA | 3 | + | 0.87 | intergenic | AGTGTTTAGATTTTAAAATTTAAAAGAGCTAGAAG**ACTTCT** |
| AE009951.2 | p_0348 | 1042396 | p_0348 | TAATATAAT | 5 | - | 1.0 | intergenic | TTTTTAATTTAAAAAAGTATATTACATAAATTTTT**TAGAAA** |
| AE009951.2 | p_0349 | 1045355 | p_0349 | TGTAAAATA | 4 | + | 1.0 | intergenic | GATATTTTTAATAAATTATAACATAAAAATAAAAA**AAATAA** |
| AE009951.2 | p_0350 | 1045367 | p_0350 | TTTTTATAT | 3 | - | 1.0 | intergenic | TTCTAAGTTTTTTCAATATTTTTATTTTTTTTTAT**CTATTT** |
| AE009951.2 | p_0351 | 1050109 | p_0351 | TATTATAAT | 5 | + | 0.97 | intergenic | TTTTTCAATGGTAAGATAAAGTTATTCTTTTATTA**TTCAAA** |
| AE009951.2 | p_0352 | 1050011 | p_0352 | AAAATTATG | 2 | - | 1.0 | intergenic | TTTTTTGAATAATAAAAGAATAACTTTATCTTACC**ATTGAA** |
| AE009951.2 | p_0353 | 1054400 | p_0353 | AGATAATTT | 3 | + | 0.8 | intergenic | CAGCTAAAGAAGAGTCAAAAAAATTGGGGTAATACA**GAAAAT** |
| AE009951.2 | p_0354 | 1071045 | p_0354 | GTTTATAAT | 4 | + | 0.99 | intergenic | ATTACTTTTTTTTCTAATGATATAATTTATTAAAA**ATTATA** |
| AE009951.2 | p_0355 | 1082793 | p_0355 | AAGAAAATA | 3 | + | 1.0 | intergenic | AATTTTAACAATGACTGAAAATGAATTAAAAAATA**AGAATT** |
| AE009951.2 | p_0356 | 1083148 | p_0356 | GTACAATAA | 2 | + | 1.0 | intergenic | ATTCTTATAAACACTTTATTCCTAGTTGGAAACTT**GACAAT** |
| AE009951.2 | p_0357 | 1095642 | p_0357 | AACTATAAT | 4 | + | 1.0 | intergenic | AAAAATTCTCTTATTTTTTATAACAAAAAATTTTA**AAAACT** |
| AE009951.2 | p_0358 | 1095723 | p_0358 | GAATTATCG | 1 | + | 1.0 | intergenic | ATTTATGTAAAAGAAGCGCCAGAACTCTTTTTAGA**GTTGAC** |
| AE009951.2 | p_0359 | 1099563 | p_0359 | TGCTATAAT | 5 | + | 1.0 | intergenic | ACTGTTTTTTACTTCATTTTTTATTTATAGCAATC**TATTAT** |
| AE009951.2 | p_0360 | 1101274 | p_0360 | TGATAAAAT | 4 | + | 1.0 | intergenic | ATAGTATTTTTTATTTTTTTTCAAAATAATTTAAT**TTGACA** |

| | | | | | | | | | |
|---|---|---|---|---|---|---|---|---|---|
| AE009951.2 | p_0361 | 1101966 | p_0361 | TGGTATAAT | 5 | + | 1.0 | intergenic | AGAAAGACACTGAATTACAAAAAAAGTTTTTTAGT**GTCTTT** |
| AE009951.2 | p_0362 | 1107322 | p_0362 | CTTTAAAAT | 3 | + | 0.99 | intergenic | TACAATAGCTTTTTTCTTAAAAATGTAGTTGTAAA**TCTTTA** |
| AE009951.2 | p_0363 | 1114761 | p_0363 | TTTTTATAT | 3 | + | 1.0 | intergenic | ACCCCTTTCTTATATATTTTTGACTAAAAGAATAA**AAAAAG** |
| AE009951.2 | p_0364 | 1116513 | p_0364 | ATTTATTTT | 3 | + | 1.0 | intergenic | AATTAGAGAATTTTTTTGAGAATAAAATCTTAAAA**AAATTC** |
| AE009951.2 | p_0365 | 1118394 | p_0365 | TTATATAAT | 5 | + | 0.98 | intergenic | AAAGTTATGTAAATTTTTTACTTATATTTTATATA**TTGACA** |
| AE009951.2 | p_0366 | 1118284 | p_0366 | TAAAATTAT | 3 | - | 1.0 | intergenic | ATAAAATATAAGTAAAAAATTTACATAACTTTAAA**AATACT** |
| AE009951.2 | p_0367 | 1122215 | p_0367 | TGTTATAAT | 5 | + | 0.97 | intergenic | TACTATATTGAATTAATAAATAAAAACCTTCTCAC**TTTTAT** |
| AE009951.2 | p_0368 | 1128395 | p_0368 | TGCAAATAT | 3 | + | 1.0 | intergenic | AATGAGGGATAAAAAATTCTCTCATTTTTTAATTA**ATTTTA** |
| AE009951.2 | p_0369 | 1132996 | p_0369 | ATATATAAA | 3 | + | 1.0 | intergenic | TATAACATAAAAAAAATAAAAAAAAAGAACAAATA**TTTATA** |
| AE009951.2 | p_0370 | 1133181 | p_0370 | TGATAAAAT | 4 | + | 1.0 | intergenic | TACATATTGTATTGAAAAAATAAAAAAATTTTTAC**ATTGAC** |
| AE009951.2 | p_0371 | 1133096 | p_0371 | TAGTATAAT | 4 | - | 1.0 | intergenic | CATATTTTTTTATATGTCAATGTAAAAATTTTTT**TATTTT** |
| AE009951.2 | p_0372 | 1137440 | p_0372 | TGTTATAAT | 5 | - | 1.0 | intergenic | TACAAATACTTTCTTTAAAAATTTAATAATATTTA**TCATTA** |
| AE009951.2 | p_0373 | 1138764 | p_0373 | TTATATAAT | 5 | + | 1.0 | intergenic | CAAAAAAATTTTTTTAAAAAAATAAAAATATTTAC**TTTTAA** |
| AE009951.2 | p_0374 | 1138760 | p_0374 | TACAATTTT | 3 | - | 1.0 | intergenic | TATATAATTTATTTTAAAATTTTTTTAAATTCATT**TCAATA** |
| AE009951.2 | p_0375 | 1141016 | p_0375 | TGATATAAT | 5 | + | 1.0 | intergenic | ATTAGTTTCTGTAATTAGAAGTTAATCCTTTTTTA**TTGAGA** |
| AE009951.2 | p_0376 | 1148687 | p_0376 | TGTTATAAT | 5 | + | 1.0 | intergenic | ATCAATACTATAAAAGAATATTTGTCCCATTACTT**TAAATT** |
| AE009951.2 | p_0377 | 1148541 | p_0377 | TTTACTATT | 2 | - | 1.0 | intergenic | AACATAAAAATATTAAAATTCCTAAAAAAATATAT**TGACAA** |
| AE009951.2 | p_0378 | 1152073 | p_0378 | AATTATAAA | 3 | - | 1.0 | intergenic | AGCTATTTTCATAGCTCTTTTTTATTTATTTTTTT**ACTTGA** |
| AE009951.2 | p_0379 | 1157820 | p_0379 | TGTTATAAT | 5 | - | 1.0 | intergenic | TTTCCGTACTTTTCTATAAAAATAAATTTTAAAAC**CTTGAA** |
| AE009951.2 | p_0380 | 1158838 | p_0380 | TAAATAATT | 3 | - | 1.0 | intergenic | AATTCTTATTTAAATTTTTTTAAAAAAGTTCTTGA**CTTTTC** |
| AE009951.2 | p_0381 | 1160419 | p_0381 | TTATATATT | 3 | + | 0.2 | intergenic | AAATAATAAATAAAATAAACTATGAAAGGAGGTTG**TATAGA** |
| AE009951.2 | p_0382 | 1160331 | p_0382 | TTATATAAT | 5 | - | 1.0 | intergenic | TATAAAATATTTATCTATACAACCTCCTTTCATAG**TTTATT** |
| AE009951.2 | p_0383 | 1162742 | p_0383 | ATCTATTAT | 3 | - | 0.97 | intergenic | AATTTGTACAGCTTTTTTAAATTTTTTTAAGAAAG**CTATTG** |
| AE009951.2 | p_0384 | 1163425 | p_0384 | ATATATAAT | 5 | + | 0.99 | intergenic | ATTATAACTACAAATTTGATTTTTTGCAAGTTTTA**TTTATA** |
| AE009951.2 | p_0385 | 1163450 | p_0385 | ATAATTATT | 3 | - | 0.23 | intergenic | TATTGTTTTGGTGCTATGATACTACCACTTTTTAG**TAATAT** |
| AE009951.2 | p_0386 | 1163347 | p_0386 | AGTTATAAT | 4 | - | 1.0 | intergenic | ATTATATATTTTTTTCTATTTATTATAAATAAAAC**TTGCAA** |
| AE009951.2 | p_0387 | 1169627 | p_0387 | TATTATAAT | 5 | - | 0.93 | intergenic | AAATAGGAAAATTATAAAAAAATATAATCTTCCTA**TTTTTT** |
| AE009951.2 | p_0388 | 1176013 | p_0388 | AATTAAAAA | 3 | - | 0.97 | intergenic | CTGAAAACTTAGAAAATATTGCAGAAAAGAATGGA**GCTTTA** |
| AE009951.2 | p_0389 | 1175883 | p_0389 | AGACTTAAA | 2 | - | 0.98 | intergenic | TTTAATAAATACTTGGAATATCTCTGAAATTGAAA**AATTTT** |
| AE009951.2 | p_0390 | 1184400 | p_0390 | TTTTAAAAA | 3 | + | 1.0 | intergenic | AATCAATTTTAAAAAGTCCTCTCTTTATTTTTTAA**AAGAAC** |
| AE009951.2 | p_0391 | 1184330 | p_0391 | ACTTTTTAA | 3 | - | 1.0 | intergenic | ATTTTTTTTATTTTTTTTTAAAATTTAAACAGTTC**TTTTAA** |
| AE009951.2 | p_0392 | 1186458 | p_0392 | TAATATAAT | 5 | - | 1.0 | intergenic | TTTTTAATTTAAAAAAGTATATTACATAAATTTTT**TAGAAA** |
| AE009951.2 | p_0393 | 1188864 | p_0393 | AAGCTATAA | 2 | + | 0.84 | intergenic | ATAAAAAATATATTTGGTATATAGAAGTAAATTTT**TTATTG** |
| AE009951.2 | p_0394 | 1188778 | p_0394 | TGTTATAAT | 5 | - | 1.0 | intergenic | TTATTAAAAAAAATGTCAATAAAAAATTTACTTCTA**TATACC** |
| AE009951.2 | p_0395 | 1191959 | p_0395 | TGTTATAAT | 5 | + | 1.0 | intergenic | TTTTTCATATATAAATCAAGTTTATTTATTATTTA**TTGAAA** |
| AE009951.2 | p_0396 | 1191870 | p_0396 | TTATATAAT | 5 | - | 1.0 | intergenic | TTAAACATCTTTTTTCAATAAATAATAAATAAACT**TGATTT** |
| AE009951.2 | p_0397 | 1194139 | p_0397 | TAGTATAAT | 4 | + | 1.0 | intergenic | TAATAATATACACTTTTTTAATTTAAAAAACAAGT**TTGCAT** |
| AE009951.2 | p_0398 | 1194234 | p_0398 | TGTTATAAT | 5 | + | 0.72 | intergenic | CATTTTTTAGAATAATTCTTTATAGATAAGATTAA**TTCACA** |
| AE009951.2 | p_0399 | 1194064 | p_0399 | TGTATATTA | 3 | - | 0.98 | intergenic | TTTATTATACTATATCAATAAAGAAAATGCAAACT**TGTTTT** |
| AE009951.2 | p_0400 | 1202629 | p_0400 | TTTTATAAT | 5 | + | 0.98 | intergenic | AACATTACTCCAAAAGGTGACAGAAACTCATTCTA**TTGACA** |
| AE009951.2 | p_0401 | 1202748 | p_0401 | AAAAAAATA | 3 | + | 0.75 | intergenic | AATATTCCAATATTCAATTAAGAATTTCTCACAAT**TGAAAT** |
| AE009951.2 | p_0402 | 1203399 | p_0402 | AGATATAAT | 4 | + | 0.83 | intergenic | TTTGTAACTCACATTTTTTTCTGGTTGGGGGAAGG**GGAATA** |
| AE009951.2 | p_0403 | 1207705 | p_0403 | ATTTTATAT | 3 | + | 0.65 | intergenic | AATTTTTATTTAATAGTATTTAATTTATTTATATA**AAACTC** |
| AE009951.2 | p_0404 | 1207685 | p_0404 | TTAATAATA | 3 | - | 1.0 | intergenic | TAAATTATATTAAATTAAATATCATTTAATTTTTT**AACTTG** |
| AE009951.2 | p_0405 | 1210531 | p_0405 | ATTTATAAT | 5 | + | 0.39 | intergenic | ATCCTAACTAATTATACCATAATATTTTTAATAAA**GTAACA** |
| AE009951.2 | p_0406 | 1210719 | p_0406 | TGTTAAAAT | 4 | + | 1.0 | intergenic | CTAAAAGAAAATAAAAAAATTAATATTTCTATAAG**TTCACA** |
| AE009951.2 | p_0407 | 1210463 | p_0407 | TGGTATAAT | 5 | - | 1.0 | intergenic | CATTATATTCATTATAAATATAAAAAATAATATTG**TTACTT** |
| AE009951.2 | p_0409 | 1220990 | p_0409 | AAAAAATAT | 3 | + | 1.0 | intergenic | ATAGTCGTTTTTTTTTTTTTTTTTTCAAGCACTGT**TTGAAA** |
| AE009951.2 | p_0410 | 1220916 | p_0410 | AAAAAAAAA | 3 | - | 1.0 | intergenic | TATTTTAGATATATTTTTTAAATATATTTTCAAAC**AGTGCT** |
| AE009951.2 | p_0411 | 1222524 | p_0411 | AAAAATATA | 3 | + | 1.0 | intergenic | CTTAAATTTGAAAAAAATATATAAAAATGATATAG**TTAAAC** |
| AE009951.2 | p_0412 | 1225429 | p_0412 | AAGATAATT | 3 | + | 0.96 | intergenic | TAATTAAGTGAAAAAATTAAGAGAGTGCAAATGAG**TATTTA** |
| AE009951.2 | p_0413 | 1225754 | p_0413 | AAGAAAAAA | 3 | + | 1.0 | intergenic | CTAAGTGCTTTTTATTTTAAAAAATAATTTAGAAC**AAAAAA** |
| AE009951.2 | p_0414 | 1234863 | p_0414 | GTTAAATGC | 2 | + | 0.99 | intergenic | AATATGGTAGTGAGTAAAATAAACGAACTGTGAAT**TGACGA** |
| AE009951.2 | p_0415 | 1245810 | p_0415 | TGCTATAAT | 5 | + | 1.0 | intergenic | ATAAAATAGTTTCATTATATATTAAAGTAATTAAA**TAAAAA** |
| AE009951.2 | p_0416 | 1245691 | p_0416 | ATATTTAAA | 3 | - | 0.9 | intergenic | AATATATAATGAAACTATTTTATATTGACAAACTA**TATAAA** |
| AE009951.2 | p_0417 | 1253867 | p_0417 | TTGTATAAT | 4 | + | 0.79 | intergenic | AATTGTAACTCACTTATTTTTTCTATATATATATT**AAAAAA** |
| AE009951.2 | p_0418 | 1253950 | p_0418 | TGTTAAAAT | 4 | + | 1.0 | intergenic | AATTTCTATCAATGCCATTTTTTAATAAAAAAAAG**TTGAGA** |
| AE009951.2 | p_0419 | 1256757 | p_0419 | TGCTATATT | 3 | + | 1.0 | intergenic | TTATATTATATTGCAGAAAATTAAATAAAACAATA**TTTATT** |
| AE009951.2 | p_0420 | 1256683 | p_0420 | CAATATAAT | 4 | - | 0.99 | intergenic | TTATAATATAGCATAAATTAAAATAAAAATAAATA**TTGTTT** |
| AE009951.2 | p_0421 | 1261689 | p_0421 | TGGTATAAT | 5 | + | 1.0 | intergenic | TTCATTTATAACAACTCTTTTTTTGTTAGAATGTT**TAATAT** |
| AE009951.2 | p_0422 | 1270989 | p_0422 | ATTAAAAAA | 3 | + | 1.0 | intergenic | GAAATAAAAAAATTTTTAATATCCCTTTTTAATTT**TAGAAA** |
| AE009951.2 | p_0423 | 1271579 | p_0423 | GTATATAAT | 4 | + | 1.0 | intergenic | CTTTTTTACTTAAAATATTTTCTTTAAATTAATAT**TTGATT** |

| | | | | | | | | | |
|---|---|---|---|---|---|---|---|---|---|
| AE009951.2 | p_0424 | 1279796 | p_0424 | AAAAAATAA | 3 | - | 1.0 | intergenic | AAAATGTAACAATTTTTTATTTAAAATATTTTTTC**TTGTGA** |
| AE009951.2 | p_0425 | 1281399 | p_0425 | TGATAAATT | 3 | + | 0.99 | intergenic | TATGATATTTTTAAATATATTATTATAAAAAAGTC**TTGTTA** |
| AE009951.2 | p_0426 | 1281263 | p_0426 | TTATATAAT | 5 | - | 0.97 | intergenic | ATCATAATATCAAGATTATATGAAAAATAAGTTTA**TTGACT** |
| AE009951.2 | p_0427 | 1286173 | p_0427 | TGGTATAAT | 5 | - | 0.99 | intergenic | TTAAAGTATTTAGTTAAAAGACTGTTGTAAATTAT**TTTTAT** |
| AE009951.2 | p_0428 | 1290972 | p_0428 | TAAAAAAAT | 3 | + | 1.0 | intergenic | AAAATGATGTCAAATTTTTTGCTTTTAACGTTAAA**TATAAT** |
| AE009951.2 | p_0429 | 1290878 | p_0429 | TGTTATTTT | 3 | - | 0.98 | intergenic | AAGAAAAATTATATTTAACGTTAAAAGCAAAAAAT**TTGACA** |
| AE009951.2 | p_0430 | 1295162 | p_0430 | TATAAAATA | 3 | + | 1.0 | intergenic | TCCTCTAATTAATATATTTTAACAGTATATTTTAG**CATAAT** |
| AE009951.2 | p_0431 | 1295209 | p_0431 | TTATATAAT | 5 | - | 0.94 | intergenic | GATTGAAGATAATTCTTTTGATAAATGTTTTAGTT**GTTAAT** |
| AE009951.2 | p_0432 | 1295112 | p_0432 | TGCTAAAAT | 3 | - | 1.0 | intergenic | TTATTGTAATTTCAATTATTAAGTAAAATATTTTA**TAGAAA** |
| AE009951.2 | p_0433 | 1309355 | p_0433 | TTTTATAAA | 3 | + | 1.0 | intergenic | ATATTATAAATGTTTATATTTTATTTTTTGTGGTT**TTATCG** |
| AE009951.2 | p_0434 | 1309354 | p_0434 | TGTTTTATA | 3 | - | 1.0 | intergenic | AAAATTATAACATAAAAATTACAAAATTTTATAAA**AATCAA** |
| AE009951.2 | p_0435 | 1310493 | p_0435 | AAGTAAAAA | 3 | + | 0.57 | intergenic | AATAAGAACAGTTAATTTTATTTTAGTTAATTTTT**GGTTTT** |
| AE009951.2 | p_0436 | 1314319 | p_0436 | TTTTATATA | 3 | + | 1.0 | intergenic | ATAAAATGAAATTTAGAGAATTTTTTTGAGAATAA**AATCTT** |
| AE009951.2 | p_0437 | 1317837 | p_0437 | TTATATAAT | 5 | + | 1.0 | intergenic | ATTCATATATATAAAAAATATGAATATAAAAGTTC**TTGATA** |
| AE009951.2 | p_0438 | 1317683 | p_0438 | TATTATATA | 3 | - | 1.0 | intergenic | TTCTATTAAAATTCTAAAAAATTTTTTTAAATAAT**TGTTGA** |
| AE009951.2 | p_0439 | 1319361 | p_0439 | TGATATAAT | 5 | + | 1.0 | intergenic | TAATTTACAATAGTTTTTTTTATTTAAAAATATTT**CTTGCA** |
| AE009951.2 | p_0440 | 1321152 | p_0440 | TTTTATATA | 3 | + | 1.0 | intergenic | TAAGATGAATAGAAATTAATATTTGTCTTATCTTT**TTTGTT** |
| AE009951.2 | p_0441 | 1323900 | p_0441 | TGCTAAAAT | 3 | + | 0.98 | intergenic | TAAAAGTTCTCACAGCTGTTACTAGATTTGAATTG**ATACCT** |
| AE009951.2 | p_0442 | 1324086 | p_0442 | TGGATATAT | 3 | + | 1.0 | intergenic | TTTATATAATAAAAAACTTATATATTAAAATAATT**ATATTG** |
| AE009951.2 | p_0443 | 1323962 | p_0443 | TTGTAAATT | 3 | - | 0.91 | intergenic | AGTTTTTTATTATATAAATAATTGATAGTAAAAGT**AATTTG** |
| AE009951.2 | p_0444 | 1323740 | p_0444 | TGATAAAAT | 4 | - | 1.0 | intergenic | ATTATTAAACATTAAGTTTAGCTAAATAAAGTTTA**TTGTAT** |
| AE009951.2 | p_0445 | 1327761 | p_0445 | TTTAAAAAA | 3 | + | 1.0 | intergenic | ATTAAAAACTTCTAAAAAAAATATTTTTTTAGTAT**TAATAC** |
| AE009951.2 | p_0446 | 1332608 | p_0446 | AAATATAAT | 5 | + | 1.0 | intergenic | AATAATATATTATTTTTCATTTGTCAATAGAAAAA**ACAACA** |
| AE009951.2 | p_0447 | 1332534 | p_0447 | TGAAAAATA | 4 | - | 1.0 | intergenic | AAAAATTATATTTTAAAAAATAAATATTGTTGTTT**TTTCTA** |
| AE009951.2 | p_0448 | 1333986 | p_0448 | TTTTAATAT | 3 | + | 0.98 | intergenic | TTATTATATTACTAAAATAAACATCAAAGAAAAAT**TATTTT** |
| AE009951.2 | p_0449 | 1338875 | p_0449 | TGATAAAAT | 4 | - | 1.0 | intergenic | CAAATGTTACAGAAATAAAAAAAATTTAAAGTTAA**TATTAA** |
| AE009951.2 | p_0450 | 1342693 | p_0450 | TGTTATAAT | 5 | + | 1.0 | intergenic | AAAAATTGCAAATTCTTTTAAATTTGTCCTTTTAT**GTCCTA** |
| AE009951.2 | p_0451 | 1342594 | p_0451 | TAACCTAAA | 2 | - | 0.98 | intergenic | AATAGGACATAAAAGGACAAATTTAAAAGAATTTG**CAATTT** |
| AE009951.2 | p_0452 | 1346544 | p_0452 | TGTTAAAAT | 4 | + | 1.0 | intergenic | TTAATCAAAATTTAATATTTTAATATATTATACAC**TTTTCT** |
| AE009951.2 | p_0453 | 1346492 | p_0453 | GTGTATAAT | 4 | - | 0.98 | intergenic | TTTTCTCCGATCTCAAATATTTTTAAATTTTAACA**TATTAT** |
| AE009951.2 | p_0454 | 1354089 | p_0454 | AAAATAATA | 3 | + | 1.0 | intergenic | AGTGGTTTTTCATTGCCAAAGGAATATTAAAAAAA**ATAAAA** |
| AE009951.2 | p_0455 | 1362048 | p_0455 | TAATATAAT | 5 | + | 1.0 | intergenic | CCTCCTTAAAAAATAAAAATATTTTTGCTATTATA**TATTAT** |
| AE009951.2 | p_0456 | 1362087 | p_0456 | TATATTTAT | 3 | - | 0.92 | intergenic | TTTATTATAATATATTTTATTTTTATTGAATAATC**TTATTA** |
| AE009951.2 | p_0457 | 1361999 | p_0457 | ATATATAAT | 5 | - | 1.0 | intergenic | TATATTTATATTAATTATAAAATTTATATATTATA**TTAGAT** |
| AE009951.2 | p_0458 | 1372197 | p_0458 | GCTTATAAT | 4 | + | 0.82 | intergenic | TTTAATTTTTATCTGTTAAGAAATCTGGCTAGTAA**TGAATT** |
| AE009951.2 | p_0459 | 1373224 | p_0459 | TAATAATAA | 3 | + | 0.37 | intergenic | TGGAGAAAAAAAGTGTAAATAATATTTCAAAAATT**GAAAAG** |
| AE009951.2 | p_0460 | 1378530 | p_0460 | TAAAATAAA | 3 | + | 1.0 | intergenic | TATTTGTATTAATAATAATATTATTATCTAAGTGT**AACATT** |
| AE009951.2 | p_0461 | 1378482 | p_0461 | AATAAATGT | 2 | - | 0.99 | intergenic | TATTTATTTCATTATAATGTTATTAAAAATTTTAT**TTTACT** |
| AE009951.2 | p_0462 | 1386449 | p_0462 | AATGTTATA | 2 | + | 1.0 | intergenic | TTAACTATTTTTAAAATATTATACCATTTTTATAT**AGAAAA** |
| AE009951.2 | p_0463 | 1386388 | p_0463 | TGGTATAAT | 5 | - | 1.0 | intergenic | TATTCTACTATATTTACTATAACATTTCAAATATT**TTATTT** |
| AE009951.2 | p_0464 | 1389678 | p_0464 | TTATATAAT | 5 | + | 0.46 | intergenic | TTATAATAGTTAATCTTTTTTAGATATTGATATTT**TATATA** |
| AE009951.2 | p_0465 | 1392800 | p_0465 | TACAATATA | 3 | + | 1.0 | intergenic | TTTCATATATAAAATTTTAAATGTTTATATCTCTT**GCATTT** |
| AE009951.2 | p_0466 | 1395932 | p_0466 | TGTTAAAAT | 4 | + | 1.0 | intergenic | TTAATATTATTTTAACATAGGAATAAAATTTTCTC**TATAAG** |
| AE009951.2 | p_0467 | 1395862 | p_0467 | TGTTAAAAT | 4 | - | 1.0 | intergenic | ATTCTTAAATTTTAACATAAAAATATACAAACTTA**TAGAGA** |
| AE009951.2 | p_0468 | 1400127 | p_0468 | TAATATAAT | 5 | + | 0.88 | intergenic | TTAGAAATTTTATATATTTTATATTTTTAAAGTCA**TTTTAA** |
| AE009951.2 | p_0469 | 1400219 | p_0469 | TGTAAAATA | 4 | + | 1.0 | intergenic | TAAATATTTTTATTTTATAGAAATTTTTAAAAAAT**TGCTAT** |
| AE009951.2 | p_0470 | 1400151 | p_0470 | TATAAAATA | 3 | - | 1.0 | intergenic | TCAATCAATATATTTTACACTATTAAGGAAAAAAT**AGCAAT** |
| AE009951.2 | p_0471 | 1407712 | p_0471 | TTATATAAT | 5 | - | 0.62 | intergenic | ATTACAAACCCATTTTGTAATAGCCTTTTCTATTA**TTGATA** |
| AE009951.2 | p_0472 | 1418979 | p_0472 | AGCTATAAT | 4 | + | 0.99 | intergenic | TTTTTAAATAGAAAAATTTACATAGCTCTATAATA**TTGACA** |
| AE009951.2 | p_0473 | 1418801 | p_0473 | TGATATTAT | 3 | - | 1.0 | intergenic | TTATTAAATTAATTAATCTACTTTTTAAAATTTCA**TTGAAA** |
| AE009951.2 | p_0474 | 1420273 | p_0474 | TGTTAAAAT | 4 | + | 1.0 | intergenic | TTGCTATTTTGATAGAAAATTATAACATTAAATTA**TAGAAA** |
| AE009951.2 | p_0475 | 1420226 | p_0475 | TTCTATAAT | 4 | - | 1.0 | intergenic | TTAATTTTTTAATTTTTAGAAGATCTTTATAATTT**TAACAT** |
| AE009951.2 | p_0476 | 1422651 | p_0476 | ATATATAAT | 5 | + | 1.0 | intergenic | AAATTTAAAATATAAAATTTATACAAAATTTGTTA**TTGATT** |
| AE009951.2 | p_0477 | 1430678 | p_0477 | TTTAAAATT | 3 | + | 1.0 | intergenic | TTATTATATCCCATTATTTTTATGTTATGTAAAAT**AAATAA** |
| AE009951.2 | p_0478 | 1439190 | p_0478 | TACAATAAA | 3 | + | 0.99 | intergenic | TTTGATTTTATTATTATATCATAATTAGTAAAAAT**AAAATA** |
| AE009951.2 | p_0479 | 1439273 | p_0479 | ATAGAATAA | 3 | + | 1.0 | intergenic | AAGAAAATTTAAAGAGAAAAATACATAAAAGGCTT**GACATA** |
| AE009951.2 | p_0480 | 1439124 | p_0480 | TGATATAAT | 5 | - | 1.0 | intergenic | ACATTTTGTAGATTTATTGTACTAAATTAATAAAG**TATTTT** |
| AE009951.2 | p_0481 | 1450510 | p_0481 | TTTATATAT | 3 | + | 1.0 | intergenic | AGAGAATTTTTTTGAGAACTAAAAACTCAAAAAAG**TTCTCT** |
| AE009951.2 | p_0482 | 1450611 | p_0482 | TAAATTATA | 3 | + | 1.0 | intergenic | AAAAACTAAATAAGAAAAGAATATTTTTTTTTAAA**AAGAAA** |
| AE009951.2 | p_0483 | 1452510 | p_0483 | TGTTAAAAT | 4 | + | 1.0 | intergenic | TATTAAATAGATGTACATTAATTAGTAAAAATAAT**TTGTTC** |
| AE009951.2 | p_0484 | 1452358 | p_0484 | AGTTATAAT | 4 | - | 0.99 | intergenic | TTTTCTATAATTAAATTATTTCAAAACAAATTTTG**TAGATA** |
| AE009951.2 | p_0485 | 1459943 | p_0485 | ATTAATATG | 3 | - | 1.0 | intergenic | AAATAATTTTGCAATGGCTTTTTTTATAAAATTTT**GTTACT** |

| | | | | | | | | | |
|---|---|---|---|---|---|---|---|---|---|
| AE009951.2 | p_0486 | 1462232 | p_0486 | ATTTTATAT | 3 | - | 0.45 | intergenic | TTATTTTATGGATAAGGAAATTAAAAATACATTTT**GTATTT** |
| AE009951.2 | p_0487 | 1464726 | p_0487 | TTATATAAT | 5 | - | 1.0 | intergenic | ACTTCAGGTATTTTTATTTTATTAAACTTTTTTAG**TATACA** |
| AE009951.2 | p_0488 | 1472861 | p_0488 | TGTTATAAT | 5 | + | 1.0 | intergenic | TTGTATTTAAAACATTATTATAAAAAATAAAATTC**TTGAAT** |
| AE009951.2 | p_0489 | 1472740 | p_0489 | TGATATAAT | 5 | - | 1.0 | intergenic | ATAATAATGTTTTAAATACAAACACTTAGTAAAAA**TAGTTT** |
| AE009951.2 | p_0490 | 1476592 | p_0490 | AATTAAAAA | 3 | + | 0.49 | intergenic | TTAATTTGATAATTTTATCATTTTGAGTTATGTCA**ATTTCT** |
| AE009951.2 | p_0491 | 1477362 | p_0491 | AATTTATAA | 3 | + | 0.97 | intergenic | CTATTCTACATTACAATTTAAACTATATCTTGAAT**TTTTTA** |
| AE009951.2 | p_0492 | 1477299 | p_0492 | AGTTTAAAT | 3 | - | 0.53 | intergenic | AAAATTTACTTAATATTATAAATTTTACTTTATCT**AAATAA** |
| AE009951.2 | p_0493 | 1477180 | p_0493 | TGTAATATA | 4 | - | 1.0 | intergenic | TATAATAAAATTTCTAATTATAAAAAATTTTTTAC**CTAAAA** |
| AE009951.2 | p_0494 | 1479465 | p_0494 | TATTAAAAT | 3 | + | 1.0 | intergenic | TTAAATAAAATTTTATATTTTTTATAAAATATTTC**TTGACT** |
| AE009951.2 | p_0495 | 1481937 | p_0495 | TGCTATAAT | 5 | + | 1.0 | intergenic | TTTTAAAAATTTGAGGTTATATTGTAATTTAATTC**ACTATA** |
| AE009951.2 | p_0496 | 1490093 | p_0496 | TGTTAAAAT | 4 | + | 1.0 | intergenic | TCAAGAGAAAAAATACAAATTTTTAGTTTTAATAT**TAAACA** |
| AE009951.2 | p_0497 | 1490198 | p_0497 | AATTATAAT | 5 | + | 0.99 | intergenic | TAACTATTGGAGCCTCAATTTTAATAAACTTTAAG**TTGCAA** |
| AE009951.2 | p_0498 | 1490360 | p_0498 | AAAGAATAA | 3 | + | 1.0 | intergenic | GGTTTGAGTGTCGCTTTTAATAAAACAGAAAAGAA**AGTAAT** |
| AE009951.2 | p_0499 | 1490137 | p_0499 | TATTAAAAT | 3 | - | 0.84 | intergenic | TCAATATTTTTTATTTCATTATAATTTATTTCATT**ATCTTT** |
| AE009951.2 | p_0500 | 1489994 | p_0500 | AAATTATTG | 2 | - | 1.0 | intergenic | TTTGTTTAATATTAAAACTAAAAATTTGTATTTTT**TCTCTT** |
| AE009951.2 | p_0501 | 1494339 | p_0501 | TATTATAAT | 5 | + | 1.0 | intergenic | ATAAAATATGTTATTTAAAAAATCAAATATTTTTT**AAAGAG** |
| AE009951.2 | p_0502 | 1494258 | p_0502 | TTTTATAAT | 5 | - | 1.0 | intergenic | ATAATATCAAAATTAATTTTCTCTTTAAAAAATAT**TTGATT** |
| AE009951.2 | p_0503 | 1500418 | p_0503 | TATTTTTAA | 3 | + | 1.0 | intergenic | AAAAACTATAAAAATAATATTTATTTATAACTACT**ATATAA** |
| AE009951.2 | p_0504 | 1500500 | p_0504 | TTTTAAATA | 3 | + | 0.01 | intergenic | ATTATAAATTAACTATATGATTTTATAATTTAATG**TTATAA** |
| AE009951.2 | p_0505 | 1501430 | p_0505 | AATTATAAA | 3 | + | 1.0 | intergenic | AGTAATTTTTATTTTGAATGATTTTATAAAAAATT**AAATAT** |
| AE009951.2 | p_0506 | 1503631 | p_0506 | TTAAAAATT | 3 | + | 1.0 | intergenic | TTGTTTTATAAATTAAAAATTATATTATAAATAAC**TTATAT** |
| AE009951.2 | p_0507 | 1503514 | p_0507 | CAGTATATA | 3 | - | 1.0 | intergenic | ATATAATTTTTAATTTATAAAACAAATTTGTATTT**ATTTGA** |
| AE009951.2 | p_0508 | 1504004 | p_0508 | AAAAATTAG | 2 | + | 1.0 | intergenic | AGCATTGGAAATACTAGATTAAAATTAAAAAATAT**TCAAAA** |
| AE009951.2 | p_0509 | 1505372 | p_0509 | TGATATAAT | 5 | + | 0.67 | intergenic | AGAGGGAGTTATTGTAAATAAATTAAAGACAATAA**TTCCCT** |
| AE009951.2 | p_0510 | 1520641 | p_0510 | TGATATTAT | 3 | - | 1.0 | intergenic | ATTTTATATTTTTTCTCAATTTCTACTATTCTATA**TAGCAA** |
| AE009951.2 | p_0511 | 1521678 | p_0511 | TAGTAAATT | 3 | - | 0.82 | intergenic | GAAAGAAGTAAAGATCCTAATATTGTTTATTATAA**ATATGC** |
| AE009951.2 | p_0512 | 1523032 | p_0512 | TTTTAAAAT | 3 | - | 1.0 | intergenic | AAGGATTGACTTTTGCAAGAAACAAGAGTTAATCC**TTTTTT** |
| AE009951.2 | p_0513 | 1531893 | p_0513 | TGATATAAT | 5 | - | 0.99 | intergenic | AAAATACAAACAGAGGTTAAAATTATAGCCTCTGT**TTTTTT** |
| AE009951.2 | p_0514 | 1535774 | p_0514 | TGTTATAAT | 5 | - | 1.0 | intergenic | TTTAGAAAAAAATTATCCATTTTTATTTTGAAAAAT**TACCAA** |
| AE009951.2 | p_0515 | 1539015 | p_0515 | ATAATATAT | 3 | - | 0.95 | intergenic | GTGACAGTCTTATTTTTTTATCTAATAAAAAATATT**GACAAT** |
| AE009951.2 | p_0516 | 1540013 | p_0516 | TGACAAAAT | 3 | + | 0.99 | intergenic | AGATTTTTTTTGGCTATCTTATTTATTATTTTTTA**GTTTTA** |
| AE009951.2 | p_0517 | 1539922 | p_0517 | TGCTATAAT | 5 | - | 1.0 | intergenic | AATTACTTGTTAAAACTAAAAAATAATAAATAAGA**TAGCCA** |
| AE009951.2 | p_0518 | 1542878 | p_0518 | TTTTATAAA | 3 | - | 1.0 | intergenic | AGTGTTTTATTTTAAAATAAAAAATATAAAAAAAAA**AACACT** |
| AE009951.2 | p_0519 | 1545788 | p_0519 | TTATATAAT | 5 | + | 0.98 | intergenic | AATTTGTATTTTTTTATTTTTATAAGCTAATAATT**GTTAGA** |
| AE009951.2 | p_0520 | 1553170 | p_0520 | TGTTAAAAT | 4 | + | 1.0 | intergenic | AAATATATTTTTAAATAAAGTCAACTAAAAAAAGC**TACATT** |
| AE009951.2 | p_0521 | 1553092 | p_0521 | TTTAAAAAT | 3 | - | 1.0 | intergenic | ATTTTAACATATTTTTTGAAAAAAATGTAGCTTTT**TTTAGT** |
| AE009951.2 | p_0522 | 1555961 | p_0522 | TGTTATAAT | 5 | + | 1.0 | intergenic | ATTATATGAATATTTTATCAAATTTTTATAAAATA**GCAAGA** |
| AE009951.2 | p_0523 | 1555894 | p_0523 | TGATAAAAT | 4 | - | 1.0 | intergenic | CCTCCATATAAATTATAACATTTTATACTTAGAAT**CTTGCT** |
| AE009951.2 | p_0524 | 1565278 | p_0524 | TCTTAAAAT | 3 | - | 0.18 | intergenic | ACTCGTTTCACTCAAACACAGCAAGATTTGCTTGG**CTCATT** |
| AE009951.2 | p_0525 | 1572676 | p_0525 | TTTTAATTT | 3 | + | 1.0 | intergenic | ATTTTCACCTTCCTAAAATAAATTTATATATTTTA**ATTATA** |
| AE009951.2 | p_0526 | 1572633 | p_0526 | TTGTATAAT | 4 | - | 1.0 | intergenic | TTTTTCCTCCATAAAAATTATTTTCTATTCAATTC**TATCAT** |
| AE009951.2 | p_0527 | 1579157 | p_0527 | TAAAAAAAT | 3 | + | 1.0 | intergenic | TTTTATATTTGGTATATAATAGCATTTTTTTTAAA**TTTTTC** |
| AE009951.2 | p_0528 | 1579099 | p_0528 | TTTAAAAAA | 3 | - | 1.0 | intergenic | TAGATTTTTATCTATAATTTTAAAATATTTTTTTA**ATTTTT** |
| AE009951.2 | p_0529 | 1584744 | p_0529 | AATTTATAA | 3 | + | 1.0 | intergenic | ATAATAGCACAAATATAAAATAAAAAATAGTATTT**TCTATT** |
| AE009951.2 | p_0530 | 1584663 | p_0530 | TATTATAAA | 3 | - | 1.0 | intergenic | TAAATTTTACTGTCAAGTTTAATAGAAAATACTAT**TTTTTA** |
| AE009951.2 | p_0531 | 1595504 | p_0531 | ATTAAAAAT | 3 | + | 1.0 | intergenic | TATACTTTTATTTCTTTTTTTTCTCTTTTTTCTCT**CTGTAA** |
| AE009951.2 | p_0532 | 1595394 | p_0532 | AACTATAAT | 4 | - | 0.61 | intergenic | GAAAAAAGAGAAAAAAAAGAAATAAAAGTATAATG**TTTATT** |
| AE009951.2 | p_0533 | 1598976 | p_0533 | TGTTAAAAT | 4 | - | 1.0 | intergenic | TCTTTTTTTAGTATTCATAATGTTTAGTAATTAAT**TGACAA** |
| AE009951.2 | p_0534 | 1600555 | p_0534 | TGCTATAAT | 5 | + | 0.74 | intergenic | CATATCGTCTATTTTTTTATTTAAATTATTTTAAAT**TTGACA** |
| AE009951.2 | p_0535 | 1611929 | p_0535 | TGAAATAAT | 4 | - | 1.0 | intergenic | AGGACAATATTATAAAAAATAAAATTGAAAAAAAC**TACAAA** |
| AE009951.2 | p_0537 | 1630682 | p_0537 | TTATATAAT | 5 | - | 1.0 | intergenic | GAATAAAAAATTTTATTTGAAAATTAAAATAAAAAG**TTGATT** |
| AE009951.2 | p_0538 | 1632161 | p_0538 | TGATATAAT | 5 | + | 1.0 | intergenic | AAAATCTTAATTTTATTATACTTTAAATTTTATCT**TTTTAA** |
| AE009951.2 | p_0539 | 1632097 | p_0539 | AAGTATAAT | 4 | - | 1.0 | intergenic | TTTTACTATAAATAATTATATCACAAGTAAAATAC**TTTTAA** |
| AE009951.2 | p_0540 | 1641083 | p_0540 | TAAAATAAA | 3 | - | 1.0 | intergenic | ATTTTTTACTTAAAATCTACAATTGTAACTCATTT**ATTTTC** |
| AE009951.2 | p_0541 | 1645916 | p_0541 | TGCTATAAT | 5 | + | 0.99 | intergenic | TTTTGTAAATAGCTAAAGAAATAAAATTATTGAAA**TAGAAA** |
| AE009951.2 | p_0542 | 1645817 | p_0542 | TGTATTATA | 3 | - | 0.99 | intergenic | AATTTCTATTTCAATAATTTTATTTCTTTAGCTAT**TTACAA** |
| AE009951.2 | p_0543 | 1645620 | p_0543 | TTAAATAAA | 3 | - | 0.07 | intergenic | TATGCTTAACAAAAAGGAGGAAACAGTGAAATTAA**TAACTA** |
| AE009951.2 | p_0544 | 1652320 | p_0544 | TTTTATATA | 3 | + | 0.96 | intergenic | AGTATTTTTCTAAAATATTAATTTTTTAAATAAAT**GAAAGT** |
| AE009951.2 | p_0545 | 1652257 | p_0545 | TTTAAAAAA | 3 | - | 1.0 | intergenic | TGAAAAGAATAAAAATATATAAAATTTATATTAAA**ATGACT** |
| AE009951.2 | p_0546 | 1655010 | p_0546 | TTCAAATAT | 3 | + | 0.94 | intergenic | TTTAAAAATAGAAAACACTTAGAAAAATTTCTAGG**TGTTTT** |
| AE009951.2 | p_0547 | 1660462 | p_0547 | TGTTATAAT | 5 | + | 1.0 | intergenic | TAAAGATAAAGTTACTTTTTGATATGTTTTTTTAT**TTGTTT** |
| AE009951.2 | p_0548 | 1661603 | p_0548 | TGGTAAAAA | 3 | + | 1.0 | intergenic | ATTTAAATTTTTTATCTATTATTTTTTATTATTTT**ATAATA** |

| | | | | | | | | | |
|---|---|---|---|---|---|---|---|---|---|
| AE009951.2 | p_0549 | 1661563 | p_0549 | TTTTTATAT | 3 | - | 1.0 | intergenic | AAAAAATATTTTAAATTTATTAATTTAATTATAAC**AAATTT** |
| AE009951.2 | p_0553 | 1664296 | p_0553 | TTATATAAT | 5 | - | 0.97 | intergenic | TTGGACTTCTTTTTTTATAAAAGTTAAAATTATAA**GGAGTT** |
| AE009951.2 | p_0554 | 1670487 | p_0554 | TGTTATAAT | 5 | + | 1.0 | intergenic | AACTAAAATATTATATCATATATAAAAAATTTATA**ATAGCA** |
| AE009951.2 | p_0555 | 1670418 | p_0555 | TGATATAAT | 5 | - | 0.99 | intergenic | TTTCAATTTATTATAACATATTTTACTTGACTTGC**TATTAT** |
| AE009951.2 | p_0556 | 1674708 | p_0556 | TGATATTAT | 3 | - | 1.0 | intergenic | AACATAAGAAAAATATAAAATCAATAGGCTTTTTA**TTGATT** |
| AE009951.2 | p_0557 | 1678153 | p_0557 | TTTTTAAAA | 3 | - | 1.0 | intergenic | TTATCACTAGCTCTAAATCCTTTTTTATTAAAAAA**TAAAAG** |
| AE009951.2 | p_0559 | 1687729 | p_0559 | AGTTATAAT | 4 | + | 1.0 | intergenic | GATTATTTATTATATCATAATTTTTATAAAAAATA**TTAAGA** |
| AE009951.2 | p_0560 | 1687666 | p_0560 | TTTATAAAA | 3 | - | 1.0 | intergenic | TTTTAAAAAGATTTTATTATAACTGCTTCTTTTAA**TTTTCT** |
| AE009951.2 | p_0561 | 1696214 | p_0561 | TGCTATAAT | 5 | + | 0.92 | intergenic | AATTTAGCATACTTATGAAGTTGCTCTTTTTATAA**TAGAAA** |
| AE009951.2 | p_0562 | 1696088 | p_0562 | TGTTAAAAT | 4 | - | 1.0 | intergenic | ATAAGTATGCTAAATTAAAAAGCAATCTATTTAAG**TTGCTT** |
| AE009951.2 | p_0563 | 1703743 | p_0563 | TATTATAAT | 5 | + | 1.0 | intergenic | TATTTTCTTATTTTTTGATAAGAATTATAAAAAAT**AGTAGA** |
| AE009951.2 | p_0564 | 1703748 | p_0564 | ATTTAATAT | 3 | - | 1.0 | intergenic | CTTTTTTCTAAAACTTACATTTTTTTCTATATTCT**AAATGT** |
| AE009951.2 | p_0565 | 1703626 | p_0565 | AGTATTATA | 3 | - | 0.96 | intergenic | ATTCTTATCAAAAAATAAGAAAATATTATTTTTGC**TATTGA** |
| AE009951.2 | p_0566 | 1709408 | p_0566 | TATAAAAAA | 3 | + | 1.0 | intergenic | TAGATAAAAAACAAATGTTTTTTTATTTTTTAATG**TACAAA** |
| AE009951.2 | p_0567 | 1709314 | p_0567 | TTGTATAAT | 4 | - | 1.0 | intergenic | ATAAAATTTTGTACATTAAAAAATAAAAAAACATT**TGTTTT** |
| AE009951.2 | p_0568 | 1711968 | p_0568 | TGCTATAAT | 5 | + | 1.0 | intergenic | TCATTTATTAAAGTGGGAGTTTTTATTATTTTTTA**TTATAA** |
| AE009951.2 | p_0569 | 1722157 | p_0569 | TGATATAAT | 5 | + | 1.0 | intergenic | TTGAGAGAAACTTAAAAAAATTCTTTTTTCATATC**ACTTAA** |
| AE009951.2 | p_0570 | 1724237 | p_0570 | TTCTAAATT | 3 | + | 1.0 | intergenic | ATAAATACCCAAAAAAGAAAAATTTTCAAAAAAAA**TATAAA** |
| AE009951.2 | p_0571 | 1724159 | p_0571 | GGTATTTAT | 2 | - | 1.0 | intergenic | AATTTAGAATAAATAAAGGTTTTTTTATATTTTTT**TTGAAA** |
| AE009951.2 | p_0572 | 1724917 | p_0572 | TTATTTATA | 3 | + | 1.0 | intergenic | AATATAATTAAATTTTAAAGGTAAGGAAAAATTCT**TTACCT** |
| AE009951.2 | p_0573 | 1725646 | p_0573 | TGGTATAAT | 5 | + | 1.0 | intergenic | AAGATTTCTAAATGAAATTTTAAATCCTTTTTATT**TGATAA** |
| AE009951.2 | p_0574 | 1727799 | p_0574 | TATTATAAA | 3 | + | 1.0 | intergenic | AAGAAACAATTAAAATTAAAATAAATATATTTAAA**TCTATA** |
| AE009951.2 | p_0575 | 1727677 | p_0575 | TGTATAAAA | 4 | - | 0.99 | intergenic | TTTAATTTTAATTGTTTCTTAATTATTAAAATAAA**ATTATA** |
| AE009951.2 | p_0576 | 1729729 | p_0576 | TGTTATAAT | 5 | + | 1.0 | intergenic | AATATGGATGCTTTTTTATAAAATAAATATAATTC**TTTTAA** |
| AE009951.2 | p_0577 | 1731777 | p_0577 | TAGTATAAT | 4 | + | 0.99 | intergenic | AAGGATTCAATCTTTTATTATAGAAATCTGCAACC**TTTACA** |
| AE009951.2 | p_0578 | 1739965 | p_0578 | AATTAAAAT | 3 | + | 1.0 | intergenic | CTTTTATTCCCTTTAAAAATATTTTTTATATTATA**TCATAG** |
| AE009951.2 | p_0579 | 1740254 | p_0579 | TGAATATAT | 4 | - | 1.0 | intergenic | TTGCTATTTTCAAACTTAATTACGCTCATTTCTCC**CCTCTC** |
| AE009951.2 | p_0580 | 1744076 | p_0580 | AAAGTATTA | 2 | + | 1.0 | intergenic | AAAGAACAGTATTAGGTATTAAAGGAAAAAATATA**TAAATT** |
| AE009951.2 | p_0581 | 1744672 | p_0581 | GAGTATAAT | 4 | + | 0.96 | intergenic | TAGAGTAAAAAAAATTTAAAAAACAACTCTAATAT**TATGTT** |
| AE009951.2 | p_0582 | 1758152 | p_0582 | TGAAATTAT | 3 | + | 1.0 | intergenic | TTTAGATAAATTAAAGAAAAAATATTATTATTTTA**AATCAA** |
| AE009951.2 | p_0583 | 1758069 | p_0583 | TAAAATATA | 3 | - | 1.0 | intergenic | TTCAAATATTTTTTCTTTTTGATTTAAAATAATAA**TATTTT** |
| AE009951.2 | p_0584 | 1761407 | p_0584 | ATATATAAT | 5 | + | 1.0 | intergenic | TTAGGATTTTTTTTATTTATAAATTTTAAATAAAT**TCATAA** |
| AE009951.2 | p_0585 | 1764792 | p_0585 | TATTATATA | 3 | - | 1.0 | intergenic | TTAAACTCAATAGCTTTTTTATTAATAAGAAATAT**AAAAAA** |
| AE009951.2 | p_0586 | 1767511 | p_0586 | TTTTAAAAA | 3 | + | 1.0 | intergenic | GTTTTTTTTAAAATCTAATTTATTTTAATTATATA**GTAAAA** |
| AE009951.2 | p_0587 | 1767421 | p_0587 | TGTTATAAT | 5 | - | 1.0 | intergenic | TTTGTCAATTTTTTTACTATATAATTAAAATAAAT**TAGATT** |
| AE009951.2 | p_0588 | 1774309 | p_0588 | TATAAAATA | 3 | + | 1.0 | intergenic | CTCAATAAAATTTAATCATTACATTTGAATATTAA**CATTTT** |
| AE009951.2 | p_0589 | 1774259 | p_0589 | TGTTAATAT | 4 | - | 1.0 | intergenic | ATCTAAAAATAAAATGCTATTAATTATTTATTTTA**TAGATT** |
| AE009951.2 | p_0590 | 1781612 | p_0590 | AGATAAAAT | 3 | + | 1.0 | intergenic | TATAGTAAAATTTAAAATAAATCAATAATTATTAT**TGACAA** |
| AE009951.2 | p_0591 | 1781532 | p_0591 | TACTATAAA | 3 | - | 0.81 | intergenic | TTTATCTGTTCTATTGTAAATTTGTCAATAATAAT**TATTGA** |
| AE009951.2 | p_0592 | 1782210 | p_0592 | ATAATATAT | 3 | + | 1.0 | intergenic | TTATAAAAATGTACAATTTAGAAACAAAATTATTT**GACATT** |
| AE009951.2 | p_0593 | 1785584 | p_0593 | AATTTTATA | 3 | + | 1.0 | intergenic | AAAATTGATATTAATACAAAAAAATTATATTTTGA**CTTGAA** |
| AE009951.2 | p_0594 | 1786235 | p_0594 | TTCTATAAT | 4 | + | 1.0 | intergenic | TTAAAATTTAAAATAAGAAATTTTACTAAATTTTT**ACCAAA** |
| AE009951.2 | p_0595 | 1791483 | p_0595 | TGTAGGATA | 2 | + | 1.0 | intergenic | TAAAAGCATTTTAAAAATTTTAAATAAATTTTACT**TGCTTT** |
| AE009951.2 | p_0596 | 1791663 | p_0596 | TTGATAATA | 3 | + | 0.56 | intergenic | GTAGCTTGTTAAAAAAACTTTAATAAAATTTATTT**ATTTTT** |
| AE009951.2 | p_0597 | 1791392 | p_0597 | TTCAAAAAA | 3 | - | 1.0 | intergenic | AAATCAAAAAAAAAGCAAGTAAAATTTATTTAAAAT**TTTTAA** |
| AE009951.2 | p_0598 | 1796050 | p_0598 | ACTTAATAT | 3 | - | 0.76 | intergenic | AGTATCCACTTTATTTTTTTATATTGTTTTTGCAA**TTAATT** |
| AE009951.2 | p_0599 | 1798113 | p_0599 | GTATATAAT | 4 | + | 0.98 | intergenic | AAATATGTTCATAATAATATTTACGATTTTTTATG**TTGCTT** |
| AE009951.2 | p_0600 | 1797980 | p_0600 | TATAAAATT | 3 | - | 0.69 | intergenic | AACATATTTTTTTTGAAGAGGGTGTACTTTTTTAA**AAATTT** |
| AE009951.2 | p_0601 | 1806722 | p_0601 | TTATATAAT | 5 | + | 1.0 | intergenic | AAAAGAAATAAAATTAAAAATCAAATTATTAGTTG**TGTACA** |
| AE009951.2 | p_0602 | 1806619 | p_0602 | TAGTATAAT | 4 | - | 0.42 | intergenic | TACACAACTAATAATTTGATTTTTAATTTTATTTC**TTTTAT** |
| AE009951.2 | p_0603 | 1820268 | p_0603 | TGTTATAAT | 5 | + | 0.96 | intergenic | TAAAATTTTCTTGCTAAAAATATAAATTTCAAAGG**TAGACT** |
| AE009951.2 | p_0604 | 1828658 | p_0604 | TGTTATAAT | 5 | - | 1.0 | intergenic | TTTTTTCTTATTTATAATTATTTAAAAGAGATTAT**TGCTTT** |
| AE009951.2 | p_0605 | 1847538 | p_0605 | TACTATTTT | 3 | - | 0.55 | intergenic | ATAGTATATTTTATTATGATTAAAATAAAAAATCA**AATAAA** |
| AE009951.2 | p_0606 | 1847421 | p_0606 | TGATAATAT | 4 | - | 1.0 | intergenic | TAAATTTCCTTACTTCTTATAATTAATATATAGGC**TTGAAA** |
| AE009951.2 | p_0607 | 1848504 | p_0607 | TTTTATAAT | 5 | + | 1.0 | intergenic | TTTATATTTTTATTTTATCAAATTTATATATATTT**TTATAG** |
| AE009951.2 | p_0608 | 1848640 | p_0608 | TATAAAATT | 3 | + | 0.83 | intergenic | TAAGGAGGATTAAAATGTTTCTTAGTAAAACAAGG**ATAAGG** |
| AE009951.2 | p_0609 | 1848728 | p_0609 | TTTTAATAA | 3 | + | 0.07 | intergenic | AAGGACTGTTTATGCCTTTGTTTTTTAGTATATTT**AGTATA** |
| AE009951.2 | p_0610 | 1848474 | p_0610 | TTATAAAAT | 3 | - | 1.0 | intergenic | TACTATAAAACAAATGTCAAATAAAATTTTATAAA**GAAATT** |
| AE009951.2 | p_0611 | 1850846 | p_0611 | AGATATAAT | 4 | + | 1.0 | intergenic | TAATAAGAATATTTTTCTATACATAAAAAAAATAC**TTGCAT** |
| AE009951.2 | p_0612 | 1855171 | p_0612 | TGATATAAT | 5 | + | 1.0 | intergenic | ATACTTTTTTTAGTAATTTAGAAAAGCCTTTCATC**TTTTTG** |
| AE009951.2 | p_0613 | 1856360 | p_0613 | TGTTATTAT | 3 | - | 0.8 | intergenic | TATCCTAACAAAAAGATTGATTTAAAACTTGTAAT**AGATGA** |
| AE009951.2 | p_0614 | 1859102 | p_0614 | ATAAAAAAG | 3 | + | 1.0 | intergenic | TCTTCTTTCTCCTTTCTAAAATTGATTTATTTTAT**TATACA** |

| | | | | | | | | | |
|---|---|---|---|---|---|---|---|---|---|
| AE009951.2 | p_0615 | 1859073 | p_0615 | CTTTTTTAT | 3 | - | 1.0 | intergenic | AATTTAAAGTTAAAACTAAAAAGAATGAAATTGAG**GTTTTA** |
| AE009951.2 | p_0616 | 1861145 | p_0616 | TGCTATAAT | 5 | - | 1.0 | intergenic | CATAACTACTCCTTTTTTATTGCAAGAAAATAAAA**CTTTTA** |
| AE009951.2 | p_0617 | 1878617 | p_0617 | TATAAAAAA | 3 | + | 1.0 | intergenic | TAAATTACAAAAAATTAAAATCTTTAATTTTTTTA**TAAACT** |
| AE009951.2 | p_0618 | 1878539 | p_0618 | TGTAATTTA | 3 | - | 1.0 | intergenic | TTTTTTATAGTGAATTTTTTAAAAGTTTATAAAAA**AATTAA** |
| AE009951.2 | p_0619 | 1879295 | p_0619 | TAATTTTAT | 3 | + | 1.0 | intergenic | ATAATACCTTCTCTTCTTCTTAGAAAAAAAATTTT**AGTATT** |
| AE009951.2 | p_0620 | 1879401 | p_0620 | TTAGTTATA | 2 | + | 1.0 | intergenic | TGGTAAAAAAAAGAACTCTGTATTTTTTTTAATGA**GTTCTT** |
| AE009951.2 | p_0621 | 1880456 | p_0621 | AACTATATA | 3 | + | 0.99 | intergenic | CTTCTTTTATTTTCATTTGAATTAAGATATTATAT**ATTGAC** |
| AE009951.2 | p_0622 | 1885801 | p_0622 | AAGAAAATT | 3 | - | 0.99 | intergenic | AAACTACCTTTCTATGATAGTGGTAATGGAGATTA**TATTGC** |
| AE009951.2 | p_0623 | 1891033 | p_0623 | ATGTTATAA | 3 | + | 1.0 | intergenic | ATTTTTAGAAAAAGAAAAGATATAAAAGTAATTAT**CTTTTA** |
| AE009951.2 | p_0624 | 1895209 | p_0624 | TACTATTAT | 3 | - | 0.95 | intergenic | TTTATATTTTATTTGAAGTAAGTTTTTATTTTGAT**AAAATA** |
| AE009951.2 | p_0625 | 1904457 | p_0625 | TGATATAAT | 5 | - | 1.0 | intergenic | AAGAAAAAACTACTTGCTCTTTTTTTTAATTAAATA**ATGCAA** |
| AE009951.2 | p_0626 | 1908079 | p_0626 | TTGATATAT | 3 | + | 1.0 | intergenic | AAAATATTAAAAAATAAATATTTTGTATAGGAAAT**ATATTG** |
| AE009951.2 | p_0627 | 1907961 | p_0627 | TGTAATATA | 4 | - | 1.0 | intergenic | AAAATATTTATTTTTTAATATTTTAAAAAAAAAGA**GATAGA** |
| AE009951.2 | p_0628 | 1915183 | p_0628 | AATTATAAA | 3 | + | 1.0 | intergenic | AATTAATTTTTTATTAAATTTTTTAATTTTAAAAT**TCTATT** |
| AE009951.2 | p_0629 | 1915115 | p_0629 | ATTTAATAA | 3 | - | 1.0 | intergenic | ATTATAACATTTTATAATTAAAAAAATTAATAAAA**TAGAAT** |
| AE009951.2 | p_0630 | 1918137 | p_0630 | TTATATAAT | 5 | + | 1.0 | intergenic | CTTCCCATTAACTTTTCCACCTCCCCCTGAAAAAT**TTTGAA** |
| AE009951.2 | p_0631 | 1917964 | p_0631 | TAGAAAAAA | 3 | - | 0.21 | intergenic | TATCATAATTTATAAATTAATTATAAAAAATAAAA**ATTAAG** |
| AE009951.2 | p_0632 | 1920765 | p_0632 | TTGTATAAT | 4 | + | 0.94 | intergenic | TACTGAGATAATGCAAAAAGAATTTGTTATAAAAA**TTTTAC** |
| AE009951.2 | p_0633 | 1920674 | p_0633 | TTATATAAT | 5 | - | 0.98 | intergenic | TTCTATTTTTGTAAAATTTTTATAACAAATTCTTT**TTGCAT** |
| AE009951.2 | p_0634 | 1921411 | p_0634 | TAAAATAAA | 3 | + | 1.0 | intergenic | TAGATTATTAGATAATTTTTAAAAGTTAATTCATA**TAAAAT** |
| AE009951.2 | p_0635 | 1928711 | p_0635 | ATTATATTT | 3 | + | 1.0 | intergenic | AATTAATTAAAAATCTAATATAAAATTTTTTTTAA**AAAACT** |
| AE009951.2 | p_0636 | 1928619 | p_0636 | TGCTAAAAT | 3 | - | 1.0 | intergenic | TTTAAGTCAAGTTTTTTAAAAAAAATTTTATATTA**GATTTT** |
| AE009951.2 | p_0637 | 1938538 | p_0637 | TATTATAAT | 5 | + | 1.0 | intergenic | TTTTTCTATACTTTTCTCACCTTCATAATTAATCT**TTGCTT** |
| AE009951.2 | p_0638 | 1938510 | p_0638 | TATTATAAT | 5 | - | 1.0 | intergenic | AAAATCACCCCTATTTTTTAAAGTTTGTTTTTATA**TTAATT** |
| AE009951.2 | p_0639 | 1938387 | p_0639 | TGTTATAAT | 5 | - | 0.89 | intergenic | ATAGAGAGTATCAAAGCTTATAAAAAAATATAATAT**TTCTTA** |
| AE009951.2 | p_0640 | 1956834 | p_0640 | ATTTAAATA | 3 | + | 1.0 | intergenic | CCTAATTTTAATTATTTTTCCAATTAATATACATT**AATTTT** |
| AE009951.2 | p_0641 | 1956749 | p_0641 | GGATATAAT | 4 | - | 1.0 | intergenic | ATTTAATAAAAATATAAAAATTAATGTATATTAAT**TGGAAA** |
| AE009951.2 | p_0642 | 1960993 | p_0642 | TGTTAAAAT | 4 | + | 1.0 | intergenic | ATTTTTATTTTTTATAAATTCAATAAATTTTTAAT**CTTTAA** |
| AE009951.2 | p_0643 | 1960878 | p_0643 | TAGAATAAA | 3 | - | 1.0 | intergenic | TTTATTGAATTTATAAAAAATAAAAATAAGGGTTG**ACTTAT** |
| AE009951.2 | p_0644 | 1961761 | p_0644 | TGATATAAT | 5 | + | 1.0 | intergenic | ATATTAAAGATTAAGAAATAAAATTTTATTTCTTA**ATCTTT** |
| AE009951.2 | p_0645 | 1967160 | p_0645 | TGTTAAAAT | 4 | + | 1.0 | intergenic | TAAGTCAAGATATTTTTTTAAGAAACTATTATTTT**TATAAA** |
| AE009951.2 | p_0646 | 1967063 | p_0646 | TATTATAAA | 3 | - | 0.99 | intergenic | AAATTTTATAAAAATAATAGTTTCTTAAAAAAATA**TCTTGA** |
| AE009951.2 | p_0647 | 1981996 | p_0647 | TGTTAAAAT | 4 | - | 1.0 | intergenic | TATAAAAATATTTTTCTAATAAAAATAAATTTAAT**TAACAA** |
| AE009951.2 | p_0648 | 1991655 | p_0648 | ATTTTTATA | 3 | + | 0.14 | intergenic | TATTTATGTTTTTACCATAAATATTATAACCACAA**ATAGAG** |
| AE009951.2 | p_0649 | 1991801 | p_0649 | TTAAAATTT | 3 | + | 1.0 | intergenic | TAGTGTCATTTTTAATTTAAAAAAGTATATTACAT**AAATTT** |
| AE009951.2 | p_0650 | 1991731 | p_0650 | AATTAAAAA | 3 | - | 0.97 | intergenic | ATTATATTAAATTTTAAGGAAAATTTCCAAAAAAT**TTATGT** |
| AE009951.2 | p_0651 | 1991599 | p_0651 | GGTTATAAT | 4 | - | 1.0 | intergenic | AACAGCCCCTCTTTTTTTAGTGTATAAAAATAGAA**TTGATT** |
| AE009951.2 | p_0652 | 1996095 | p_0652 | TGAAAATAT | 4 | - | 1.0 | intergenic | GAAATATGGATATATTAGAAATTAAAAGAGAATTT**TTAGAA** |
| AE009951.2 | p_0653 | 1997329 | p_0653 | TGCTATAAT | 5 | - | 0.99 | intergenic | AGTAATTAATAAGGATTAATTTCTAAAAAATAGGA**GTTAAT** |
| AE009951.2 | p_0654 | 1998147 | p_0654 | TATTATAAT | 5 | - | 1.0 | intergenic | GTAAAAAAATTAATTTTCAAGACTGTAAAAATTTA**TATGTT** |
| AE009951.2 | p_0655 | 1997929 | p_0655 | TGATAACAA | 2 | - | 1.0 | intergenic | AGTAGCAATATAAAAATAGAAATGCCTAAAAATAG**ATGGAT** |
| AE009951.2 | p_0656 | 2002927 | p_0656 | AAAATAAGG | 2 | - | 0.08 | intergenic | TACATTCGTAATTCGTTTATTTTTATTTATTAATA**ATATAT** |
| AE009951.2 | p_0657 | 2008146 | p_0657 | TTATATAAT | 5 | + | 1.0 | intergenic | TATAAAATATTTATCTATACAACCTCCTTTCATAG**TTTATT** |
| AE009951.2 | p_0658 | 2008058 | p_0658 | TTATATATT | 3 | - | 0.2 | intergenic | AAATAATAAATAAAATAAACTATGAAAGGAGGTTG**TATAGA** |
| AE009951.2 | p_0659 | 2009637 | p_0659 | CCTATCTTT | 1 | + | 1.0 | intergenic | AAAATTCTTATTTAAATTTTTTTAAAAAAGTTCTT**GACTTT** |
| AE009951.2 | p_0660 | 2009803 | p_0660 | TGGTAAAAT | 3 | + | 0.97 | intergenic | TTGAGAAATATTTAATTACTTGTTTATTAAAAAAT**TTCACA** |
| AE009951.2 | p_0661 | 2010751 | p_0661 | TTCTAAAAA | 3 | + | 0.98 | intergenic | AAGGACATAAAAGGACAAATGTATGAAAATTTGAT**TTTTTA** |
| AE009951.2 | p_0662 | 2022520 | p_0662 | TAGTATAAT | 4 | + | 0.98 | intergenic | CCATAAAATCCTAAAATTTAAACATTTTCTCTCAC**TTTACA** |
| AE009951.2 | p_0663 | 2022436 | p_0663 | TGGTAAAAT | 3 | - | 1.0 | intergenic | CTATTTATATATAGTTTTGTAAAGTGAGAGAAAAT**GTTTAA** |
| AE009951.2 | p_0664 | 2043726 | p_0664 | AAGTATTAT | 3 | - | 0.92 | intergenic | TCTCTCTTTTTTTATTTAAATTTTCTTTTAATTTA**TTGGAA** |
| AE009951.2 | p_0665 | 2051902 | p_0665 | TTGTATAAT | 4 | + | 0.94 | intergenic | TTATTCTATTAATCAAACATTTCAATATTCTTTGC**TTGACA** |
| AE009951.2 | p_0666 | 2051791 | p_0666 | TGATATATA | 4 | - | 1.0 | intergenic | AGAATATTGAAATGTTTGATTAATAGAATAAATAA**AATCAA** |
| AE009951.2 | p_0667 | 2051649 | p_0667 | TGATATAAT | 5 | - | 0.52 | intergenic | TAAGTAAGAGTAGTTCATTAGAAAATAGTACTCTC**TTTCTA** |
| AE009951.2 | p_0668 | 2054470 | p_0668 | TTATATAAT | 5 | + | 0.95 | intergenic | AGCAAGAGAATAAAATCTCTTGCTATTTTTTATTA**ACGAAT** |
| AE009951.2 | p_0669 | 2058662 | p_0669 | AAATATAAT | 5 | + | 1.0 | intergenic | ATTTAAAAAAAATTTTTTTAAATATAAAAATAAAA**ACGGAA** |
| AE009951.2 | p_0670 | 2070756 | p_0670 | TTTTTTATA | 3 | + | 0.97 | intergenic | GAAGCAAAAGTTAATCCTTTTAATTTTTGCTTAAT**TTTTTA** |
| AE009951.2 | p_0671 | 2077017 | p_0671 | ATTTAATAG | 3 | + | 1.0 | intergenic | TAAAAAAACTGTACATGAACAGTTTTATAAAAATG**CTTGTT** |
| AE009951.2 | p_0672 | 2080679 | p_0672 | TATTATAAT | 5 | + | 0.95 | intergenic | TGTACTATGACAGTTTTTAGTATAAATAATTTATT**TATAAA** |
| AE009951.2 | p_0673 | 2080623 | p_0673 | AATTATTTA | 3 | - | 0.07 | intergenic | CTAGTATATTTGATATTTTTATATTATAATATTAA**CATAAA** |
| AE009951.2 | p_0674 | 2088833 | p_0674 | ATTATAAAA | 3 | - | 1.0 | intergenic | AGTAATCTTCTTTTTTATTAAAAAATAAAAATAAT**TAAATT** |
| AE009951.2 | p_0675 | 2088571 | p_0675 | TTTTATAAA | 3 | - | 0.99 | intergenic | CCTTCTTAAATGTAAAAATAATTTTATAGTTATTT**TTTATA** |

| | | | | | | | | | |
|---|---|---|---|---|---|---|---|---|---|
| AE009951.2 | p_0679 | 2095949 | p_0679 | TGTTATAAT | 5 | + | 1.0 | intergenic | TTTTTTATCCCAAATGTTTAAAAATTTTGACATTC**TTTTAC** |
| AE009951.2 | p_0680 | 2095870 | p_0680 | ATAAAAAAA | 3 | - | 0.99 | intergenic | TTATAACATTTTTAAGAATTTAGTAAAAGAATGTC**AAAATT** |
| AE009951.2 | p_0681 | 2100430 | p_0681 | TACAATTTA | 3 | - | 0.63 | intergenic | AAAAACAACTTCTGCTTTATAATATGAATAGCTTA**CTAAAT** |
| AE009951.2 | p_0682 | 2101712 | p_0682 | TTTTTTTTT | 3 | - | 1.0 | intergenic | ATAAACTGTCTTTTTTATTGATACAAAAATATTAA**AAAAAT** |
| AE009951.2 | p_0683 | 2105094 | p_0683 | TGGTATAAT | 5 | - | 1.0 | intergenic | TTTGAAAAGGTATGACTAAAAGATAAAAAAATCAA**ATTTCT** |
| AE009951.2 | p_0684 | 2106357 | p_0684 | TGGTAAAAA | 3 | + | 1.0 | intergenic | TTTAATTTATAAAAAAATCAAGAATAAAAATCTATA**AAATTC** |
| AE009951.2 | p_0685 | 2106270 | p_0685 | TGATAAAAT | 4 | - | 1.0 | intergenic | TTTAATAAAAAAAAGAATTTTATAGATTTTTATTC**TTGATT** |
| AE009951.2 | p_0686 | 2111702 | p_0686 | TGATATAAT | 5 | + | 0.67 | intergenic | GTGTCCAAGATTATGGGTGCAGTTCAATTATATAT**TGGGAC** |
| AE009951.2 | p_0687 | 2116115 | p_0687 | TGTTTAAAA | 3 | + | 0.7 | intergenic | ATGTTAATATAATATTTTTGATATGTTAAGATTAG**GTATTT** |
| AE009951.2 | p_0688 | 2118578 | p_0688 | TAGAATATA | 3 | - | 1.0 | intergenic | AAAGAAGGTTGAAATTTTCAATCTTCTTTTTATTT**GTTATA** |
| AE009951.2 | p_0689 | 2129971 | p_0689 | TCTTATAAT | 4 | - | 1.0 | intergenic | TTGTATATATTACAATTGTATTTTTTAAATTTTTA**TTGAAA** |
| AE009951.2 | p_0690 | 2131612 | p_0690 | TTAAAAAAA | 3 | - | 1.0 | intergenic | TTTTTTCTGTAATAATTAATTTTAAAATTTTTTGT**TAGTTT** |
| AE009951.2 | p_0691 | 2143547 | p_0691 | TGATATAAT | 5 | - | 1.0 | intergenic | AGCATTAAAAGAAAATAAAAAAATAAAAATAATAG**ATACTT** |
| AE009951.2 | p_0692 | 2147948 | p_0692 | CCTTTAAAA | 3 | + | 1.0 | intergenic | TTTTCCAAAGTTTCCTCTATCTGTTTTCTTTGTTT**CAACAA** |
| AE009951.2 | p_0693 | 2148028 | p_0693 | TAAAAAATA | 3 | - | 1.0 | intergenic | TAACTTCTATATTTATTATAATATTTTATGTTGAA**AAATCA** |
| AE009951.2 | p_0694 | 2151932 | p_0694 | TACAATATT | 3 | + | 0.73 | intergenic | AAACTTTAAAAATAAAAGTCAATAAAATATTTGAC**ATTGTC** |
| AE009951.2 | p_0695 | 2151816 | p_0695 | AAAAAATTA | 3 | - | 1.0 | intergenic | TTTATTGACTTTTATTTTTAAAGTTTTGATAATAA**TGTTTT** |
| AE009951.2 | p_0696 | 2159636 | p_0696 | TTTTATAAT | 5 | + | 0.69 | intergenic | ACTATATCACACTCTAAATTTTACTTCATTTATAA**GTTTGC** |
| AE009951.2 | p_0697 | 2169527 | p_0697 | TTCGCTAAA | 2 | - | 1.0 | intergenic | AAGAATTCGCATCTAAAAAACTCTAAGCAATAAAT**TGCTAA** |
