## Supplementary_README for "Integration of Bioinformatics and Machine Learning to characterize *Fusobacterium nucleatum*’s pathogenicity"

SUPPLEMENTARY MATERIALS README
==============================

Integration of Bioinformatics and Machine Learning to Characterize 
Fusobacterium nucleatum's Pathogenicity

Authors: Zihan Tian, Pietro Lio'

---
SUPPLEMENTARY Zip
--------------------
Unzip the supplementary.zip; below is the contents.

SUPPLEMENTARY TABLES
--------------------

Table S1a-c: BLAST analysis results for genes in predicted pathogenicity 
islands PAI1, PAI2, and PAI3.

Table S2: Promoter predictions for F. nucleatum ATCC 25586 genome using 
PePPER. Contains predicted promoters with confidence scores 
(0.0-1.0), -10 box sequences, and promoter sequences.

Table S3: STRING protein-protein interaction network data for PAI2 proteins.

Table S4: AlphaFold-Multimer structure predictions and TMHMM transmembrane 
helix predictions for PAI2 proteins. Includes pTM and ipTM confidence scores.

Table S5: Co-expression network edges predicted by Bayesian network learning 
(bnlearn) from F. nucleatum transcriptomic data (n=18 samples) that is significant.

Table S6: Overlap between bnlearn co-expression predictions and STRING 
functional associations.

Table S7: Flux balance analysis (FBA) results showing predicted growth rates 
under different iron availability conditions.

---

SUPPLEMENTARY DATA
------------------

Data S1: Genome-scale metabolic model of F. nucleatum ATCC 25586 in SBML 
format, generated using CarveMe.

Data S2: F. nucleatum ATCC 25586 genome sequence in FASTA format 
(NCBI accession: AE009951).

Data S3: Genome annotation in GenBank format.

Data S4: Python analysis pipeline (Jupyter notebook) containing code for:
- Pathogenicity island prediction
- CAI analysis
- Metabolic modeling

Data S5: R script for Bayesian network learning (bnlearn) co-expression 
analysis. With readme for code.

---

SOFTWARE VERSIONS
-----------------

Python 3.9
- COBRApy 0.26.0
- Biopython 1.79
- pandas 1.4.0
- numpy 1.22.0

R 4.1.0
- bnlearn 4.7

External tools:
- CarveMe 1.5.1
- IslandViewer 4
- Alien Hunter 1.7
- PePPER (web server)
- STRING v11.5
- AlphaFold-Multimer v2.3

---

FILE NAMING NOTE
----------------

Input files referenced in the analysis code use their original names from 
public databases. The supplementary files have been renamed for clarity:

Original Name -> Supplementary Name
EX:
25586_NCBI.gbk -> Supplementary_data_S3_ geneBank.gbk

---

DATA AVAILABILITY
-----------------

F. nucleatum ATCC 25586 genome: NCBI accession AE009951
Transcriptomic data: GEO accession [GSE161360]

For questions, contact: []
